# Machine learning links T cell function and spatial localization to neoadjuvant immunotherapy and clinical outcome in pancreatic cancer

**DOI:** 10.1101/2023.10.20.563335

**Authors:** Katie E. Blise, Shamilene Sivagnanam, Courtney B. Betts, Konjit Betre, Nell Kirchberger, Benjamin Tate, Emma E. Furth, Andressa Dias Costa, Jonathan A. Nowak, Brian M. Wolpin, Robert H. Vonderheide, Jeremy Goecks, Lisa M. Coussens, Katelyn T. Byrne

**Affiliations:** Department of Biomedical Engineering, Oregon Health & Science University, Portland, OR USA; The Knight Cancer Institute, Oregon Health & Science University, Portland, OR USA; Department of Cell, Developmental & Cancer Biology, Oregon Health & Science University, Portland, OR USA; Current affiliation: Akoya Biosciences, 100 Campus Drive, 6th Floor, Marlborough, MA USA; Immune Monitoring and Cancer Omics Services, Oregon Health & Science University, Portland, OR USA; Department of Pathology and Laboratory Medicine, Hospital of the University of Pennsylvania, Philadelphia, PA USA; Abramson Cancer Center, Perelman School of Medicine, University of Pennsylvania, Philadelphia, PA USA; Department of Medical Oncology, Dana-Farber Cancer Institute, Harvard Medical School, Boston, MA USA; Department of Pathology, Brigham and Women’s Hospital and Harvard Medical School, Boston, MA USA; Parker Institute for Cancer Immunotherapy, Perelman School of Medicine, University of Pennsylvania, Philadelphia, PA USA; Current affiliation: Department of Machine Learning, H. Lee Moffitt Cancer Center, Tampa, FL USA; Current affiliation: Department of Biostatistics and Bioinformatics, H. Lee Moffitt Cancer Center, Tampa, FL USA

**Keywords:** Tumor microenvironment, machine learning, T cell, pancreatic ductal adenocarcinoma, spatial proteomics

## Abstract

Tumor molecular datasets are becoming increasingly complex, making it nearly impossible for humans alone to effectively analyze them. Here, we demonstrate the power of using machine learning to analyze a single-cell, spatial, and highly multiplexed proteomic dataset from human pancreatic cancer and reveal underlying biological mechanisms that may contribute to clinical outcome. A novel multiplex immunohistochemistry antibody panel was used to audit T cell functionality and spatial localization in resected tumors from treatment-naive patients with localized pancreatic ductal adenocarcinoma (PDAC) compared to a second cohort of patients treated with neoadjuvant agonistic CD40 (αCD40) monoclonal antibody therapy. In total, nearly 2.5 million cells from 306 tissue regions collected from 29 patients across both treatment cohorts were assayed, and more than 1,000 tumor microenvironment (TME) features were quantified. We then trained machine learning models to accurately predict αCD40 treatment status and disease-free survival (DFS) following αCD40 therapy based upon TME features. Through downstream interpretation of the machine learning models’ predictions, we found αCD40 therapy to reduce canonical aspects of T cell exhaustion within the TME, as compared to treatment-naive TMEs. Using automated clustering approaches, we found improved DFS following αCD40 therapy to correlate with the increased presence of CD44^+^ CD4^+^ Th1 cells located specifically within cellular spatial neighborhoods characterized by increased T cell proliferation, antigen-experience, and cytotoxicity in immune aggregates. Overall, our results demonstrate the utility of machine learning in molecular cancer immunology applications, highlight the impact of αCD40 therapy on T cells within the TME, and identify potential candidate biomarkers of DFS for αCD40-treated patients with PDAC.

## INTRODUCTION

Pancreatic ductal adenocarcinoma (PDAC) is one of the most aggressive treatment-refractory cancers with a median overall survival rate of just months, thus there is a critical need for new treatment strategies for this disease (1-3). Recent reports reveal that immunological responses in PDAC can be temporarily induced via approaches that promote priming of T cell responses against PDAC, such as occurs following agonistic CD40 (αCD40) monoclonal antibodies (4) and peptide vaccination (5) strategies. We and others have previously reported that αCD40 binds to CD40 on dendritic cells (DCs), thereby licensing DCs to subsequently enhance T cell activation and bolster anti-tumor immunity (6-8). However, little is known regarding selective impact of αCD40 therapy on specific T cell states within the TME and to what degree this therapy sustains T cell functionality in situ, or instead promotes T cell dysfunction that could in turn limit potential use of αCD40 combination therapies in the clinical setting.

Upon antigen stimulation, T cells upregulate CD44 and exist along a spectrum of diverse differentiation states with varying functionalities (9-11). On one end of the spectrum, T cells possess stem-cell-like plasticity, accompanied by memory, proliferative and cytotoxic capabilities, and are identified by expression of T-BET, and/or TCF-1 (12, 13). On the other end of the spectrum, T cells are exhausted and/or dysfunctional and canonically express TOX1 and/or EOMES (14, 15). Along the spectrum, expressed in varying combinations, are T cells expressing immune checkpoint molecules such as PD-1, LAG- 3 and TIM3, with increased numbers of immune checkpoint molecules correlating with more exhausted T cells (16). These partially exhausted T cells are susceptible to reinvigoration by immune checkpoint blockade (ICB) and regain the ability to proliferate and produce effector cytokines after PD-1 blockade, for example (17, 18). However, terminally differentiated T cells expressing TOX1 or EOMES are resistant to rescue by ICB and cannot proliferate or exert cytotoxic activity (19). Flow cytometric analyses of T cells in preclinical tumor models or tissues following viral infection have elucidated notable T cell states associated with priming, proliferation, cytotoxic capability, or functional exhaustion; however, characterizations of effector versus exhausted T cell phenotypes from tumors in patients are scant. Moreover, with the advent of single-cell sequencing approaches, the diversity of T cell subset possibilities within tumors is seemingly endless and in need of clarification (20, 21). We sought to clarify T cell subset characteristics within the PDAC TME, and to identify those subsets associated with response to therapeutic αCD40 therapy. Recognizing that both the cellular composition and spatial organization of cells, including T cells, is a critical metric associated with therapeutic response and clinical outcome (22-32), we investigated and report on the impact of αCD40 therapy on the complex spatial contexture of T cells within the PDAC TME and association with survival.

We previously developed a multiplex immunohistochemistry (mIHC) single-cell spatial proteomics imaging platform to interrogate leukocyte heterogeneity and spatial landscape within the TMEs of various cancer types (33-35). Following a cyclical staining protocol, the mIHC platform iteratively utilizes up to 30 antibodies to stain protein biomarkers on a single tissue specimen, thus preserving spatial context of the TME. The resulting data provide single-cell resolution maps, which can be quantified by several metrics including cellular density, cell-cell spatial interactions, and cellular neighborhoods. While it is possible to quantify and correlate each of these TME features individually with clinical outcome, this approach fails to combine and weight features together to capture biological complexities of the TME. Machine learning can be used to address TME complexity, as it is a form of artificial intelligence whereby computational models are trained to identify which combinations of data features can predict a given output. Models can then be used to make accurate predictions from new data and interrogated to understand which combinations of features are most important in making the predictions. Machine learning is becoming widely utilized in precision oncology to decipher patterns in large data sets resulting from deep interrogations (36-39).

Here, we leveraged machine learning to elucidate the frequency of various T cell states in PDAC and investigated the impact of αCD40 therapy on those states. We first designed a novel mIHC antibody panel to deeply audit T cell functionality and spatial organization in two PDAC cohorts, one of which was treatment-naive and the other of which received neoadjuvant αCD40 therapy. Using this mIHC panel, we generated a dataset of nearly 2.5 million cells with spatially resolved single-cell phenotypic and functional measurements. Interrogation of this dataset presented a unique opportunity to elucidate: 1) the types of T cells present at baseline in a treatment-refractory disease and 2) to what degree αCD40 therapy sustains T cell functionality in situ, or instead promotes T cell dysfunction that could in turn limit potential use of αCD40 or other T cell priming therapies in the clinical setting. Given the vast amount of spatially resolved data and complexity of T cell function in the TME, we leveraged machine learning approaches to discern new biological insights regarding T cells in the pancreatic TME and their association with clinical outcome for pancreatic cancer patients.

## MATERIALS AND METHODS

### Tissue Acquisition

Human PDAC tissue specimens from cohort 1 were obtained from patients with approval from the Oregon Pancreas Tissue Registry under Oregon Health & Science University IRB protocol #3609 and Dana Farber Harvard Cancer Center protocols #03-189 and #12- 013 (**Fig. 1A**). Specimens from cohort 2 were obtained from patients treated with neoadjuvant selicrelumab with approval under the IRBs of four sites across the United States involved in an open-label phase I clinical trial (Cancer Immunotherapy Trials Network CITN11-01; NCT02588443; **Fig. 1A**), including 8 patients with neoadjuvant selicrelumab alone and 3 patients with neoadjuvant selicrelumab combined with gemcitabine and nab-paclitaxel. For the purposes of this paper, this cohort has been combined as the αCD40-treated cohort given previous analyses of the TME (4). All studies were conducted in accordance with the Declaration of Helsinki and written informed consent was obtained.

**Figure 1.**
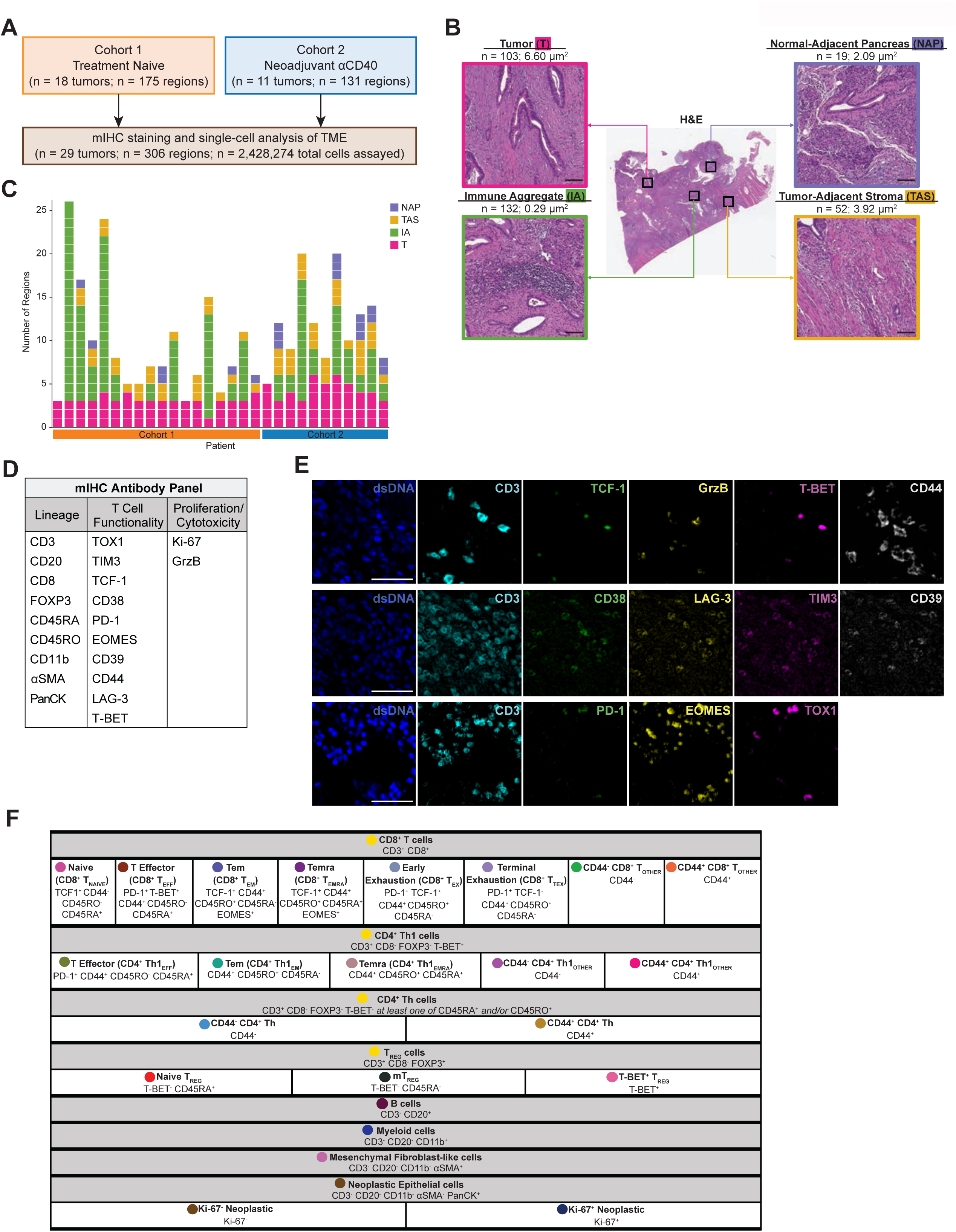
Designing a novel mIHC antibody panel to deeply phenotype T cells within the PDAC TME. **A.** Overview of two PDAC cohorts assayed via mIHC. **B.** Representative PDAC tissue resection stained with H&E (middle) showing four histopathologic sites annotated. Total number of regions assayed and average tissue area per histopathologic site are listed. Scale bars = 100 µm. **C.** Number of regions assayed per patient. Each box represents one tissue region and is colored according to its histopathologic site. **D.** 21-marker mIHC antibody panel used to assay tissue regions. **E.** Representative pseudo-colored mIHC images showing T cell functionality markers with CD3 expression. Scale bars = 50 µm. **F.** Cell phenotyping strategy by hierarchical gating of lineage markers and functional markers. Circles indicate colors associated with each cell state in the following figures.

### Multiplex Immunohistochemistry Image Acquisition and Analysis

Formalin-fixed, paraffin-embedded (FFPE) surgical tissue samples were sectioned and assessed using hematoxylin and eosin (H&E), as well as chromogen based mIHC. Using pathologist annotations overlaid from the H&E-stained slides, sections were assessed with regard to histopathologic regions of interest (ROI; **Fig. 1B**) annotated as tumor (T), immune aggregate (IA), tumor-adjacent stroma (TAS), or normal-adjacent pancreas (NAP), as defined by the pathologist (35). Multiplex staining was performed on 5 µm sections, as previously described, and each stained image was scanned at 20x magnification on an Aperio AT2 scanner (Leica Biosystems) (35). The antibody panel was designed to delineate epithelial cells, mesenchymal cells, B cells, myeloid cells, and 18 T cell subpopulations, spanning T cell activation and exhausted states. Human tonsil and spleen were included in all rounds of mIHC as staining controls. Image processing was performed using previously described methods (35). Each region was registered to the final hematoxylin using Matlab Computer Vision Toolbox (The Mathworks, Inc., Natick, MA), color deconvolution and watershed-based nuclei segmentation was performed using ImageJ, and single cell mean intensity for each stain was quantified using Cell Profiler (40). Single marker positivity thresholds were set using FCS Express Image Cytometry RUO (De Novo Software, Glendale, CA) to visually validate protein biomarker expression overlaid on signal extracted images. Single cell classification was performed using R Statistical Software based on filtering exclusive populations in a defined hierarchy.

### Machine Learning Classifiers and Feature Importance Analyses

Elastic net (EN) classifier models were built to predict: 1) treatment status and 2) disease-free survival (DFS), using scikit-learn’s LogisticRegression function (41). Predictions were made on a region basis to maximize sample size and model robustness. Separate models were created for each histopathologic site in order to: 1) compare performance of models derived from different histopathologic sites; 2) identify where therapy was potentially exerting greatest impact; and 3) mitigate broad variation in average tissue area from each histopathologic site (**Fig. 1B**). A leave-one-patient-out cross-validation approach was used to split the train and test sets. Thus, within each cross-validation loop, a new EN model was created and trained on all regions except those from one patient, and testing was then performed on regions from the patient withheld from training. This process was repeated until all patients were cycled through the test set. This approach prevents data leakage by ensuring regions from the same patient were not in both the train and test sets for one model, thus preventing the EN models from learning patient-specific features, which often artificially increases model accuracy. Test set predictions were aggregated across all cross-validation loops to construct one final confusion matrix, from which performance of the models was assessed by calculating accuracy, F1 score, and area under the receiver operating characteristic curve (AUC). These metrics address both precision and recall (F1 score), in addition to the true positive rate and false positive rate (AUC) – these are often used to assess performance of classifier models. Model overfitting was mitigated by using the same model hyperparameters across cross-validation loops. The penalty term was set to “elasticnet,” and the “l1_ratio” hyperparameter was set to 0.5, representing an equal balance of the lasso model and ridge model effects. All features were log10+1 normalized and scaled using a minmax [0,1] scaler in order to equally compare features spanning different orders of magnitude and improve model interpretability. To further prevent data leakage, in each cross-validation loop, the scaler was fit to the train set and then applied to the train set and subsequently the test set. Test feature outliers were clipped to [0,1] following this normalization. The train set was balanced within each cross-validation loop using Synthetic Minority Over-sampling Technique (SMOTE) to up-sample the minority class to equal the majority class (42). Feature importance analyses were conducted by computing Shapley Additive exPlanations (SHAP) values for each model (43). SHAP values enable the interpretation of which combinations of features contribute to the overall model predictions.

### Cell-Cell Spatial Interaction Quantification

Cell-cell interactions were defined as cells separated by a distance of ≤ 20 µm, according to centroid (x,y) coordinates of any two cells, enriching for cells that were directly adjacent as previously reported (44-46). Total cell-cell interactions were normalized by dividing summed densities of cell states involved in the interactions to avoid skewing by cell states present in high abundances.

### Recurrent Cellular Neighborhood Analysis

Recurrent cellular neighborhoods were quantified to assess spatial organization of tissues. A neighborhood was created for every cell by counting all cells within a 60 µm radius of each seed cell’s center, as inferred from previous studies (47-50). Using proportions of cells comprising the neighborhoods as features, neighborhoods were grouped using K-means clustering. The elbow method was used to determine the number of clusters, resulting in groupings of spatial neighborhoods that were similar in cellular composition that could be found across all regions of interest in the analysis.

### Statistics

Mann-Whitney U tests were used to determine statistically significant differences in top TME features between treatment cohorts or DFS groups. The Benjamini-Hochberg correction was used to account for multiple hypothesis testing for each analysis. *P*-values less than 0.05 were considered statistically significant. Statistical calculations were performed with the Scipy and statsmodels packages using Python software (51, 52).

## DATA AVAILABILITY

mIHC data used for this study is available for download on Zenodo at https://doi.org/10.5281/zenodo.8357193.

## CODE AVAILABILITY

The code used to generate all computational results of this research was created using Python version 3.9.4 and is available at https://github.com/kblise/PDAC_mIHC_paper.

## RESULTS

### Designing a novel mIHC antibody panel to deeply phenotype T cells within the PDAC TME

29 PDAC tumors were surgically resected from patients across two treatment cohorts (**Fig. 1A**). Tumors from cohort 1 were treatment-naive upon resection and were previously evaluated for immune contexture as a subset of a larger study (n=104 tumors) designed to broadly evaluate leukocyte composition in treatment-naive PDAC (35). The 18 treatment-naive tumors evaluated in the current study were selected as a representative subset of the larger PDAC cohort based on cellular subsets not being statistically different from the full cohort in terms of leukocyte densities or patient survival durations (**Supplementary Table S1**). Thus, PDAC specimens from cohort 1 served as a representative baseline comparison to the 11 specimens from cohort 2, which reflected patients who had received neoadjuvant αCD40 therapy alone (n=8) or in combination with gemcitabine and nab-paclitaxel (n=3) prior to resection (4). Three to 26 ROIs per PDAC resection were selected by a pathologist and quantitatively assayed by mIHC, with each region annotated as one of four histopathologic sites within the resected samples: tumor (T), immune aggregate (IA), tumor-adjacent stroma (TAS), or normal-adjacent pancreas (NAP) (**Fig. 1B**) (35). Breakdown of region types assayed per patient are shown (**Fig. 1C**).

In total, nearly 2.5 million cells were assayed across 306 tissue regions by our novel 21- antibody mIHC panel (**Fig. 1D and E, Supplementary Table S2**, **Supplementary Fig. S1A**). Given our goal of revealing T cell states present in the PDAC TME at baseline, and the impact of αCD40 therapy on those T cells, the majority of antibodies included in the mIHC panel were chosen for their ability to elucidate differentiation, exhaustion, proliferative and cytotoxic status of various T cell lineage subtypes as defined by prior literature (14, 53-57). First, 423,317 T cells were identified by CD3 expression and then subsequently stratified by CD8α expression. CD8^+^ T cells were further classified as one of six cell states (TNAIVE, TEFF, TEM, TEMRA, TEX, or TTEX) (**Fig. 1F**). Due to biomarker selection and positional restrictions within the cyclic multiplex panel, we did not include a CD4 antibody. However, the majority (72%) of CD3^+^ CD8^-^ T cells were CD4^+^ as determined in a testing panel using a subset of the data (**Supplementary Fig. S1B**); we therefore refer to CD3^+^ CD8^-^ T cells as CD4^+^ T cells herein, although it is possible other minor lineages may be represented (20). Based on this schema, CD3^+^ CD8^-^ T cells were further evaluated; CD4^+^ Th1 cells were defined by T-BET^+^ expression, and further classified as one of three cell states (TEFF, TEM, or TEMRA) (**Fig. 1F**). We originally anticipated the majority of CD8^+^ T cells and CD4^+^ Th1 cells would fall into one of the clearly defined states (e.g., TNAIVE, TEFF, TEM, TEMRA, TEX, TTEX); however, only 6% of CD8^+^ T or CD4^+^ Th1 cells were phenotyped as one of these states (**Supplementary Table S3**). To investigate the remaining 94% – labeled TOTHER (**Fig. 1F**) – we stratified cells based on expression of CD44, a canonical biomarker of prior cognate antigen experience. Furthermore, there was a population of T-BET^-^ CD4^+^ Th cells (non-Th1-specific T helper cells), which we stratified into two cell states based upon CD44 expression only (**Fig. 1F**). Finally, the remaining CD3^+^ CD8^-^ T cells were FOXP3^+^ TREG cells, and classified into three cell states (Naive TREG, mTREG, and T-BET^+^ TREG) (**Fig. 1F**). Despite low numbers of certain T cell states (**Supplementary Table S3**), we included all 18 T cell states in downstream analyses based upon calculations from previous single-cell studies (58), as well as the fact that these populations were manually gated and thus represent real phenotypes of T cells present in the PDAC TME. In addition, we utilized antibodies for 10 T cell functionality biomarkers to assess differentiation/exhaustion status on all T cells (TOX1, TIM3, TCF-1, CD38, PD-1, EOMES, CD39, CD44, LAG-3, and T-BET), as well as for proliferation (Ki-67) and cytotoxicity (granzyme B, GrzB) (**Fig. 1D, E**). Finally, the remaining CD3^-^ (non-T) cells were defined by a hierarchical gating strategy and classified as B cells, myeloid cells, mesenchymal fibroblast-like cells (also referred to as mesenchymal cells), or neoplastic epithelial cells (**Fig. 1F**, **Supplementary Fig. S1C**). Altogether, cells were phenotyped as one of 23 different cell lineages and states (**Fig. 1F**).

### Interrogating cell states and spatial interactions within the PDAC TME

Using single-cell spatial data collected from the mIHC assay, we used three approaches to quantify treatment-naive and αCD40-treated PDAC TMEs: 1) Cell state densities, to identify types of cells present (**Supplementary Table S3**); 2) T cell “functionality barcodes,” to reflect functional phenotypes of T cells by unique combinations of functional biomarker expression (**Supplementary Table S4**); and 3) Cell-cell spatial interactions, to reveal relative locations between cell types (**Supplementary Table S5**). Using these three metrics, we created a granular map of leukocyte infiltration, T cell functionality status, and cellular spatial orientation in the PDAC TME (**Fig. 2**).

**Figure 2.**
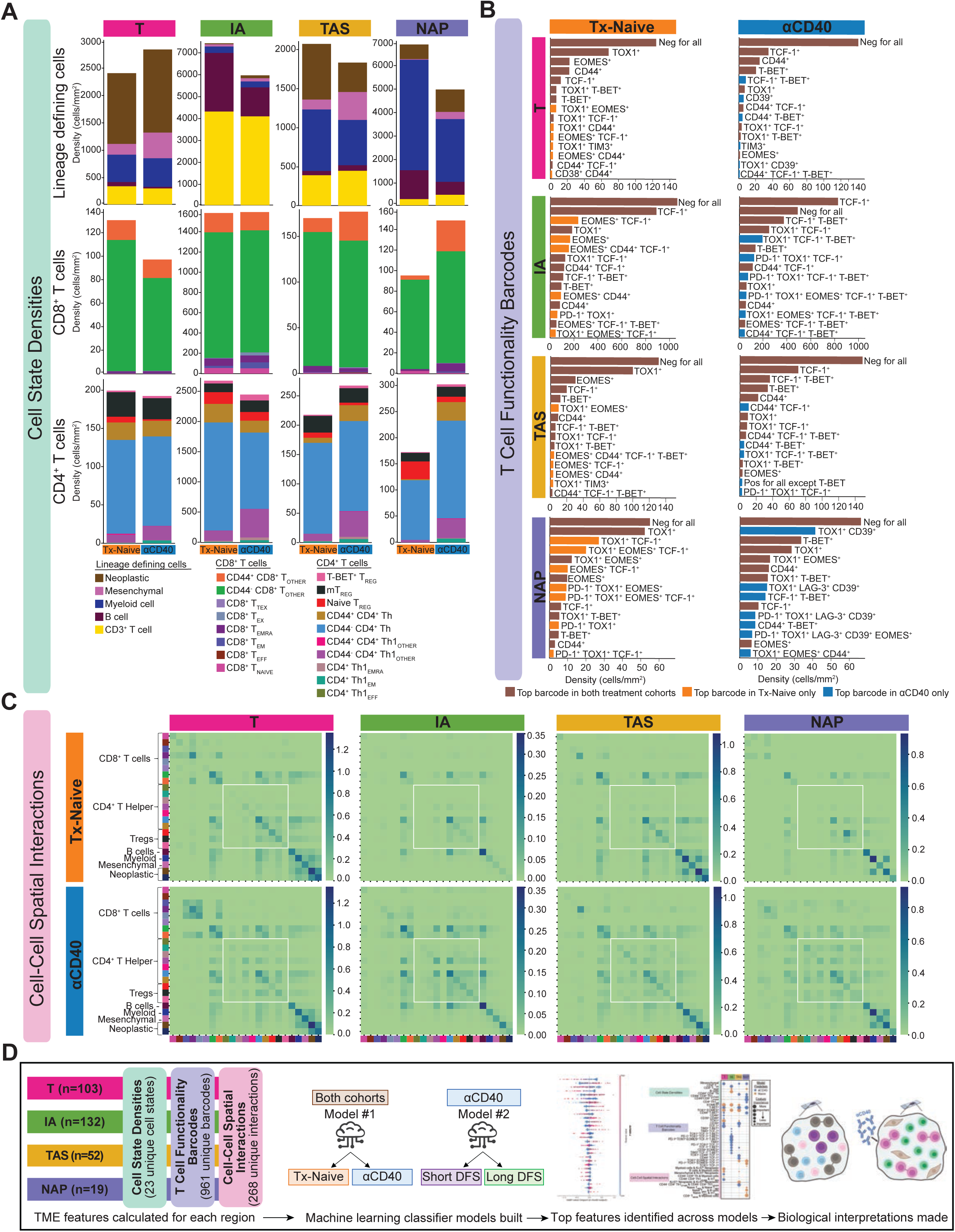
Interrogating cell states and spatial interactions within the PDAC TME. **A.** Stacked bar charts showing the average cell state densities for each treatment cohort and histopathologic site. Top row: lineage defining cells including neoplastic epithelial cells, mesenchymal fibroblast-like cells, myeloid cells, B cells, and CD3^+^ T cells; Middle row: CD8^+^ T cell states; Bottom row: CD4^+^ T cell states. Columns denote histopathologic site, and each plot is further broken into treatment cohort. **B.** Bar charts showing average densities of barcoded T cells for each treatment cohort and histopathologic site. Only the 15 most abundant barcodes are shown as measured by average density. Rows denote histopathologic site, and columns denote treatment cohort. Brown bars denote barcoded T cells that are in the top 15 most abundant barcodes in both cohorts. Orange bars denote barcoded T cells that are in the top 15 most abundant barcodes in the treatment-naive cohort only. Blue bars denote barcoded T cells that are in the top 15 most abundant barcodes in the αCD40-treated cohort only. **C.** Heatmaps showing average number of spatial interactions between two cell states for each treatment cohort and histopathologic site. Cell states are denoted by colors shown in Figure 1F. Interactions were normalized first by density of cells participating in the interaction and were then log10+1 transformed. Rows denote treatment cohort and columns denote histopathologic site. **D.** Overview schematic of analyses performed in this study. TME features were calculated for each tissue region. Two machine learning classifier models were built for each histopathologic site to predict treatment status and DFS. Feature importance analyses were performed to interpret biological meaning.

To identify the types and amounts of cell states present in the PDAC TME, we first quantified densities of each of the 23 cell states for each tissue region assayed by dividing raw counts of cells (**Supplementary Table S3**) by tissue area. Varying densities of leukocytes, mesenchymal fibroblast-like cells, and neoplastic epithelial cells were present in regions of each histopathologic annotation and each treatment cohort (**Fig. 2A**, **top**). In T regions, neoplastic epithelial cells were dominant. In IA regions, T cells and B cells were most prevalent, while in TAS regions, which encompassed the tumor borders, there was a mix of neoplastic cells, T cells, myeloid cells, and mesenchymal cells. Finally, myeloid cells were the dominant population in distal NAP regions. T cells were further divided into 18 T cell states defined by the mIHC gating strategy (**Fig. 1F**, **white boxes**). On average, CD4^+^ T cells were present in increased densities, nearly two-fold, as compared to CD8^+^ T cells (**Fig. 2A**, **middle**, **bottom rows**), regardless of histopathologic type or treatment cohort. However, average densities of the CD8^+^ and CD4^+^ T cell states often differed by histopathologic site and treatment cohort. These results demonstrate the importance of identifying spatial and histopathological information for interpreting how – and where – αCD40 therapy alters T cells in the PDAC TME, important information that could not be captured by flow cytometric methodologies.

To understand how T cell functionality differed based on histopathology and prior treatment, we assigned all T cell states a “Functionality Barcode,” as defined by binary positive or negative expression of unique combinations of 10 T cell functionality biomarkers (**Fig. 1D**, **middle column**). Among the 423,317 T cells present in the dataset, we identified expression of 961 unique barcodes (**Supplementary Table S4**). The top 15 most common barcodes by average density are shown for each histopathologic type and treatment cohort (**Fig. 2B**). Over half of the most common barcoded T cells were present in both treatment cohorts, regardless of histopathologic site, as indicated by brown bars (**Fig. 2B**). Of the barcodes present in ‘high’ densities only within the treatment-naive cohort (orange bars), the majority contained two functionality biomarkers; however, of the most abundant barcodes present in the αCD40 cohort only (blue bars, **Fig. 2B**), the majority contained three or more functionality biomarkers. This result supports the hypothesis that αCD40 therapy shifts T cell differentiation/functionality within the PDAC TME, as represented by an increase in the number of functionality biomarkers expressed. However, further elucidation of the specific patterns of biomarkers expressed by T cells is needed to identify the direction of this shift following immunotherapy.

Finally, to address spatial organization of cells in the PDAC TME, we leveraged the fact that mIHC preserves spatial context and quantified frequency of two cell states interacting, based on their cell centers being within 20 µm from each other (44-46). We identified 268 unique pairs of cell-cell interactions present in the datasets (**Supplementary Table S5**). On average, there were increased interactions between CD4^+^ T cells of various subpopulation states with other CD4^+^ T cells in the αCD40 cohort, regardless of histopathologic site (**Fig. 2C**, **white boxes**). Altogether, these results support the hypothesis that αCD40 drives an increase in CD4^+^ T cell density, functional capacity, and spatial proximity in the PDAC TME, as compared to treatment-naive PDAC TMEs.

Following these three TME single-cell quantification methods, all 2,428,274 cells present across the 306 regions were phenotyped as one of 23 cell states. In addition, all 423,317 T cells were assigned one of 961 T cell functionality barcodes. Finally, the immediate spatial neighbors of each cell were computed and binned into one of 268 types of pairwise cell-cell interactions. In total, 1,252 TME features were computed across the 306 regions, and each region was annotated as one of four histopathologic sites. Given the complexity and large amount of data, we leveraged machine learning and feature importance analyses to identify: 1) impact of αCD40 therapy on composition, functional capacity, and spatial organization of cells within the PDAC TME, and 2) the likely mechanism(s) we hypothesized underlying improved clinical outcome following αCD40 therapy. To do this, we trained EN classifier models to predict treatment status and DFS within the αCD40- treated cohort from the 1,252 TME features quantified above (**Fig. 2D** and **Methods**). EN models perform well on datasets as generated herein where there are more features than samples (tissue regions) (59). This is because EN models use mathematical regularization approaches to identify and select the most informative subset of features to make model predictions while accounting for feature collinearity (60). Further, our approach was unbiased as we did not perform any prior feature selection and instead provided all 1,252 TME features to the models, leveraging the EN algorithm’s ability to perform aggressive feature selection within model training. Finally, SHAP values were used to interpret cellular biology underpinning model predictions (43). SHAP values denote the relative importance of a given feature in driving a model’s prediction – this method has been used to explain machine learning predictions in prior cancer studies (38, 61, 62).

### Machine learning models classify αCD40-treated TMEs as having reduced T cell exhaustion phenotypes

To reveal the impact of αCD40 therapy on T cell exhaustion phenotype, we first trained four EN classifier models – one per histopathologic annotation (**Fig. 1B**; T, IA, TAS, and NAP) – to predict the treatment status of the tissue. All models, regardless of histopathologic site, performed well, as measured by the accuracy, F1 score, and area under the receiver operating characteristic curve (AUC) for test sets of each of the models (**Fig. 3A** and **B**). Across the four models, accuracy ranged from 0.83 to 0.85, F1 score ranged from 0.73 to 0.89, and AUC ranged from 0.87 to 0.90. As models were trained to differentiate treatment-naive from αCD40-treated PDAC, high performance of all four models indicates that αCD40 modulates all types of histopathologic regions across the TME evaluated herein.

**Figure 3.**
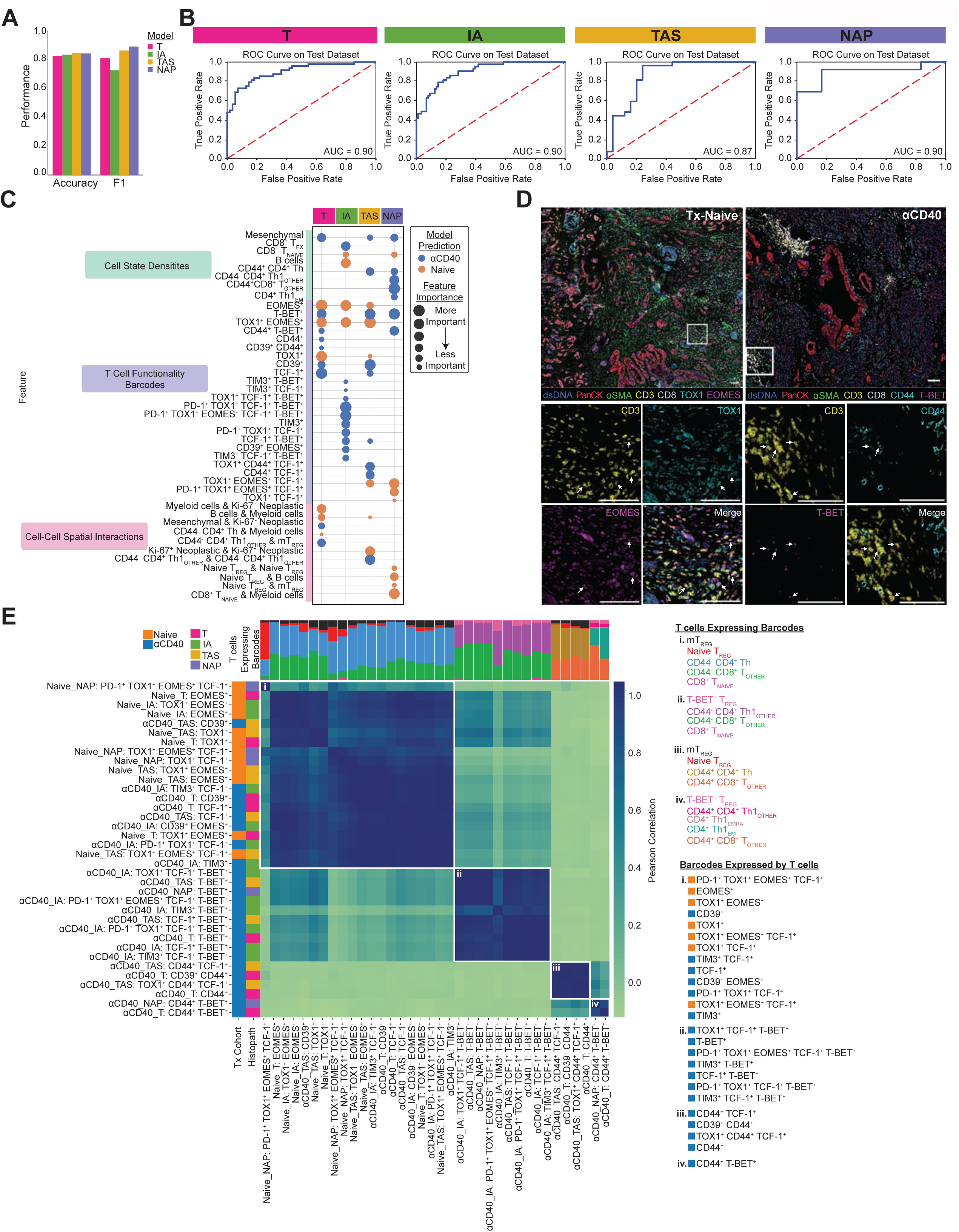
Machine learning models classify αCD40-treated TMEs as having reduced T cell exhaustion phenotypes. **A.** Bar chart showing accuracy and F1 score for each histopathologic model that predicts treatment status. **B.** ROC curve with corresponding AUC for each histopathologic model. **C.** Bubble chart showing top 15 features whose increased presence drove each histopathologic model to predict treatment-naive (orange) or αCD40-treated (blue). Features are grouped by TME feature type (density, barcode, interaction). Bubble size denotes relative importance of the feature for a given histopathologic model. Bubbles appearing multiple times in the same row indicate TME feature is a top feature across histopathologic models. **D.** Representative pseudo-colored mIHC images showing TOX1^+^ and/or EOMES^+^ CD3^+^ T cells in treatment-naive tissue (left) and CD44^+^ and/or T-BET^+^ CD3^+^ T cells in aCD40-treated tissue (right). **E.** Matrix showing correlations between top barcodes from the models with each other based on types and proportions of T cell states expressing the barcodes. Stacked bars at the top of correlation matrixes show proportions of T cell states expressing barcodes, with T cells color coded and listed for each group to the right of the heatmap, along with corresponding barcodes in each group. Leftmost columns are color coded according to which treatment group the presence of the barcode was predicted by the model, followed by the histopathologic site the model was derived from.

To identify which combinations of features drove model predictions, and thus reveal how αCD40 therapy impacted T cells in the PDAC TME, we calculated SHAP values for each of the four models (**Supplementary Fig. S2A**). The top 30 weighted features out of 1,252 features in total accounted for the majority of feature importance according to SHAP values (T model: 84%; IA model: 74%; TAS model: 87%; NAP model: 92%), while remaining features exhibited limited impact on the models. Comparison of the top 15 features driving model predictions for each of the four histopathologic models revealed that 13 features were top contributors across multiple models (**Fig. 3C**), indicating some amount of shared T cell densities, differentiation states, and spatial organizations across histopathologic sites within a given treatment cohort. To further bolster the machine learning results, we compared the normalized values for each of the top features detected by SHAP analysis between treatment-naive samples versus αCD40-treated samples within each histopathologic site (**Supplementary Fig. S2B, C, D, and E**). Following correction for multiple hypothesis testing, all 15 features derived from the T model were significantly different between treatment cohorts, 14 of 15 features derived from the IA and TAS models were significantly different between treatment cohorts, and 9 of 15 features derived from the NAP models were significantly different between treatment cohorts. Given that NAP sites had the fewest number of ROIs present in the data, it is unsurprising that fewer features were statistically significant.

Overall, the models identified αCD40-treated TMEs as containing increased densities of mesenchymal fibroblast-like cells and several T cell states, including three CD4^+^ T helper populations and two antigen-experienced CD8^+^ T cell populations (TEX and CD44^+^ TOTHER) (**Fig. 3C**, **Supplementary Fig. S2B, C, D, and E**), as compared to treatment-naïve TMEs, which contained increased densities of naive CD8^+^ T cells (although not significantly different) and B cells. In addition to cell state densities, analysis of T cell functionality barcodes revealed that αCD40-treated TMEs contained increased densities of T cells expressing combinations of T-BET, CD44, CD39, TIM3, and TCF-1 (**Fig. 3C**, **Supplementary Fig. S2B, C, D, and E**). On the other hand, treatment-naive TMEs contained increased densities of T cells expressing combinations of TOX1 and EOMES, concordant with mIHC stained tissue images (**Fig. 3D**). Finally, spatial interactions involving CD4^+^ Th1 cells were more likely identified within the αCD40-treated cohort, whereas interactions involving myeloid cells, naive TREGs (not significantly different), and Ki-67^+^ neoplastic epithelial cells were more likely to be associated with treatment-naive tissue (**Fig. 3C**, **Supplementary Fig. S2B, C, D, and E**). Altogether, these results indicate αCD40 TMEs contained increased presence of T cells in close spatial proximity to one another – in particular, CD4^+^ T helper cells – with reduced exhaustion profiles, as compared to treatment-naive TMEs.

Of the top features across all four models, the majority of features were densities of T cell functionality barcodes. As all barcodes present on any T cell (regardless of the T cell state) were provided to the machine learning models, we sought to understand the types of T cells expressing each of the predictive barcodes to determine whether there were patterns in the types and proportions of T cell states expressing each barcode. We ensured that only T cells within regions of the histopathologic site the predictive model was derived from were counted, as well as from the treatment cohort the barcoded T cell was predictive of. We then correlated barcodes based off of the types and proportions of T cell states expressing each barcode (**Fig. 3E**). Barcodes expressed by similar proportions of the same T cells were highly correlated and appear adjacent to each other in the heatmap. This analysis resulted in four clusters (i. – iv.) of barcodes, each distinct in types of T cells expressing them, and unrelated to histopathologic site, indicating that barcodes expressed by similar T cells exist across the TME. Barcodes belonging to cluster (i) were expressed by antigen-inexperienced (as defined by a lack of CD44 expression) CD8^+^ T cells, antigen-inexperienced CD4^+^ T helper cells, naive TREGs, and mTREGs. However, barcodes in cluster (i) ranged from TOX1^+^ and EOMES^+^ expression profiles when their increased densities were predictive of treatment-naive, to CD39^+^ TIM3^+^, and TCF-1^+^ expression profiles when their increased densities were predictive of αCD40 therapy. This result supports the notion that, while the same T cell types were present regardless of therapy exposure, their functional capacity differed following αCD40 treatment. Higher densities of barcodes in clusters (ii), (iii), and (iv) were all predictive of αCD40 therapy and included combinations of T-BET^+^, TIM3^+^, TCF-1^+^, and CD44^+^. These barcoded cells were expressed by distinct groups of T cell states, including multiple antigen-experienced CD4^+^ Th1 cell states, as well as T-BET^+^ TREGs, which have been reported to be similar to CD4^+^ Th1 cells in their function (63).

### Machine learning model classifies long disease-free survivors as having more T cell effector functionality following αCD40 therapy

The clinical trial from which the αCD40-treated specimens were derived was not designed to assess correlates with survival. However, we hypothesized that we could train machine learning models to accurately predict DFS for these patients, with the ultimate goal of identifying which combinations of TME features were associated with long versus short DFS. In contrast to the αCD40-treated patients who all received the same adjuvant treatment regimen (αCD40 therapy and chemotherapy), the patients from the treatment-naive cohort went on to receive varying adjuvant therapies. Thus, we did not train models to predict DFS for the treatment-naive cohort, as identifying the underlying biology driving improved survival for these patients would be challenging. Therefore, we trained EN classifier models to predict long or short DFS for tissue regions only from patients treated with αCD40 therapy. The median DFS timepoint (9.8 months) across all patients in the αCD40-treated cohort was used to segregate long and short disease-free survivors. Again, separate models were built for regions annotated as each histopathologic site, including T, IA, and TAS. There were too few NAP regions to build a model assessing DFS from the αCD40-treated patients for this histopathologic site. Only the model trained from IA regions performed well in predicting both long and short DFS, with an accuracy of 0.81, F1 score of 0.88, and AUC of 0.77 (**Fig. 4A** and **B**).

**Figure 4.**
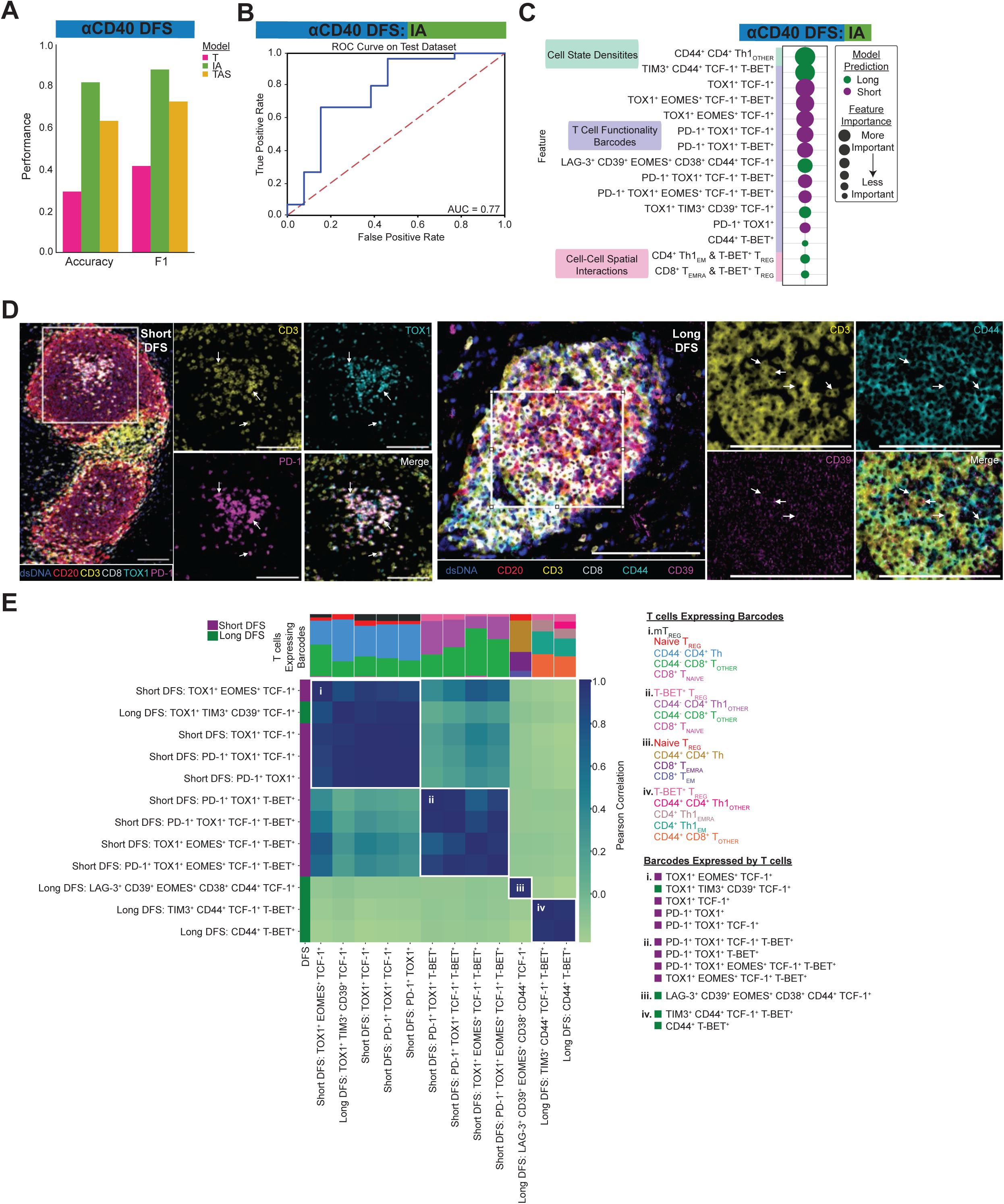
Machine learning model classifies long disease-free survivors as having more T cell effector functionality following αCD40 therapy. **A.** Bar chart showing accuracy and F1 score for each histopathologic model predicting αCD40 DFS. **B.** ROC curve with corresponding AUC for the IA histopathologic model. **C.** Bubble chart showing the top 15 features whose increased presence drove the IA model to predict short DFS (purple) or long DFS (green). Features are grouped by TME feature type (density, barcode, interaction). Bubble size denotes relative importance of the feature. **D.** Representative pseudo-colored mIHC images showing TOX1^+^ and/or PD-1^+^ CD3^+^ T cells in short DFS tissue (left) and CD44^+^ and/or CD39^+^ CD3^+^ T cells in long DFS tissue (right). Scale bars = 100 µm. **E.** Matrix showing correlations between the top barcodes from the IA model with each other based on types and proportions of T cell states expressing the barcodes. Stacked bars at the top of the correlation matrix show proportions of T cell states expressing the barcodes, and T cells are color coded and listed for each group to the right of the heatmap, along with the corresponding barcodes in each group. Leftmost column is color coded according to which DFS group the presence of the barcode predicted by the model.

To identify which combinations of features drove predictions for the IA-derived model, and thus hypothesize potential mechanisms contributing to improved DFS following αCD40 therapy, we followed a similar model interpretation analysis as described above. SHAP values were calculated to highlight feature importance (**Supplementary Fig. S3A**, **Fig. 4C**), and the top 30 features out of 1,252 features fed into the model accounted for 78% of feature importance. All of the top 15 features detected by the SHAP analysis for the IA-derived model were significantly different between DFS groups (**Supplementary Fig. S3B**), thus demonstrating how machine learning can be used to reveal potential combinations of candidate biomarkers of DFS in the PDAC TME.

Of note, an increased density of CD44^+^ CD4^+^ Th1 cells was observed in IA regions from patients with long DFS, as compared to those with from patients with short DFS (**Fig. 4C**, **Supplementary Fig. S3B**). In addition, we found increased spatial interactions between CD4^+^ Th1 TEM cells or CD8^+^ TEMRA cells and T-BET^+^ TREGs to be associated with long DFS rather than short DFS (**Fig. 4C**, **Supplementary Fig. S3B**). Interestingly, of the three types of TME features, densities of T cell functionality barcodes were the most common feature predictive of DFS, accounting for 12 of the top 15 features. Eight of these barcode densities were associated with short DFS, while the remaining four were associated with long DFS (**Fig. 4C**, **Supplementary Fig. S3B**). A number of biomarkers segregated based on DFS status (**Fig. 4C, D**); namely, TOX1 was expressed across all eight barcodes whose increased densities were associated with short DFS, and PD-1 expression was found exclusively in five of the eight barcodes whose increased densities were associated with short DFS. In contrast, expression of CD44, CD38, CD39, TIM3, and LAG-3 were unique to the four barcodes whose increased densities were associated with long DFS (**Fig. 4C, D**).

To identify whether the types and amounts of T cell states expressing each of the predictive barcodes were similar, we correlated the barcodes based off of the proportions of T cells expressing each of them (**Fig. 4E**). We found four distinct clusters of barcodes, and within each cluster, the barcodes were expressed by similar types of T cell states and in similar proportions. All but one of the barcodes in clusters (i) and (ii) were among the features whose increased densities were associated with short DFS and were expressed by (i) antigen-inexperienced CD8^+^ T cells, antigen-inexperienced CD4^+^ T helper cells, naive TREGs, and mTREGs; and (ii) antigen-inexperienced CD8^+^ T cells and CD4^+^ Th1 cells, as well as T-BET^+^ TREGs. In contrast, increased densities of all barcodes in clusters (iii) and (iv) were associated with long DFS. Cluster (iii) consisted of one barcode that was uniquely expressed by CD8^+^ TEM and TEMRA cells, CD44^+^ CD4^+^ T helper cells, as well as naive TREG cells. Finally, cluster (iv) barcodes were expressed by CD44^+^ CD8^+^ T cells, CD4^+^ Th1 TEM and TEMRA cells, CD44^+^ CD4^+^ Th1 cells, and T-BET^+^ TREGs. In summary, these findings indicate αCD40 therapy stimulates an anti-tumor T cell response, characterized by presence of CD44^+^ T cells, in particular CD4^+^ Th1 cells, and is associated with prolonged DFS time within IAs.

### Cellular neighborhood analysis identifies spatial organization of T cells to correlate with DFS following αCD40 therapy

The majority of top TME feature types driving the αCD40 DFS IA model predictions were presence of specific barcoded T cells, supporting the notion that these T cell subsets likely play a major role in extending DFS. However, the spatial organization of the specific barcoded T cells identified by the machine learning analyses remained unknown, as the spatial interaction features did not account for T cell functionality barcodes. TME spatial architecture is associated with clinical outcomes across cancer types (24-27, 31); thus, we aimed to identify the spatial neighbors of the top barcoded T cells whose densities were associated with DFS to better understand their potential role in prolonging DFS following αCD40 therapy within IA regions.

To quantify the spatial organization of the tissue, we performed a recurrent cellular neighborhood (RCN) analysis across all IA regions within the αCD40-treated cohort (26, 64). Here, we established a 60 µm radius (47-50) around every cell as its neighborhood and identified all cells within that neighborhood (**Fig. 5A**). We then clustered all cells according to the cellular neighborhoods’ compositions (**Methods**). This resulted in seven RCNs (**Supplementary Fig. S4A**) – each one representing spatial neighborhoods present across multiple IAs that were distinct in proportions and types of cell states located within the neighborhood. The average cellular composition of each RCN is shown (**Fig. 5B**). We confirmed that no single RCN dominated the IA regions analyzed (**Supplementary Fig. S4B** and **C**) and that no RCN was exclusively derived from any single IA region or patient (**Supplementary Fig. S4D** and **E**). Upon viewing the scatterplot reconstructions of regions, clearly defined spatial patterns within IAs were revealed (**Fig. 5C**, **Supplementary Fig. S4F**). For example, cells in RCN1, whose neighborhood consisted of majority B cells, were often found to be spatially clustered together, potentially representing “germinal center”-like pockets within IA regions.

**Figure 5.**
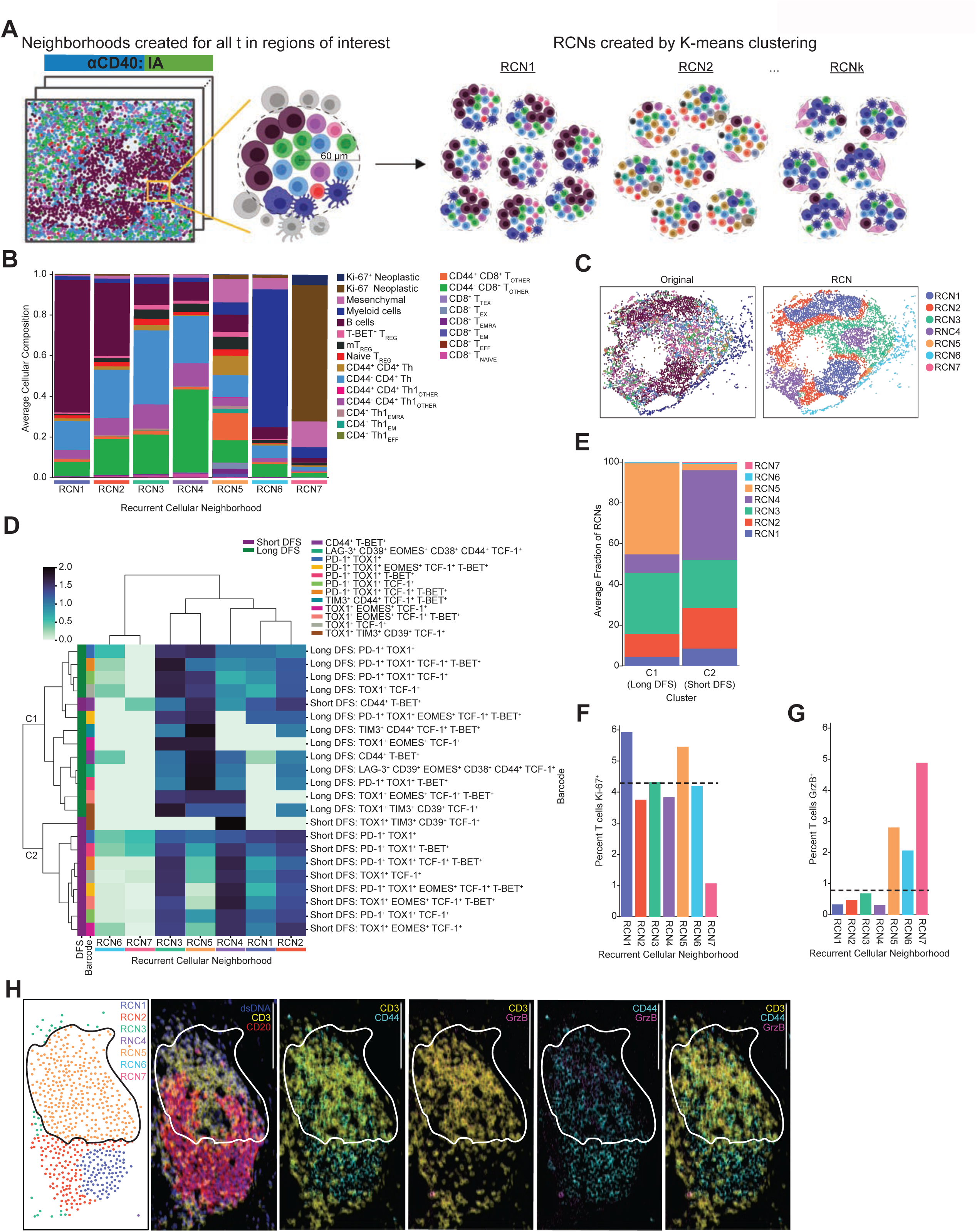
Cellular neighborhood analysis identifies spatial organization of T cells to correlate with DFS following αCD40 therapy. **A.** Schematic depicting RCN analysis. Cellular neighborhoods were defined by identifying all cells within a 60 µm radius of a given cell. Neighborhoods were calculated for all cells in αCD40-treated IA regions. Neighborhoods were then grouped using K-means clustering to identify RCNs. Created with BioRender.com. **B.** Stacked bar chart showing average cellular composition of each of seven RCNs from the αCD40-treated IA regions. Bars are colored by cell state and represent average proportions (out of 1.0) of each cell state present in neighborhoods assigned to each RCN. **C.** Representative IA tissue region as depicted by scatterplot reconstructions. Each dot represents a cell present in the IA, and each cell is colored by its original cell state phenotype (left scatterplot) or RCN assignment (right scatterplot). **D.** Heatmap showing top T cell barcodes from the αCD40 IA DFS model clustered by proportion of RCNs the T cell barcodes were assigned to. Rows are barcoded T cells from IA regions from patients associated with short DFS or long DFS ordered by hierarchical clustering of their RCN assignment. Columns are RCNs used as clustering features. Proportion of RCNs was normalized using a log10+1 transformation prior to clustering. Leftmost columns are color coded by DFS group followed by barcode. **E.** Stacked bar chart showing average fraction of RCNs barcoded T cells were assigned to for each of two hierarchically clustered groups (C1 or C2). **F.** Bar chart showing percentage of T cells expressing Ki-67 residing in each of seven RCNs for αCD40 IA regions. Horizontal dashed line represents percentage of Ki- 67^+^ T cells across all αCD40 IA regions, regardless of RCN assignment. **G.** Bar chart showing percentage of T cells expressing GrzB residing in each of seven RCNs for αCD40 IA regions. Horizontal dashed line represents percentage of GrzB^+^ T cells across all αCD40 IA regions, regardless of RCN assignment. TREG populations were excluded from this analysis. **H.** Representative IA region from a patient with long DFS with cells colored by RCN assignment in the upper left scatterplot. Remaining images show mIHC staining of GrzB^+^ CD44^+^ CD3^+^ T cells localized within RCN5. Scale bars = 100 µm.

Given that our goal was to identify neighbors surrounding the top barcoded T cells from the αCD40 DFS model, we first identified which of the seven RCNs the barcoded T cells were assigned to. We then assessed whether there was a relationship between spatial organization of the barcoded T cells and clinical outcome by comparing the RCNs of barcoded T cells from patients with long DFS versus the RCNs of barcoded T cells from patients with short DFS. We performed unsupervised clustering of barcoded T cells from long and short DFS patients together based on proportions of the seven RCNs the barcoded T cells resided in (**Fig. 5D**). This resulted in two distinct clusters based on which RCNs the barcoded T cells resided in; the clusters also separated barcoded T cells from patients associated with long DFS versus short DFS. Cluster 1 (C1) included all barcoded T cells from patients with long DFS, as well as CD44^+^ T-BET^+^ barcoded T cells from patients with short DFS. Cluster 2 (C2) consisted of all remaining barcoded T cells from patients with short DFS.

The most striking difference between the two clusters was proportions of barcoded T cells residing in RCN5 (**Fig. 5E**). Of all RCNs, RCN5 contained the greatest proportion of CD44^+^ T cells, spanning both CD8^+^ T cells and CD4^+^ T helper lineages (**Fig. 5B**). 45% of T cells in C1 (long DFS) resided in RCN5, whereas only 3% of T cells from C2 (short DFS) resided in RCN5 (**Fig. 5E**). Given that C1 consisted of T cells from patients with long DFS and C2 consisted of T cells from patients with short DFS, our results highlight a relationship between cellular spatial organization and clinical outcome. Specifically, T cells correlated with long DFS were frequently found to be surrounded by CD44^+^ T cells, thus supporting the hypothesis that αCD40 therapy promotes T cell priming and/or recruitment of primed T cells to PDAC TMEs.

Finally, to further elucidate potential cellular mechanisms at play within various RCNs, we calculated proportions of T cells expressing Ki-67 or GrzB in each RCN and compared values to the overall proportion of Ki-67^+^ or GrzB^+^ T cells across all αCD40 IA regions. Given our prior findings that the majority of T cell barcodes whose increased density correlated with long DFS were assigned to RCN5 (**Fig. 5E**), we hypothesized that T cells assigned to RCN5 would possess increased proliferative and/or cytotoxic capabilities. We found a larger proportion of T cells expressing Ki-67 residing in RCN1 and RCN5 as compared to the overall T cell population (dashed line) (**Fig. 5F**). In addition, we found a larger proportion of T cells (excluding TREGs) expressing GrzB residing in RCN5, RCN6, and RCN7, as compared to the overall non-TREG T cell population (dashed line) (**Fig. 5G**). However, raw counts of GrzB^+^ T cells in RCN6 and RCN7 were low (RCN6: n=8 cells; RCN7: n=11 cells), whereas RCN5 contained the highest level of GrzB^+^ T cells across all RCNs (n=168). Visualization of a representative mIHC image depicts presence of GrzB^+^ T cells localized to RCN5 in an IA region from a patient with long DFS (**Fig. 5H**). Collectively, these results provide further support that T cells within RCN5 are likely activated and possess an effector phenotype capable of an anti-tumor cytotoxic response.

## DISCUSSION

In this study, we advance spatial proteomic imaging technology and leverage machine learning approaches to understand the role of T cell phenotypes and spatial organization in the complex TME of human pancreatic cancer. In contrast to previous single-cell spatial proteomic studies, which often group T cells together as broad populations, such as CD8^+^ T cells, CD4^+^ T cells, and TREGs (23, 25, 65-67), our novel mIHC panel was curated to phenotype T cells as one of 18 distinct states along with functionality status from 10 different markers, all while preserving the spatial orientation of each cell in the TME. In considering the full spectrum of T cell states in addition to their spatial organization within the TME, the data became increasingly complex, highlighting the need for quantitative analyses that consider integration of all features together, rather than one-off correlations which fail to represent intertwining biological mechanisms at play. We leveraged machine learning, as the models detect combinations and weights of tissue features associated with a given output, such as treatment status or DFS, and thus it is an appropriate and powerful tool for interrogating TME complexity. Through our machine learning approaches we identified TME features most associated with αCD40 therapy exposure or prolonged DFS out of the >1000 features that we measured. This study demonstrates the value of merging single-cell spatial proteomic assays with machine learning analyses to interrogate how immunotherapy modulates the PDAC TME and drives improved survival from a holistic perspective.

Despite the multitude of unique T cell states identified herein, our machine learning models were able to identify T cell subsets associated with anti-tumor characteristics and prolonged DFS. Consistent with our preclinical studies revealing that CD4^+^ T cells are a major contributor to PDAC immunity following αCD40 therapy (8), our machine learning models revealed that αCD40-treated patient tumors contain increased densities of effector memory cells specifically within the CD4^+^ Th1 lineage. Along the T cell spectrum, TEM cells are fully functional with some self-renewal capacities (68-70), and mediate potent anti-tumor immunity (71). No increase was observed in CD8^+^ memory T cell populations, supporting the notion that αCD40 therapy promotes immunological memory from the CD4^+^ T cell lineage in both mouse and human PDAC tissue. Our models also identified antigen-experienced CD4^+^ Th1 helper cells as the main cell type whose density associated with prolonged DFS following αCD40 therapy. This result is supported by two independent studies, including characterization of immune cells in biopsied PDAC liver metastases following CD40 agonism (72), and the second investigating primary resected PDAC TMEs after treatment with a granulocyte-macrophage colony-stimulating factor-secreting allogenic PDAC vaccine (GVAX) (rather than αCD40 therapy) (73). Both studies reported that presence of CD4^+^ T helper cells contribute to improved survival following immunotherapy in PDAC (72, 73). Here, we further characterized expression features of the CD4^+^ T helper cells as CD44^+^ Th1 cells, which correlated with improved outcomes. Reports have also highlighted the direct role of CD4^+^ T cells in mediating anti-tumor immunity, including via cytotoxicity (74) and production of effector cytokines like interferon-γ and tumor necrosis factor-α (75, 76). As such, we hypothesize that future therapies designed to harness effector and memory functions of CD4^+^ T helper cells following administration of αCD40 therapy may be clinically beneficial.

In addition to investigating presence of various T cell states, our machine learning analyses also shed light on localization and spatial organization of T cells within PDAC TMEs related to prolonged survival following αCD40 treatment. High performance of the IA-derived DFS model, coupled with poor performance of the T and TAS-derived DFS models, supports the notion that IAs are a major site of αCD40-induced immune response contributing to prolonged DFS. We previously reported an increase in number of IAs following αCD40 treatment in PDAC-bearing mice (77), and in the aforementioned GVAX study, survival was linked to increased CD4^+^ T helper pathway genes specifically within IAs (and not tumor regions) (73). Additionally, a recent study found the enrichment of gene signatures representing mature tertiary lymphoid structures in pre-treatment PDACs associated with improved survival in patients following treatment with varying chemoimmunotherapies (78). Our RCN analyses further revealed that the key T cells associated with prolonged DFS were often surrounded by antigen-experienced CD8^+^ T cells and CD4^+^ T helper cells, as well as a higher proportion of proliferating and cytotoxic T cells, as compared to all T cells regardless of spatial neighborhood. Collectively, the results indicate IAs may function as sites of T cell priming or second signal, promoting T cell activation and function in PDAC TMEs, contributing to prolonged DFS following multiple types of immunotherapies.

Our interrogation of IAs also revealed T cell features correlating with short DFS following αCD40 therapy, identifying T cell states that are concordant with dysfunctional tumor-infiltrating T cell phenotypes. TOX1^+^ T cells are at the far end of the exhausted T cell spectrum (14, 79), and TOX1 expression correlates with PD-1 on T cells and impaired immunotherapy response in hepatocellular carcinoma (80). Correspondingly, we observed that expression of TOX1 and/or PD-1 on CD8^+^ T cells and CD4^+^ T helper cells associated with shorter DFS following αCD40 therapy. Further, T cells linked to short DFS expressing TOX1 and/or PD-1 were largely CD44^-^ T cells, which may represent a population of antigen-naïve T cells that have aberrantly upregulated these markers, or could potentially represent T cells that have become terminally exhausted and differentiated due to repeated T cell receptor stimulation (and thus downregulated CD44 expression) (81). We also found increased presence of TOX1^+^ T cells within treatment-naive TMEs, indicating a baseline terminally exhausted T cell phenotype in PDAC. Interestingly, despite the presence of TOX1^+^ exhausted T cells within treatment-naive TMEs, we did not identify abundant expression of PD-1^+^ T cells in treatment-naive TMEs, in contrast to other tumors, such as melanoma (82), where PD-1^+^ T cells are abundant in TMEs. The paucity of PD-1^+^ T cells in PDAC TMEs could be a contributing factor to the failure of ICBs targeting PD-1 or PD-L1 in the vast majority of patients with PDAC (83). Together, our data support the conclusion that TOX1, but not PD-1, is a dominant feature of exhausted T cells in PDAC. As such, therapies that modulate TOX1^+^ T cells in the TME – such as αCD40 agonism in this study – may improve clinical outcomes for patients with PDAC. An overview of the main TME features detected by our machine learning models in this study are visually described (**Fig. 6**).

**Figure 6.**
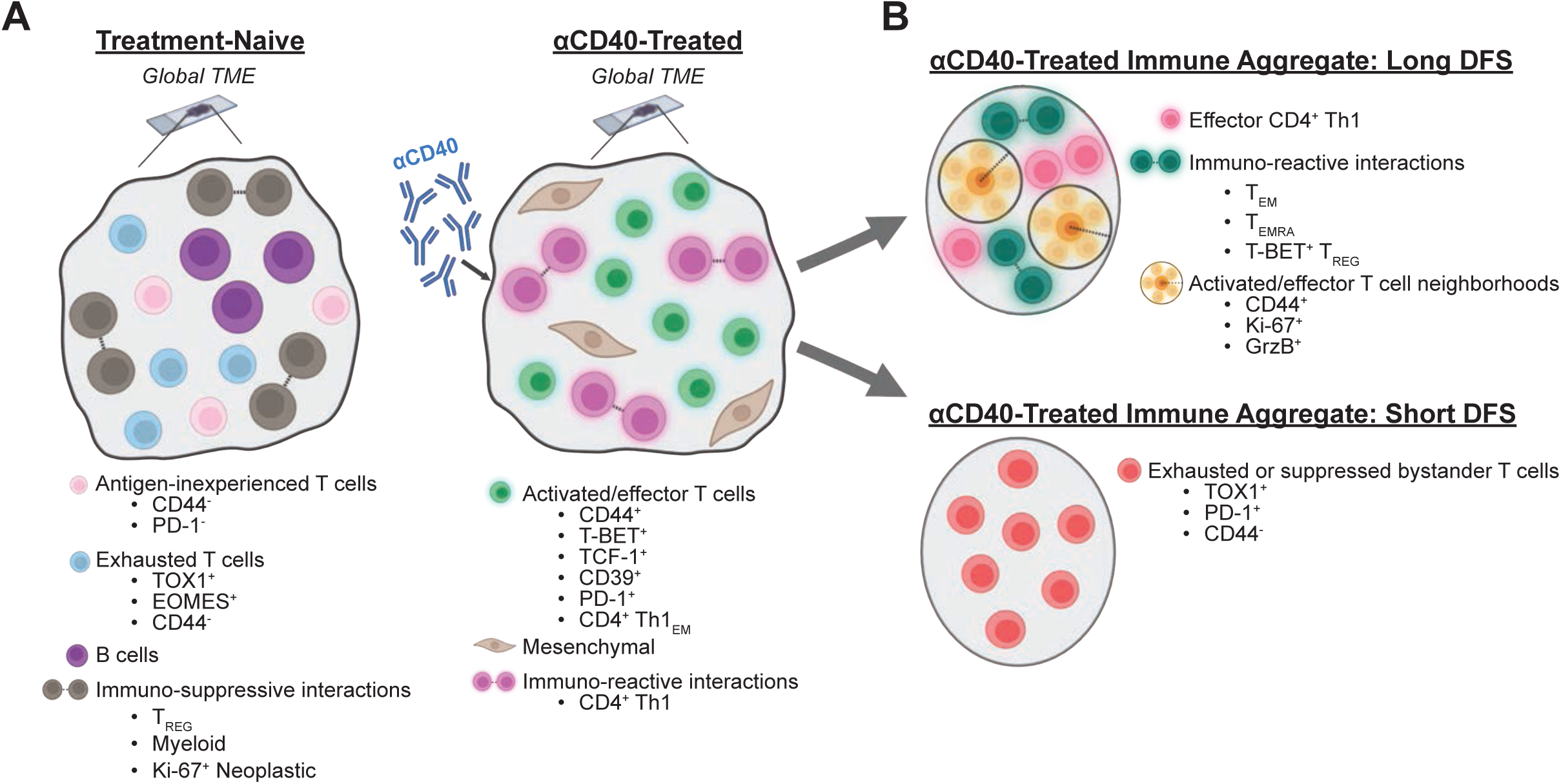
Spatial features of T cells associated with αCD40 therapy and prolonged DFS in the PDAC TME. **A.** T cell subsets that best define resected tumor samples from treatment-naïve (left) or αCD40-treated (right) patients. In the absence of therapy, T cells appear in an exhausted state, while T cells present with activated and effector phenotypes after CD40 agonism. Created with BioRender.com. **B.** T cell phenotypes in IAs from αCD40-treated patients associated with long (top) or short (bottom) DFS. IAs from patients with long DFS are characterized by the presence of spatial neighborhoods of effector T cells capable of proliferating and cytotoxicity, while IAs from patients with short DFS have a preponderance of exhausted T cell states. Created with BioRender.com
.

Notably, the dataset we used to conduct this study was unique in several aspects – including therapy administered, mIHC panel used, and histopathologic sites assayed – making absolute validation of our results challenging and highlighting the need for additional studies on larger cohorts to address the potential of building a model capable of more generalized predictions. Patient samples were collected from multiple institutions per treatment cohort, yet our machine learning models were still able to accurately classify patient samples according to therapy. This suggests any institutional or technical differences in tissue processing were not driving features in model predictions. Importantly, the machine learning models we trained performed comparably to or better than several other models derived from similar studies (36, 38, 84-88). Additionally, our biological conclusions on impacts of αCD40 are concordant with several prior studies (8, 14, 72, 73, 77, 79, 80, 83), providing further support for our methods and findings. Future antibody panels may incorporate additional markers, such as chemokine receptors, to further characterize key T cell subsets, or additional cell lineage markers, such as myeloid cell markers (89), to further phenotype cell-cell interactions. It should be noted that in our study, treatment with αCD40 did not prolong DFS as compared to the treatment-naive cohort, and similar machine learning analyses on data from clinical trials that did improve outcome will be useful to validate our conclusions where the patient cohort size is better powered for survival analyses. Finally, tumors were resected 12 days after αCD40 administration, so it is possible that the T cells involved in prolonged DFS did not have sufficient time to transit beyond IAs and into surrounding TMEs following priming. Further analyses investigating timing of T cell trafficking into various sites within PDAC TMEs are necessary to determine if analysis of T or TAS regions sampled at later timepoints following treatment could be used to assess clinical outcome for these patients.

This study provides proof-of-principle for leveraging computational approaches to evaluate highly multiplexed cancer datasets and supports the notion that similar analytic approaches could be utilized in future studies to identify important, and otherwise inconspicuous alterations in TMEs correlating with patient treatment or response. Future studies could expand upon these findings to target pathways identified via this approach to benefit patients who do not respond to current treatment strategies.

## ACKNOWLEDGEMENTS

The authors thank Drs. David L. Bajor, E. Gabriela Chiorean, Daniel A. Laheru, and Mark H. O’Hara for efforts as site investigators for the neoadjuvant selicrelumab (CD40 agonist) clinical trial. We also thank John Wherry for helpful discussion related to panel design and marker selection. This research was supported by the National Cancer Institute (NCI) of the National Institutes of Health grants T32CA254888 (K.E.B.), P50 CA127003 (J.A.N.), R01 CA248857 (J.A.N.), R01 CA205406 (J.A.N.), R01 CA169141 (J.A.N.), R35 CA197735 (J.A.N.), U01 CA250549 (J.A.N.), U24CA231877 (J.G.), and U2CCA233280 (J.G.), the DFCI Hale Family Center for Pancreatic Cancer Research (J.A.N. and B.M.W.), the Lustgarten Foundation Dedicated Laboratory program (J.A.N., B.M.W.), the Parker Institute for Cancer Immunotherapy (R.H.V. and K.T.B.), the Brenden-Colson Center for Pancreatic Care (L.M.C., S.S., C.B.B., K.T.B., N.K.), the Robert L. Fine Cancer Research Foundation (K.T.B.), funding from the Prospect Creek Foundation to the OHSU SMMART (Serial Measurement of Molecular and Architectural Responses to Therapy) Program (J.G.), and a generous startup package from the Knight Cancer Institute and the Brenden-Colson Center for Pancreatic Care (K.T.B.).

## AUTHOR CONTRIBUTIONS

*Conceptualization:* K.E.B., J.G., L.M.C., K.T.B.

*Formal analysis:* K.E.B., S.S., C.B.B., B.T.

*Funding acquisition:* J.G., L.M.C., K.T.B.

*Investigation:* K.E.B., S.S., C.B.B., K.B., N.K., E.E.F., A.D.C., J.A.N.

*Methodology:* K.E.B., S.S., C.B.B., K.B., N.K., B.T., J.G., L.M.C., K.T.B.

*Resources:* R.H.V., B.M.W., J.G., L.M.C., K.T.B.

*Supervision:* J.G., L.M.C., K.T.B.

*Visualization:* K.E.B., S.S., N.K.

*Writing – original draft:* K.E.B., S.S., J.G., L.M.C., K.T.B.

*Writing – review and editing:* all authors.

## CONFLICT OF INTEREST

R.H.V. is an inventor on licensed patents relating to cancer cellular immunotherapy and cancer vaccines, and mutant Kras specific T cell receptors; has received consulting fees from BMS; and receives royalties from Children’s Hospital Boston for a licensed research-only monoclonal antibody and from the University of Pennsylvania for licensed research cell lines. J.A.N. receives consulting fees from Leica Biosystems and research support from Natera. B.M.W. receives research funding from AstraZeneca, Celgene/BMS, Eli Lilly, Novartis, and Revolution Medicines, and consulting for Celgene, GRAIL, Ipsen, Mirati, Third Rock Ventures unrelated to the current work. C.B.B. is an employee of, and holds equity in, Akoya Biosciences, Inc. K.T.B. receives royalties from the University of Pennsylvania for licensed research cell lines and has received consulting fees from Guidepoint. L.M.C. has received reagent support from Cell Signaling Technologies, Syndax Pharmaceuticals, Inc., ZielBio, Inc., and Hibercell, Inc.; holds sponsored research agreements with Syndax Pharmaceuticals, Hibercell, Inc., Prospect Creek Foundation, Lustgarten Foundation for Pancreatic Cancer Research, Susan G. Komen Foundation, and the National Foundation for Cancer Research; is on the Advisory Board for Carisma Therapeutics, Inc., CytomX Therapeutics, Inc., Kineta, Inc., Hibercell, Inc., Cell Signaling Technologies, Inc., Alkermes, Inc., Raska Pharma, Inc., NextCure, Guardian Bio, AstraZeneca Partner of Choice Network (OHSU Site Leader), Genenta Sciences, Pio Therapeutics Pty Ltd., and Lustgarten Foundation for Pancreatic Cancer Research Therapeutics Working Group, Inc.

## SUPPLEMENTARY FIGURE LEGENDS

**Supplementary Figure S1.**
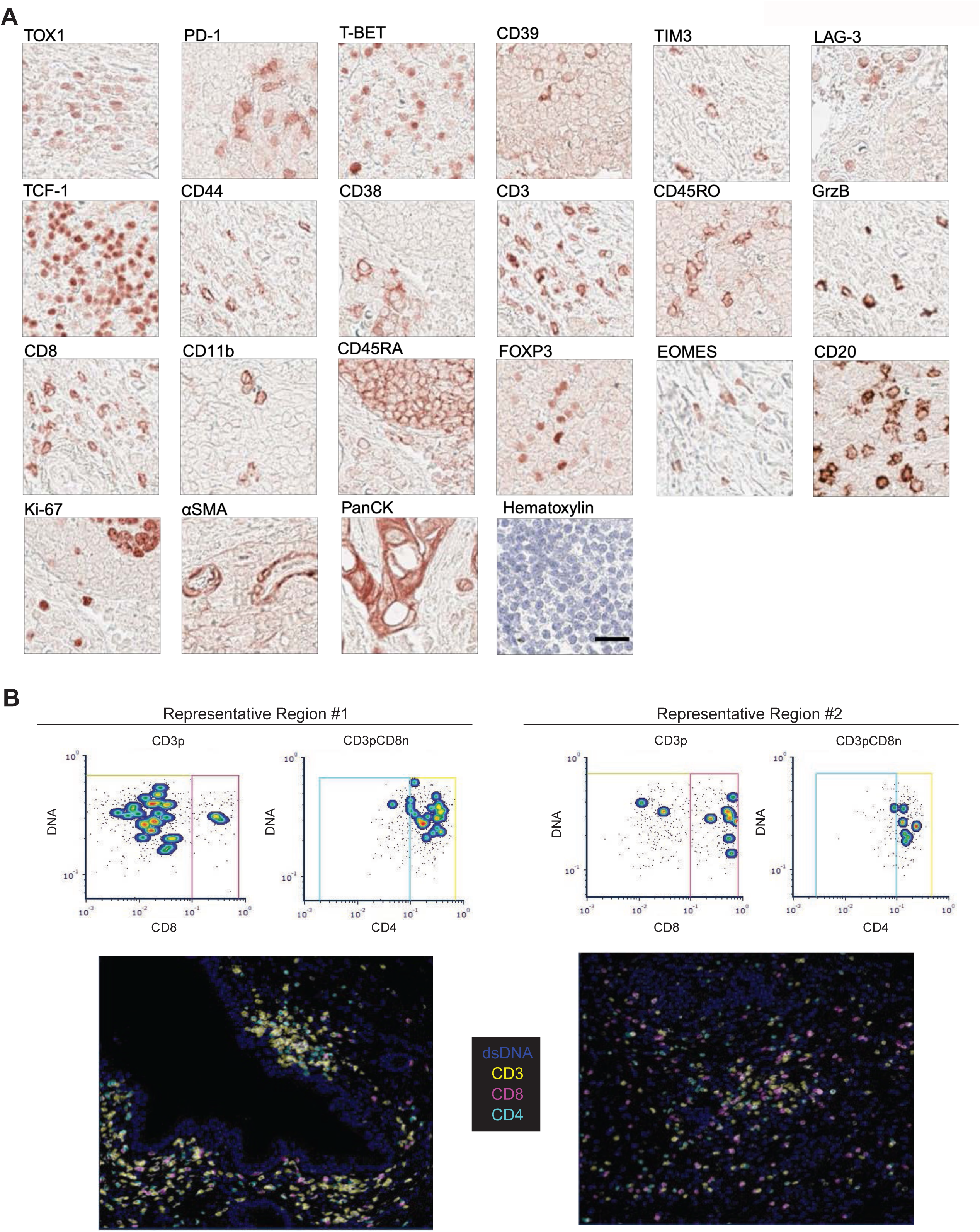
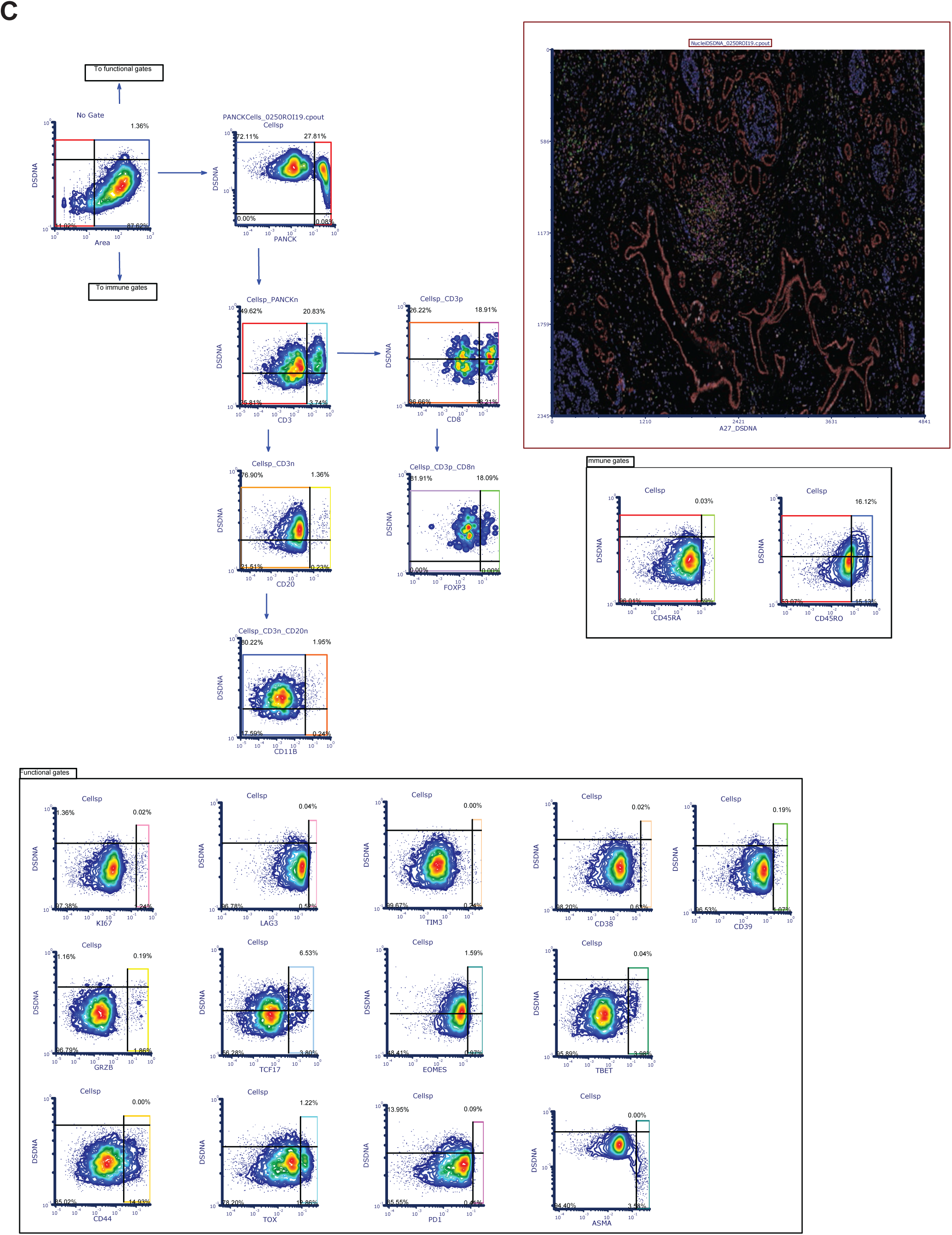
**A.** Representative IHC staining of each antibody used in sequence in the panel. Scale bar = 50 µm. **B.** Two representative regions stained with CD3, CD8, and CD4 antibodies. For each region, top images show gates for CD8 on CD3^+^ population (left) and CD4 on CD3^+^ CD8^-^ population (right), and bottom row shows pseudo-colored mIHC images. **C.** Hierarchical gating template used to phenotype cells using image gating cytometry in FCS Image Cytometry RUO.

**Supplementary Figure S2.**
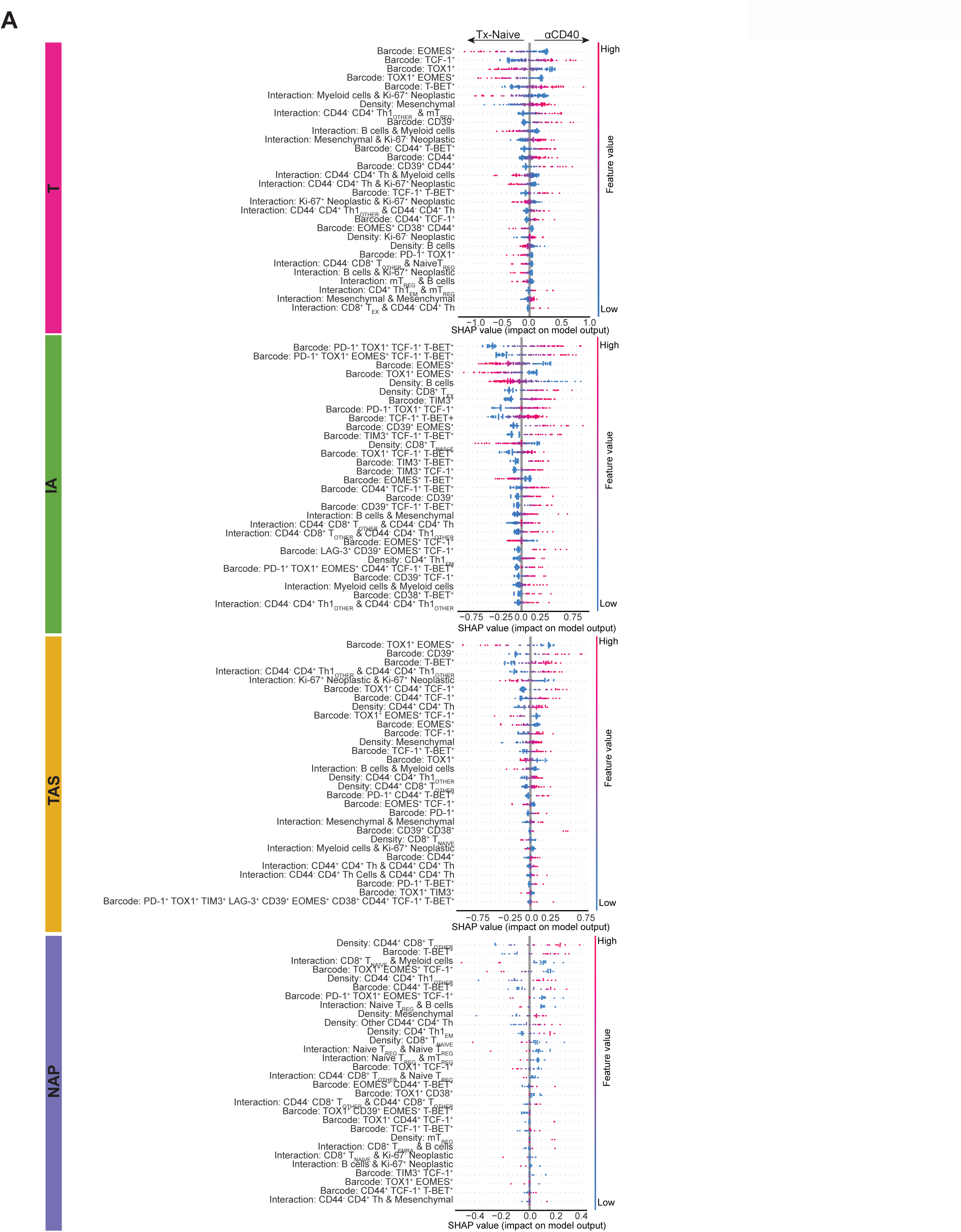
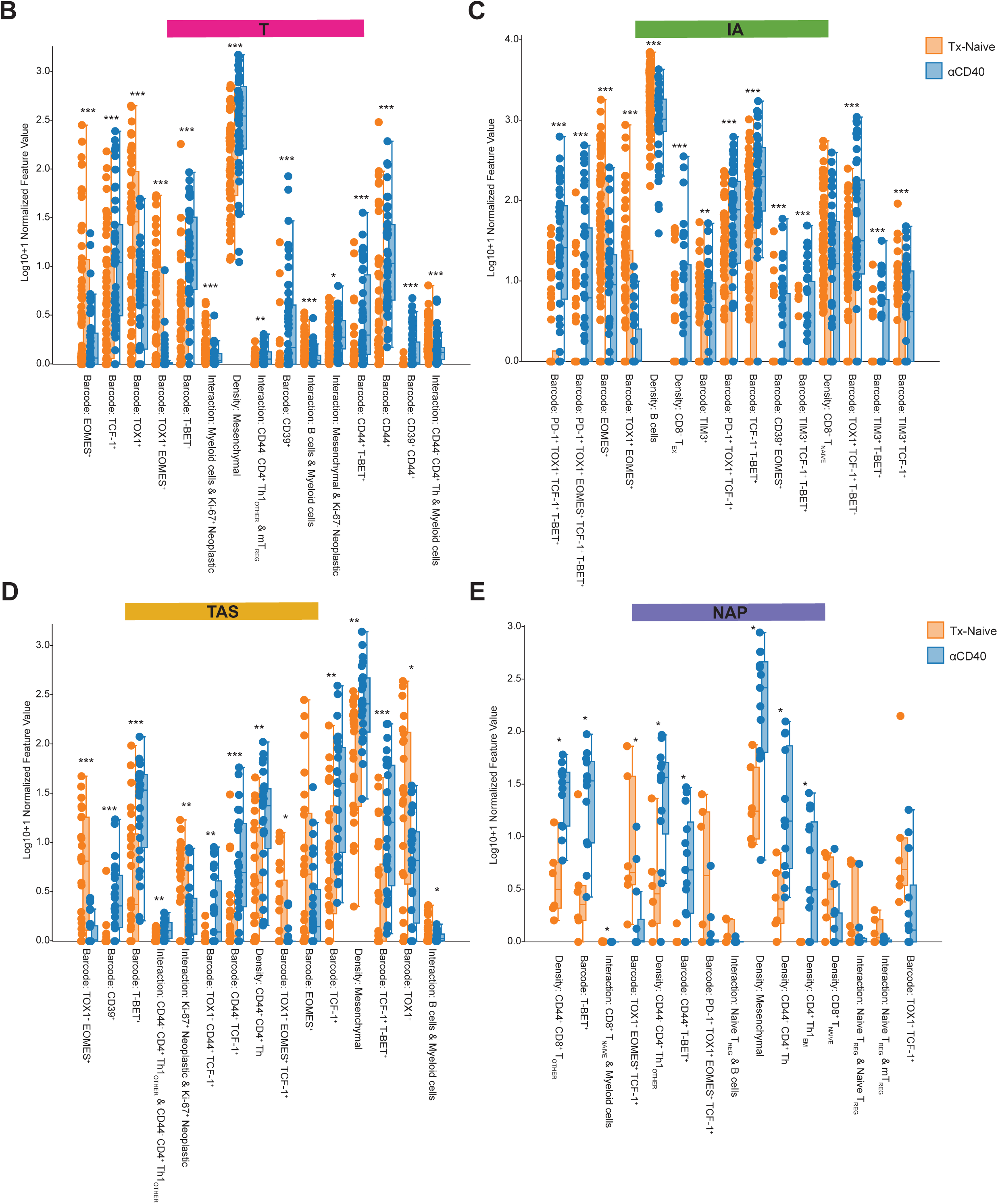
**A.** SHAP plots showing the top 30 features driving each histopathologic model. Features are ordered on the y-axis such that those with a larger impact on model’s predictions appear at the top of the SHAP plots. SHAP values are shown on the x-axis, with a value of zero (center) indicating no impact on the model, and negative or positive SHAP values predicting treatment-naive or αCD40-treated tissues, respectively. Red or blue dots indicate presence or absence, respectively, of the corresponding feature in the tissue. **B-E.** Box plots showing feature values for each of the top 15 features for models derived from T, IA, TAS, or NAP sites, respectively, split by treatment cohort. Each dot represents the log10+1 normalized feature value for one tissue region, inputted into the classifier model. Boxes = quartile 1 (Q1) to quartile 3 (Q3); whiskers = smallest and largest datapoints within 1.5*interquartile range (IQR) +/- Q3/Q1; solid line = median. Mann–Whitney *U*-test used to determine statistical significance. *P*-values corrected using the Benjamini–Hochberg procedure. *, *P* ≤ 0.05; **, *P* ≤ 0.01; ***, *P* ≤ 0.001. **B.** T site, n= 55 treatment-naive and n = 48 αCD40-treated regions per feature. **C.** IA site, n= 89 treatment-naive and n = 43 αCD40-treated regions per feature. **D.** TAS site, n = 25 treatment-naive and n = 27 αCD40-treated regions per feature. **E.** NAP site, n = 6 treatment-naive and n = 13 αCD40-treated regions per feature.

**Supplementary Figure S3.**
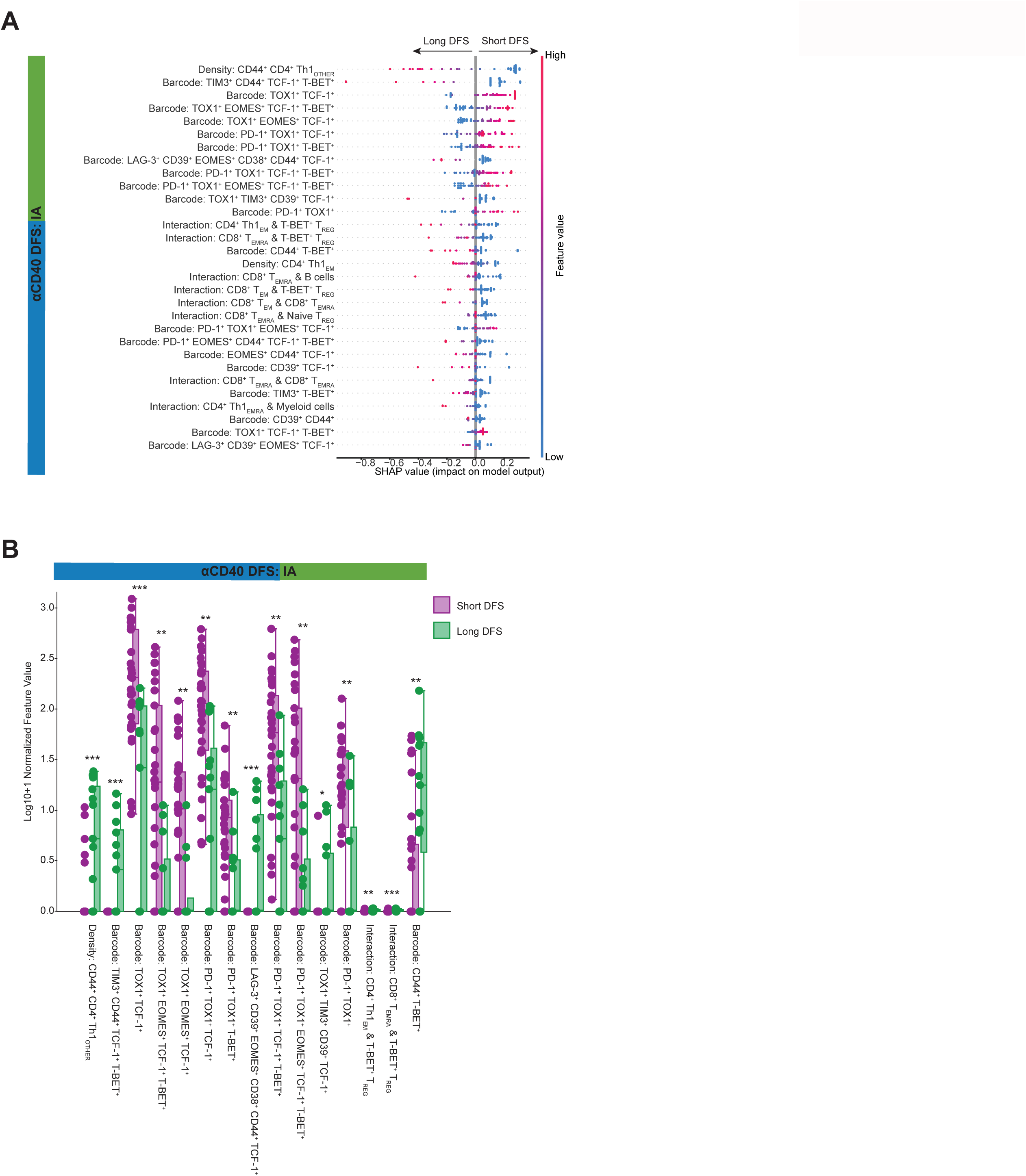
**A.** SHAP plot showing the top 30 features driving the IA model. Features are ordered on the y-axis such that those with a larger impact on the model’s predictions appear at the top of the SHAP plot. SHAP values are shown on the x-axis, with a value of zero (center) indicating no impact on the model, and negative or positive SHAP values predicting long DFS or short DFS, respectively. Red or blue dots indicate presence or absence, respectively, of the corresponding feature in tissues. **B.** Box plot showing feature values for each of the top 15 features for the model derived from IA regions of the αCD40 cohort split by DFS group (n = 30 regions from short DFS patients per feature; n = 13 regions from long DFS patients per feature). Each dot represents the log10+1 normalized feature value for one tissue region, which was inputted into the classifier model. Boxes = Q1 to Q3; whiskers = smallest and largest datapoints within 1.5*IQR +/- Q3/Q1; solid line = median. Mann–Whitney *U*-test used to determine statistical significance. *P*-values corrected using the Benjamini–Hochberg procedure. *, *P* ≤ 0.05; **, *P* ≤ 0.01; ***, *P* ≤ 0.001.

**Supplementary Figure S4.**
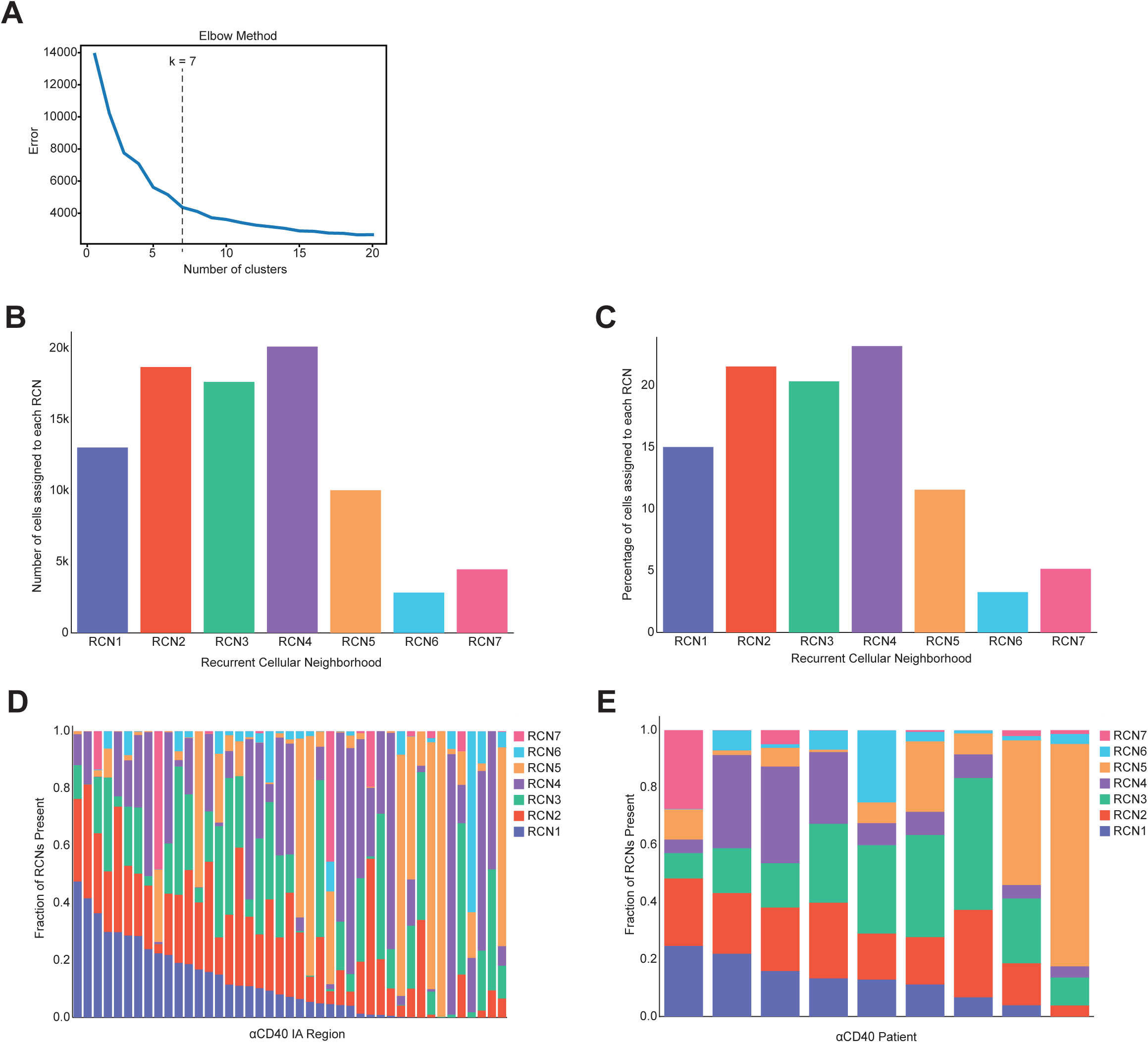
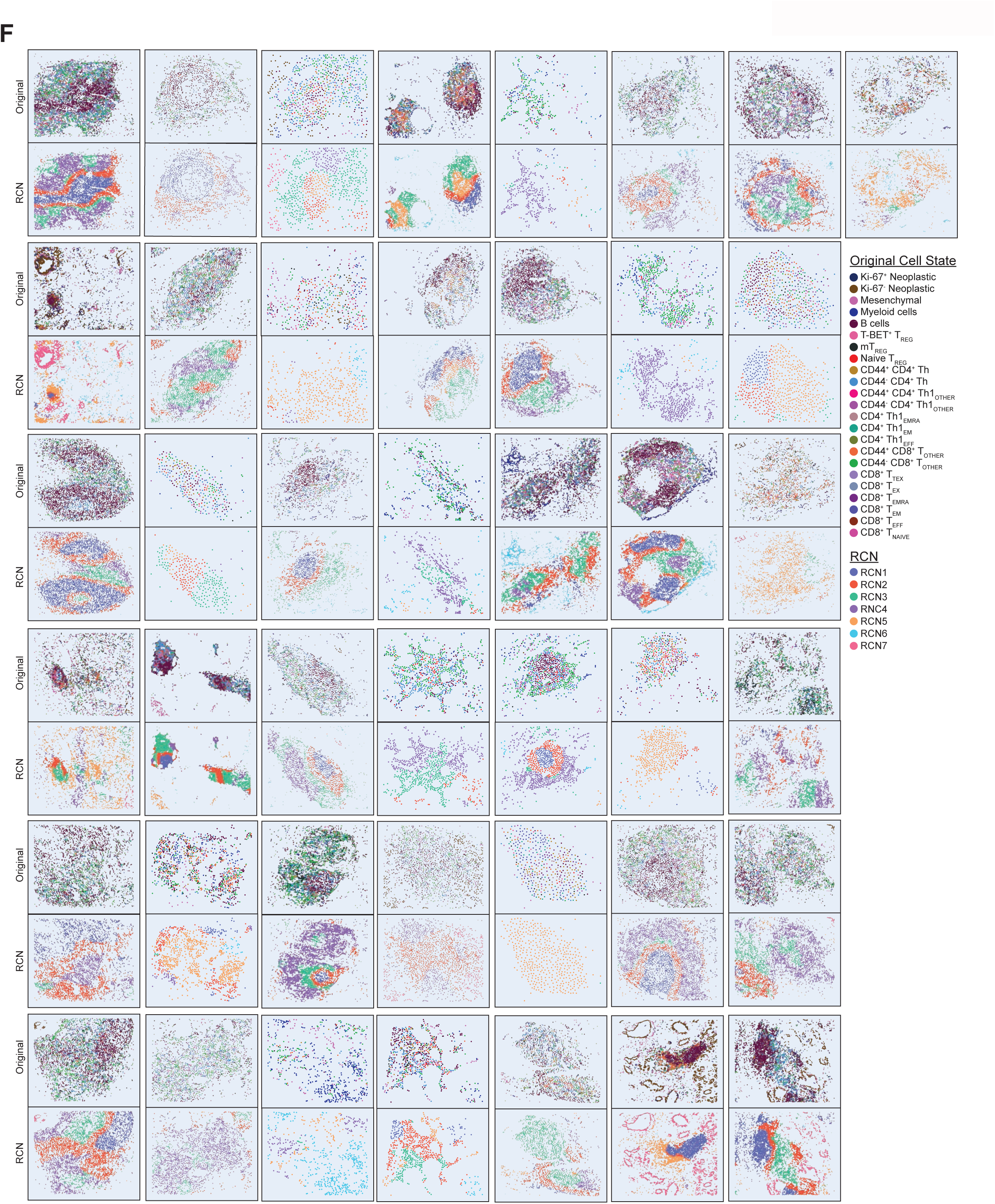
**A.** Elbow plot showing optimal number of RCNs (k=7) for grouping cellular neighborhoods. **B.** Bar chart showing the number of cells assigned to each of the seven RCNs across all αCD40 IA regions. **C.** Bar chart showing the percentage (out of 100) of cells assigned to each of the seven RCNs across all αCD40 IA regions. **D.** Stacked bar chart showing fraction (out of 1.0) of RCNs present per αCD40 IA region. **E.** Stacked bar chart showing average proportion (out of 1.0) of RCNs present in IA regions for each αCD40-treated patient. **F.** Scatterplot reconstructions for each αCD40 IA region. Each dot represents a cell present in the IA, and each cell is colored by its original cell state phenotype (top scatterplot) or RCN assignment (bottom scatterplot).

## SUPPLEMENTARY TABLE LEGENDS

**Supplementary Table S1.** Statistical comparison between the Liudahl et al. original PDAC cohort(35) and the selected subset used as Cohort 1 in this study. Mean value and standard error of the mean (SEM) shown for each variable. *P*-values computed using Fisher’s exact test for categorical variables, Wilcoxon rank-sum test for continuous variables, and log rank test for overall survival.

|  | **Original Cohort** | **Selected Subset** | ***P*** |
| --- | --- | --- | --- |
| OS (months) (mean ± SEM) | 27.34 ± 2.25 | 27.27 ± 4.75 | 0.90 |
| # cases OHSU/DFCI | 45/59 | 6/12 | 0.61 |
| CD8<sup>+</sup> T cell density (mean ± SEM) | 144.57 ± 11.49 | 152.51 ± 26.35 | 0.63 |
| CD4<sup>+</sup> T cell density (mean ± SEM) | 185.95 ± 12.45 | 215.15 ± 30.41 | 0.33 |
| CD20<sup>+</sup> B cell density (mean ± SEM) | 98.01 ± 10.68 | 97.73 ± 21.11 | 0.85 |
| Plasmablast density (mean ± SEM) | 12.13 ± 2.13 | 10.78 ± 2.86 | 0.89 |
| Plasma cell density (mean ± SEM) | 3.31 ± 0.69 | 2.51 ± 1.57 | 0.67 |
| Mast cell density (mean ± SEM) | 30.00 ± 2.51 | 21.25 ± 3.32 | 0.34 |
| Neutrophil/Eosinophil density (mean ± SEM) | 155.74 ± 29.28 | 220.87 ± 94.11 | 0.39 |
| Mature Dendritic cell density (mean ± SEM) | 3.32 ± 0.32 | 3.46 ± 0.67 | 0.59 |
| Immature Dendritic cell density (mean ± SEM) | 118.52 ± 11.86 | 108.79 ± 19.72 | 0.89 |
| CD163<sup>-</sup> Monocyte -Macrophage density (mean ± SEM) | 51.68 ± 4.17 | 37.79 ± 5.50 | 0.30 |
| CD163<sup>+</sup> Monocyte -Macrophage density (mean ± SEM) | 73.94 ± 7.16 | 62.31 ± 14.26 | 0.59 |
| CD8<sup>+</sup>/CD68<sup>+</sup> Ratio (mean ± SEM) | 1.86 ± 0.24 | 2.41 ± 0.89 | 0.21 |
| Total CD68<sup>+</sup> cell density (mean ± SEM) | 125.61 ± 9.84 | 100.09 ± 17.97 | 0.35 |

**Supplementary Table S2.** Table of antibodies used in mIHC panel.

| **Primary Antibody** | **Clone** | **Species** | **Supplier** | **Catalog number** |
| --- | --- | --- | --- | --- |
| TOX1 (Tox) | NAN488B | Rat | Abcam | ab237009 |
| PD-1 (PD1) | NAT105 | Mouse | Abcam | ab52597 |
| T-BET (TBET) | D6N8B | Rabbit | Cell Signaling Technologies | 13232S |
| CD39 | A1 | Mouse | Biolegend | 328202 |
| TIM3 | D5D5R | Rabbit | Cell Signaling Technologies | 45208S |
| LAG-3 (Lag3) | 17B4 | Mouse | Novus Biologicals | NBP1-97657 |
| TCF-1 (TCF1/7) | C6D39 | Rabbit | Cell Signaling Technologies | 2203 |
| CD44 | 156-3C11 | Mouse | Cell Signaling Technologies | 3570 |
| CD38 | 38C03 | Mouse | Thermofisher Scientific | MA5-14413 |
| CD3 | SP7 | Rabbit | Thermofisher Scientific | RM-9107-S |
| CD45RO | UCHL-1 | Mouse | Thermofisher Scientific | MA5-11532 |
| GrzB (Granzyme B) | Polyclonal | Rabbit | Abcam | ab4059 |
| CD8 | C8/144B | Mouse | Thermofisher Scientific | MA5-13473 |
| CD11b (CD11B) | EPR1344 | Rabbit | Abcam | ab133357 |
| CD45RA | F8-11-13 | Mouse | Abcam | ab59168 |
| FOXP3 (Foxp3) | 236A/E7 | Mouse | eBioscience | 14-4777-82 |
| EOMES | Polyclonal | Rabbit | EMD Millipore | AB2283 |
| CD20 | L26 | Mouse | Abcam | ab9475 |
| Ki-67 (KI67) | SP6 | Rabbit | Sigma/Cell Marque | 275R-14 |
| αSMA (Alpha-SMA) | Polyclonal | Rabbit | Abcam | ab5694 |
| PanCK (Pan-Cytokeratin) | AE1/AE3 | Mouse | Abcam | ab27988 |

**Supplementary Table S3.**
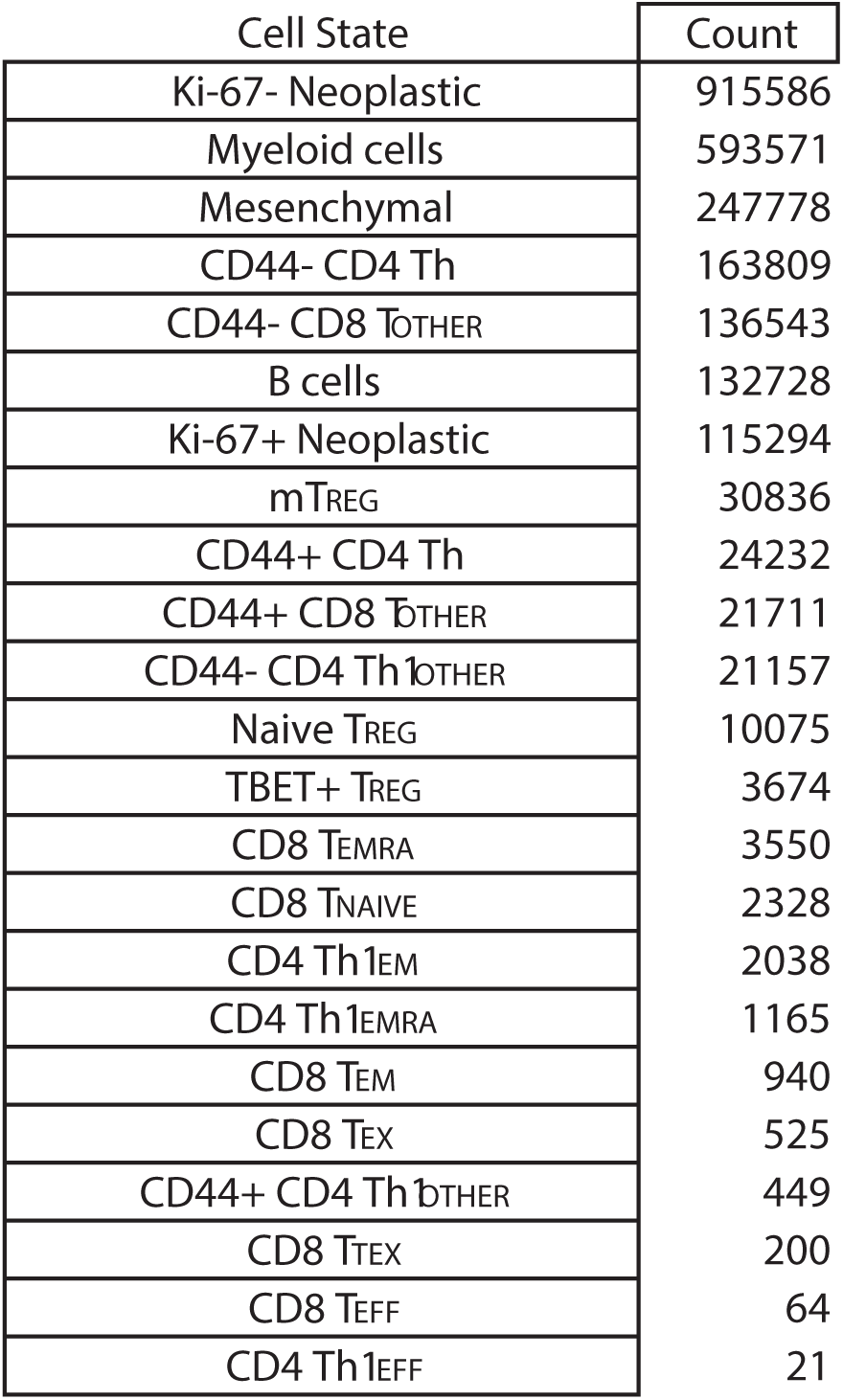
Raw counts of cell states defined by mIHC gating strategy present in the dataset.

**Supplementary Table S4.** Raw counts of T cells expressing each functionality barcode present in the dataset.

| T cell Functionality Barcode | Count |
| --- | --- |
| Negative for all | 144053 |
| TOX1+ | 51606 |
| TCF-1+ | 50915 |
| CD44+ | 19724 |
| T-BET+ | 15539 |
| EOMES+ | 15405 |
| TCF-1+ T-BET+ | 12775 |
| TOX1+ TCF-1+ | 10565 |
| CD44+ TCF-1+ | 7742 |
| EOMES+ TCF-1+ | 6118 |
| TOX1+ T-BET+ | 5000 |
| TOX1+ EOMES+ | 4876 |
| TOX1+ TCF-1+ T-BET+ | 4403 |
| EOMES+ CD44+ | 3505 |
| EOMES+ CD44+ TCF-1+ | 3318 |
| TOX1+ CD44+ | 3002 |
| PD-1+ TOX1+ TCF-1+ | 2684 |
| PD-1+ TOX1+ | 2669 |
| CD44+ T-BET+ | 2174 |
| CD44+ TCF-1+ T-BET+ | 2103 |
| EOMES+ TCF-1+ T-BET+ | 1862 |
| CD39+ | 1804 |
| TOX1+ EOMES+ TCF-1+ | 1629 |
| TOX1+ CD44+ TCF-1+ | 1591 |
| TOX1+ TIM3+ | 1574 |
| TOX1+ CD39+ | 1472 |
| CD38+ CD44+ | 1356 |
| PD-1+ TOX1+ TCF-1+ T-BET+ | 1262 |
| PD-1+ | 1177 |
| TOX1+ EOMES+ TCF-1+ T-BET+ | 1153 |
| EOMES+ T-BET+ | 1152 |
| TIM3+ | 1084 |
| TOX1+ CD38+ | 949 |
| CD38+ | 936 |
| PD-1+ TOX1+ EOMES+ TCF-1+ T-BET+ | 923 |
| TOX1+ EOMES+ CD44+ | 851 |
| PD-1+ TOX1+ EOMES+ | 840 |
| PD-1+ TCF-1+ | 833 |
| LAG-3+ | 819 |
| PD-1+ TOX1+ EOMES+ TCF-1+ | 785 |
| EOMES+ CD44+ TCF-1+ T-BET+ | 751 |
| PD-1+ TOX1+ TIM3+ LAG-3+ CD39+ EOMES+ CD38+ CD44+ TCF-1+ T-BET+ | 728 |
| TOX1+ EOMES+ T-BET+ | 582 |
| PD-1+ TOX1+ CD44+ TCF-1+ | 553 |
| TOX1+ CD38+ CD44+ | 512 |
| TOX1+ CD39+ EOMES+ | 495 |
| TOX1+ CD44+ TCF-1+ T-BET+ | 483 |
| CD39+ EOMES+ | 478 |
| LAG-3+ TCF-1+ | 473 |
| TOX11+ CD44+ T-BET+ | 461 |
| PD-1+ TOX11+ T-BET+ | 457 |
| CD39+ TCF-1+ | 448 |
| TOX11+ EOMES+ CD44+ TCF-1+ | 437 |
| TIM3+ CD44+ | 430 |
| PD-1+ T-BET+ | 386 |
| EOMES+ CD38+ CD44+ | 351 |
| CD38+ CD44+ TCF-1+ | 341 |
| EOMES+ CD38+ CD44+ TCF-1+ | 319 |
| PD-1+ TOX1+ CD39+ | 310 |
| TOX1+ LAG-3+ CD39+ | 304 |
| TOX1+ LAG-3+ | 303 |
| EOMES+ CD44+ T-BET+ | 292 |
| TIM3+ TCF-1+ | 281 |
| LAG-3+ T-BET+ | 269 |
| PD-1+ EOMES+ TCF-1+ | 256 |
| EOMES+ CD38+ | 252 |
| TIM3+ T-BET+ | 249 |
| PD-1+ TOX1+ TIM3+ LAG-3+ CD39+ EOMES+ CD38+ CD44+ TCF-1+ | 247 |
| PD-1+ TOX1+ CD44+ | 244 |
| CD38+ TCF-1+ | 228 |
| PD-1+ TCF-1+ T-BET+ | 216 |
| PD-1+ EOMES+ | 216 |
| TOX1+ TIM3+ T-BET+ | 208 |
| LAG-3+ EOMES+ TCF-1+ | 198 |
| LAG-3+ TCF-1+ T-BET+ | 193 |
| CD39+ CD44+ | 190 |
| TOX1+ CD38+ CD44+ TCF-1+ | 178 |
| PD-1+ TOX1+ LAG-3+ CD39+ | 175 |
| PD-1+ TOX1+ CD44+ TCF-1+ T-BET+ | 173 |
| LAG-3+ EOMES+ | 167 |
| PD-1+ CD44+ TCF-1+ | 166 |
| TOX1+ TIM3+ CD39+ CD44+ TCF-1+ T-BET+ | 165 |
| TOX1+ EOMES+ CD44+ TCF-1+ T-BET+ | 162 |
| CD39+ T-BET+ | 161 |
| TOX1+ TIM3+ CD44+ | 157 |
| TOX1+ TIM3+ TCF-1+ | 150 |
| PD-1+ CD44+ | 149 |
| TIM3+ TCF-1+ T-BET+ | 144 |
| TOX1+ EOMES+ CD44+ T-BET+ | 144 |
| CD39+ EOMES+ TCF-1+ | 144 |
| CD39+ TCF-1+ T-BET+ | 141 |
| PD-1+ TOX1+ EOMES+ T-BET+ | 139 |
| TOX1+ CD39+ EOMES+ TCF-1+ | 138 |
| TOX1+ TIM3+ CD39+ CD38+ CD44+ TCF-1+ T-BET+ | 137 |
| PD-1+ TOX1+ LAG-3+ CD39+ EOMES+ | 135 |
| TOX1+ EOMES+ CD38+ CD44+ TCF-1+ | 133 |
| TOX1+ CD39+ TCF-1+ | 133 |
| PD-1+ TOX1+ TIM3+ LAG-3+ CD39+ CD38+ CD44+ TCF-1+ T-BET+ | 129 |
| TOX1+ CD38+ T-BET+ | 125 |
| PD-1+ TOX1+ CD39+ EOMES+ | 124 |
| EOMES+ CD38+ TCF-1+ | 124 |
| TOX1+ LAG-3+ TCF-1+ | 122 |
| PD-1+ TOX1+ LAG-3+ | 118 |
| PD-1+ TOX1+ EOMES+ CD44+ TCF-1+ | 116 |
| TOX1+ LAG-3+ EOMES+ | 115 |
| PD-1+ EOMES+ TCF-1+ T-BET+ | 115 |
| TIM3+ EOMES+ | 113 |
| PD-1+ TOX1+ LAG-3+ CD39+ EOMES+ CD38+ CD44+ TCF-1+ | 112 |
| TOX1+ EOMES+ CD38+ CD44+ | 111 |
| PD-1+ TOX1+ TIM3+ | 109 |
| LAG-3+ CD44+ TCF-1+ | 109 |
| LAG-3+ CD39+ | 107 |
| LAG-3+ CD44+ | 107 |
| TOX1+ TIM3+ EOMES+ | 105 |
| TIM3+ CD44+ TCF-1+ | 103 |
| PD-1+ TOX1+ CD39+ EOMES+ TCF-1+ | 101 |
| CD38+ T-BET+ | 100 |
| TOX1+ CD39+ EOMES+ CD38+ TCF-1+ | 99 |
| TIM3+ CD39+ | 97 |
| TOX1+ TIM3+ TCF-1+ T-BET+ | 97 |
| PD-1+ TOX1+ EOMES+ CD44+ TCF-1+ T-BET+ | 97 |
| LAG-3+ EOMES+ TCF-1+ T-BET+ | 95 |
| CD39+ EOMES+ TCF-1+ T-BET+ | 94 |
| CD39+ CD38+ | 94 |
| TOX1+ LAG-3+ CD39+ EOMES+ | 94 |
| PD-1+ TOX1+ TIM3+ LAG-3+ EOMES+ CD38+ CD44+ TCF-1+ T-BET+ | 94 |
| CD38+ CD44+ T-BET+ | 93 |
| CD39+ EOMES+ CD38+ CD44+ | 93 |
| TOX1+ CD39+ T-BET+ | 91 |
| TIM3+ CD44+ T-BET+ | 91 |
| CD38+ TCF-1+ T-BET+ | 90 |
| CD39+ EOMES+ CD38+ | 90 |
| TOX1+ EOMES+ CD38+ | 88 |
| TOX1+ CD38+ TCF-1+ | 88 |
| TOX1+ CD39+ EOMES+ CD44+ TCF-1+ T-BET+ | 88 |
| TOX1+ LAG-3+ EOMES+ TCF-1+ | 86 |
| CD38+ CD44+ TCF-1+ T-BET+ | 85 |
| LAG-3+ EOMES+ CD44+ TCF-1+ | 83 |
| CD39+ CD38+ CD44+ | 83 |
| TOX1+ TIM3+ CD39+ EOMES+ CD38+ CD44+ TCF-1+ T-BET+ | 82 |
| TOX1+ TIM3+ LAG-3+ CD39+ EOMES+ CD38+ CD44+ TCF-1+ T-BET+ | 78 |
| PD-1+ TOX1+ LAG-3+ EOMES+ | 75 |
| TOX1+ EOMES+ CD38+ TCF-1+ | 75 |
| PD-1+ TOX1+ CD39+ TCF-1+ | 73 |
| PD-1+ CD44+ TCF-1+ T-BET+ | 71 |
| PD-1+ TOX1+ CD38+ | 70 |
| PD-1+ EOMES+ T-BET+ | 69 |
| PD-1+ TOX1+ TIM3+ LAG-3+ CD38+ CD44+ TCF-1+ T-BET+ | 67 |
| PD-1+ TOX1+ EOMES+ CD44+ | 67 |
| PD-1+ EOMES+ CD44+ TCF-1+ T-BET+ | 67 |
| TOX1+ CD39+ EOMES+ CD38+ CD44+ TCF-1+ | 64 |
| TOX1+ CD39+ TCF-1+ T-BET+ | 63 |
| PD-1+ TOX1+ TIM3+ LAG-3+ CD39+ EOMES+ CD44+ TCF-1+ T-BET+ | 63 |
| TIM3+ EOMES+ CD44+ | 62 |
| CD39+ EOMES+ T-BET+ | 61 |
| TOX1+ TIM3+ EOMES+ TCF-1+ | 61 |
| TIM3+ EOMES+ T-BET+ | 60 |
| TOX1+ LAG-3+ T-BET+ | 60 |
| PD-1+ EOMES+ CD44+ TCF-1+ | 59 |
| CD39+ EOMES+ CD44+ | 59 |
| PD-1+ TOX1+ LAG-3+ EOMES+ TCF-1+ | 59 |
| LAG-3+ CD39+ TCF-1+ | 57 |
| PD-1+ TOX1+ CD44+ T-BET+ | 57 |
| PD-1+ TOX1+ LAG-3+ CD39+ EOMES+ CD38+ TCF-1+ | 56 |
| LAG-3+ CD44+ TCF-1+ T-BET+ | 56 |
| TIM3+ EOMES+ TCF-1+ | 54 |
| PD-1+ TOX1+ LAG-3+ T-BET+ | 53 |
| PD-1+ TOX1+ CD39+ EOMES+ TCF-1+ T-BET+ | 53 |
| CD39+ EOMES+ CD38+ CD44+ TCF-1+ | 53 |
| PD-1+ TOX1+ TIM3+ TCF-1+ | 53 |
| CD39+ CD44+ TCF-1+ | 52 |
| TOX1+ TIM3+ CD44+ TCF-1+ | 52 |
| PD-1+ TOX1+ LAG-3+ TCF-1+ | 51 |
| TOX1+ CD39+ EOMES+ CD38+ | 50 |
| CD39+ EOMES+ CD38+ TCF-1+ | 49 |
| PD-1+ CD44+ T-BET+ | 48 |
| TOX1+ CD39+ EOMES+ CD38+ CD44+ | 47 |
| TIM3+ EOMES+ CD44+ TCF-1+ | 47 |
| PD-1+ TOX1+ CD39+ TCF-1+ T-BET+ | 47 |
| TOX1+ CD39+ CD44+ TCF-1+ T-BET+ | 47 |
| CD39+ EOMES+ CD44+ TCF-1+ | 46 |
| PD-1+ TIM3+ LAG-3+ CD39+ EOMES+ CD38+ CD44+ TCF-1+ T-BET+ | 45 |
| PD-1+ TOX1+ TIM3+ TCF-1+ T-BET+ | 44 |
| PD-1+ TOX1+ TIM3+ LAG-3+ EOMES+ CD38+ CD44+ TCF-1+ | 43 |
| TOX1+ CD39+ CD44+ | 43 |
| CD39+ EOMES+ CD44+ TCF-1+ T-BET+ | 43 |
| LAG-3+ CD38+ CD44+ TCF-1+ | 43 |
| TOX1+ TIM3+ CD39+ | 43 |
| LAG-3+ EOMES+ CD44+ | 42 |
| PD-1+ TOX1+ LAG-3+ CD39+ EOMES+ CD38+ CD44+ | 42 |
| TOX1+ CD38+ CD44+ TCF-1+ T-BET+ | 42 |
| EOMES+ CD38+ CD44+ TCF-1+ T-BET+ | 42 |
| LAG-3+ EOMES+ T-BET+ | 41 |
| TIM3+ CD39+ TCF-1+ | 41 |
| PD-1+ TOX1+ LAG-3+ CD39+ EOMES+ CD38+ | 41 |
| PD-1+ CD39+ | 40 |
| LAG-3+ EOMES+ CD38+ CD44+ TCF-1+ | 40 |
| PD-1+ TOX1+ LAG-3+ CD39+ EOMES+ CD38+ CD44+ TCF-1+ T-BET+ | 40 |
| LAG-3+ CD39+ EOMES+ | 39 |
| PD-1+ EOMES+ CD44+ | 39 |
| PD-1+ TOX1+ TIM3+ LAG-3+ CD39+ EOMES+ CD38+ TCF-1+ | 38 |
| PD-1+ TOX1+ TIM3+ CD39+ EOMES+ CD38+ CD44+ TCF-1+ T-BET+ | 37 |
| PD-1+ TOX1+ LAG-3+ EOMES+ T-BET+ | 37 |
| TOX1+ LAG-3+ TCF-1+ T-BET+ | 37 |
| PD-1+ TOX1+ TIM3+ T-BET+ | 37 |
| TOX1+ TIM3+ CD38+ CD44+ | 37 |
| PD-1+ TOX1+ CD38+ CD44+ | 37 |
| PD-1+ TOX1+ LAG-3+ CD39+ EOMES+ TCF-1+ | 36 |
| TOX1+ TIM3+ LAG-3+ EOMES+ CD38+ CD44+ TCF-1+ T-BET+ | 36 |
| TOX1+ CD39+ EOMES+ T-BET+ | 35 |
| TOX1+ TIM3+ EOMES+ CD44+ | 35 |
| PD-1+ TOX1+ TIM3+ CD38+ CD44+ TCF-1+ T-BET+ | 34 |
| TIM3+ CD39+ CD44+ | 34 |
| LAG-3+ EOMES+ CD44+ TCF-1+ T-BET+ | 34 |
| TOX1+ TIM3+ CD38+ | 34 |
| TOX1+ CD38+ TCF-1+ T-BET+ | 33 |
| PD-1+ LAG-3+ EOMES+ TCF-1+ | 33 |
| PD-1+ TOX1+ TIM3+ CD39+ EOMES+ CD38+ CD44+ TCF-1+ | 32 |
| PD-1+ TOX1+ TIM3+ CD38+ CD44+ T-BET+ | 32 |
| LAG-3+ CD44+ T-BET+ | 32 |
| TOX1+ CD39+ EOMES+ CD44+ TCF-1+ | 32 |
| TIM3+ CD44+ TCF-1+ T-BET+ | 32 |
| TIM3+ CD39+ CD38+ | 32 |
| CD39+ CD44+ TCF-1+ T-BET+ | 32 |
| TIM3+ CD38+ CD44+ | 31 |
| LAG-3+ EOMES+ CD38+ CD44+ | 31 |
| PD-1+ TOX1+ TIM3+ LAG-3+ CD39+ CD44+ TCF-1+ T-BET+ | 31 |
| PD-1+ LAG-3+ EOMES+ | 31 |
| PD-1+ TOX1+ EOMES+ CD44+ T-BET+ | 31 |
| TOX1+ TIM3+ LAG-3+ CD39+ CD38+ CD44+ TCF-1+ T-BET+ | 31 |
| LAG-3+ CD38+ TCF-1+ | 31 |
| PD-1+ TOX1+ TIM3+ LAG-3+ EOMES+ CD38+ TCF-1+ T-BET+ | 30 |
| TOX1+ CD39+ CD38+ | 30 |
| TOX1+ TIM3+ CD44+ T-BET+ | 30 |
| TIM3+ EOMES+ TCF-1+ T-BET+ | 30 |
| TOX1+ LAG-3+ CD39+ EOMES+ CD44+ TCF-1+ T-BET+ | 30 |
| PD-1+ LAG-3+ | 30 |
| PD-1+ TOX1+ LAG-3+ CD39+ EOMES+ TCF-1+ T-BET+ | 30 |
| TOX1+ CD39+ CD44+ TCF-1+ | 29 |
| PD-1+ TOX1+ CD39+ T-BET+ | 29 |
| TOX1+ TIM3+ CD44+ TCF-1+ T-BET+ | 29 |
| PD-1+ TOX1+ TIM3+ LAG-3+ CD39+ EOMES+ CD38+ CD44+ T-BET+ | 29 |
| LAG-3+ CD39+ EOMES+ CD38+ CD44+ TCF-1+ | 29 |
| LAG-3+ CD39+ EOMES+ CD38+ CD44+ | 29 |
| TIM3+ LAG-3+ | 29 |
| LAG-3+ CD38+ | 28 |
| TOX1+ EOMES+ CD38+ CD44+ TCF-1+ T-BET+ | 28 |
| LAG-3+ CD39+ EOMES+ TCF-1+ T-BET+ | 28 |
| TIM3+ LAG-3+ T-BET+ | 28 |
| TOX1+ LAG-3+ EOMES+ TCF-1+ T-BET+ | 28 |
| TOX1+ CD39+ EOMES+ TCF-1+ T-BET+ | 28 |
| TOX1+ TIM3+ EOMES+ CD44+ TCF-1+ | 28 |
| LAG-3+ CD39+ EOMES+ CD38+ | 28 |
| PD-1+ TOX1+ LAG-3+ EOMES+ CD38+ TCF-1+ | 27 |
| TIM3+ EOMES+ CD44+ TCF-1+ T-BET+ | 27 |
| PD-1+ TOX1+ LAG-3+ CD39+ TCF-1+ | 27 |
| PD-1+ LAG-3+ TCF-1+ | 27 |
| PD-1+ LAG-3+ CD38+ TCF-1+ T-BET+ | 27 |
| TOX1+ LAG-3+ CD39+ TCF-1+ | 27 |
| PD-1+ TOX1+ LAG-3+ CD39+ T-BET+ | 27 |
| TOX1+ CD38+ CD44+ T-BET+ | 26 |
| TOX1+ TIM3+ LAG-3+ CD39+ EOMES+ CD38+ CD44+ TCF-1+ | 26 |
| PD-1+ TOX1+ LAG-3+ EOMES+ CD38+ | 26 |
| LAG-3+ EOMES+ CD38+ TCF-1+ | 26 |
| TOX1+ LAG-3+ CD39+ EOMES+ CD38+ TCF-1+ | 26 |
| LAG-3+ CD39+ EOMES+ CD38+ TCF-1+ | 26 |
| TOX1+ TIM3+ LAG-3+ CD38+ CD44+ TCF-1+ T-BET+ | 26 |
| PD-1+ TOX1+ TIM3+ CD39+ EOMES+ CD44+ TCF-1+ | 26 |
| LAG-3+ CD39+ EOMES+ TCF-1+ | 26 |
| PD-1+ TIM3+ | 25 |
| CD39+ CD38+ CD44+ TCF-1+ | 25 |
| PD-1+ TOX1+ LAG-3+ TCF-1+ T-BET+ | 25 |
| PD-1+ TIM3+ LAG-3+ EOMES+ CD38+ CD44+ TCF-1+ T-BET+ | 25 |
| LAG-3+ CD38+ CD44+ | 25 |
| TOX1+ TIM3+ LAG-3+ | 25 |
| LAG-3+ CD39+ TCF-1+ T-BET+ | 25 |
| TOX1+ LAG-3+ EOMES+ T-BET+ | 25 |
| TOX1+ TIM3+ CD38+ CD44+ TCF-1+ T-BET+ | 25 |
| TOX1+ TIM3+ LAG-3+ EOMES+ CD38+ TCF-1+ T-BET+ | 24 |
| PD-1+ TOX1+ CD39+ EOMES+ CD38+ | 24 |
| CD39+ CD38+ TCF-1+ | 24 |
| TIM3+ EOMES+ CD38+ CD44+ TCF-1+ | 24 |
| TOX1+ TIM3+ EOMES+ CD38+ CD44+ TCF-1+ | 23 |
| PD-1+ TOX1+ TIM3+ CD44+ TCF-1+ | 23 |
| TOX1+ LAG-3+ CD39+ EOMES+ CD38+ | 23 |
| PD-1+ TOX1+ CD39+ CD44+ | 23 |
| PD-1+ TOX1+ CD39+ EOMES+ CD38+ TCF-1+ | 23 |
| PD-1+ TOX1+ TIM3+ EOMES+ CD38+ CD44+ TCF-1+ T-BET+ | 23 |
| CD39+ CD44+ T-BET+ | 23 |
| PD-1+ TOX1+ CD39+ EOMES+ CD44+ TCF-1+ | 23 |
| TIM3+ LAG-3+ CD39+ EOMES+ CD38+ CD44+ TCF-1+ T-BET+ | 22 |
| LAG-3+ CD38+ CD44+ TCF-1+ T-BET+ | 22 |
| TOX1+ TIM3+ EOMES+ T-BET+ | 22 |
| TOX1+ CD39+ EOMES+ CD44+ | 22 |
| PD-1+ TOX1+ LAG-3+ CD39+ EOMES+ CD38+ TCF-1+ T-BET+ | 22 |
| EOMES+ CD38+ TCF-1+ T-BET+ | 22 |
| LAG-3+ CD38+ TCF-1+ T-BET+ | 22 |
| TIM3+ CD39+ T-BET+ | 22 |
| TOX1+ LAG-3+ CD39+ T-BET+ | 22 |
| LAG-3+ CD39+ EOMES+ CD44+ TCF-1+ T-BET+ | 22 |
| PD-1+ TOX1+ LAG-3+ EOMES+ CD38+ CD44+ TCF-1+ | 22 |
| PD-1+ LAG-3+ EOMES+ CD44+ TCF-1+ | 21 |
| PD-1+ TOX1+ CD39+ CD44+ TCF-1+ | 21 |
| PD-1+ EOMES+ CD44+ T-BET+ | 21 |
| TIM3+ LAG-3+ CD38+ CD44+ TCF-1+ T-BET+ | 21 |
| TIM3+ LAG-3+ EOMES+ TCF-1+ | 21 |
| PD-1+ TOX1+ LAG-3+ EOMES+ CD38+ CD44+ | 21 |
| TOX1+ TIM3+ EOMES+ TCF-1+ T-BET+ | 21 |
| PD-1+ TOX1+ LAG-3+ CD39+ EOMES+ T-BET+ | 21 |
| TOX1+ EOMES+ CD38+ T-BET+ | 21 |
| TOX1+ LAG-3+ EOMES+ CD38+ TCF-1+ | 20 |
| TIM3+ LAG-3+ TCF-1+ | 20 |
| TOX1+ LAG-3+ CD39+ EOMES+ CD38+ CD44+ | 20 |
| TOX1+ TIM3+ CD38+ TCF-1+ T-BET+ | 20 |
| TOX1+ TIM3+ EOMES+ CD38+ CD44+ TCF-1+ T-BET+ | 20 |
| PD-1+ TOX1+ TIM3+ CD44+ | 20 |
| TOX1+ TIM3+ CD39+ TCF-1+ | 20 |
| PD-1+ TOX1+ EOMES+ CD38+ | 20 |
| EOMES+ CD38+ CD44+ T-BET+ | 20 |
| PD-1+ TOX1+ CD38+ CD44+ TCF-1+ | 20 |
| PD-1+ TOX1+ CD39+ EOMES+ T-BET+ | 19 |
| TOX1+ CD39+ CD44+ T-BET+ | 19 |
| TIM3+ LAG-3+ EOMES+ CD38+ CD44+ TCF-1+ | 19 |
| LAG-3+ CD39+ T-BET+ | 19 |
| TOX1+ TIM3+ LAG-3+ EOMES+ CD38+ TCF-1+ | 19 |
| LAG-3+ EOMES+ CD38+ | 19 |
| PD-1+ TOX1+ TIM3+ LAG-3+ CD39+ CD38+ CD44+ T-BET+ | 19 |
| TOX1+ TIM3+ LAG-3+ EOMES+ CD38+ CD44+ TCF-1+ | 19 |
| PD-1+ CD39+ EOMES+ | 18 |
| CD39+ CD38+ CD44+ TCF-1+ T-BET+ | 18 |
| TIM3+ CD39+ CD38+ CD44+ | 18 |
| TIM3+ LAG-3+ EOMES+ CD38+ CD44+ TCF-1+ T-BET+ | 18 |
| PD-1+ TOX1+ CD38+ TCF-1+ | 18 |
| PD-1+ TOX1+ LAG-3+ EOMES+ TCF-1+ T-BET+ | 18 |
| TIM3+ CD38+ | 18 |
| TOX1+ LAG-3+ CD44+ TCF-1+ | 18 |
| LAG-3+ CD39+ CD44+ TCF-1+ | 17 |
| PD-1+ CD39+ TCF-1+ | 17 |
| CD39+ CD38+ T-BET+ | 17 |
| PD-1+ TOX1+ TIM3+ EOMES+ TCF-1+ | 17 |
| TIM3+ CD38+ CD44+ TCF-1+ T-BET+ | 17 |
| PD-1+ TOX1+ CD38+ T-BET+ | 17 |
| PD-1+ TOX1+ EOMES+ CD38+ CD44+ | 17 |
| PD-1+ TOX1+ CD39+ EOMES+ CD38+ CD44+ | 17 |
| PD-1+ TOX1+ TIM3+ EOMES+ | 17 |
| PD-1+ TOX1+ TIM3+ LAG-3+ CD39+ EOMES+ CD44+ TCF-1+ | 17 |
| PD-1+ TOX1+ TIM3+ CD39+ CD38+ CD44+ TCF-1+ T-BET+ | 17 |
| PD-1+ TOX1+ TIM3+ LAG-3+ CD44+ TCF-1+ T-BET+ | 17 |
| TIM3+ EOMES+ CD44+ T-BET+ | 17 |
| TOX1+ LAG-3+ CD39+ EOMES+ CD38+ TCF-1+ T-BET+ | 17 |
| LAG-3+ CD39+ CD38+ | 17 |
| PD-1+ TOX1+ TIM3+ CD38+ CD44+ | 16 |
| PD-1+ TOX1+ TIM3+ LAG-3+ CD39+ EOMES+ CD38+ TCF-1+ T-BET+ | 16 |
| LAG-3+ EOMES+ CD38+ TCF-1+ T-BET+ | 16 |
| PD-1+ LAG-3+ T-BET+ | 16 |
| CD39+ CD38+ CD44+ T-BET+ | 16 |
| PD-1+ LAG-3+ TCF-1+ T-BET+ | 16 |
| TOX1+ LAG-3+ EOMES+ CD38+ | 16 |
| TOX1+ TIM3+ EOMES+ CD38+ TCF-1+ | 16 |
| TOX1+ LAG-3+ CD39+ EOMES+ TCF-1+ T-BET+ | 16 |
| PD-1+ TOX1+ LAG-3+ EOMES+ CD44+ | 15 |
| TOX1+ LAG-3+ CD39+ EOMES+ CD44+ TCF-1+ | 15 |
| PD-1+ TOX1+ LAG-3+ CD44+ | 15 |
| PD-1+ LAG-3+ CD38+ TCF-1+ | 15 |
| PD-1+ TOX1+ CD39+ EOMES+ CD44+ TCF-1+ T-BET+ | 14 |
| PD-1+ TOX1+ EOMES+ CD38+ CD44+ TCF-1+ | 14 |
| LAG-3+ CD39+ CD38+ CD44+ TCF-1+ | 14 |
| LAG-3+ EOMES+ CD38+ CD44+ TCF-1+ T-BET+ | 14 |
| LAG-3+ CD39+ CD38+ T-BET+ | 14 |
| EOMES+ CD38+ T-BET+ | 14 |
| TOX1+ LAG-3+ CD44+ | 14 |
| PD-1+ TOX1+ LAG-3+ CD39+ CD44+ TCF-1+ T-BET+ | 14 |
| TOX1+ TIM3+ EOMES+ CD44+ TCF-1+ T-BET+ | 14 |
| PD-1+ TOX1+ TIM3+ CD39+ CD38+ CD44+ T-BET+ | 14 |
| PD-1+ LAG-3+ CD39+ | 14 |
| PD-1+ TOX1+ EOMES+ CD38+ CD44+ TCF-1+ T-BET+ | 14 |
| PD-1+ LAG-3+ CD39+ T-BET+ | 14 |
| TOX1+ LAG-3+ EOMES+ CD38+ CD44+ TCF-1+ | 14 |
| TOX1+ EOMES+ CD38+ TCF-1+ T-BET+ | 14 |
| TIM3+ LAG-3+ EOMES+ T-BET+ | 14 |
| TIM3+ CD39+ EOMES+ | 14 |
| PD-1+ TIM3+ T-BET+ | 14 |
| PD-1+ CD39+ TCF-1+ T-BET+ | 14 |
| PD-1+ TOX1+ LAG-3+ CD39+ TCF-1+ T-BET+ | 13 |
| TOX1+ LAG-3+ CD38+ TCF-1+ T-BET+ | 13 |
| PD-1+ TOX1+ CD38+ TCF-1+ T-BET+ | 13 |
| TIM3+ LAG-3+ CD38+ TCF-1+ T-BET+ | 13 |
| LAG-3+ CD39+ CD38+ TCF-1+ T-BET+ | 13 |
| LAG-3+ CD39+ EOMES+ CD38+ TCF-1+ T-BET+ | 13 |
| TOX1+ LAG-3+ EOMES+ CD38+ TCF-1+ T-BET+ | 13 |
| TOX1+ LAG-3+ CD39+ EOMES+ CD38+ CD44+ TCF-1+ | 13 |
| PD-1+ TOX1+ CD39+ EOMES+ CD44+ | 13 |
| PD-1+ TIM3+ LAG-3+ CD38+ TCF-1+ T-BET+ | 13 |
| PD-1+ TOX1+ TIM3+ CD44+ TCF-1+ T-BET+ | 13 |
| TOX1+ CD39+ CD38+ CD44+ | 13 |
| TOX1+ LAG-3+ CD39+ CD38+ | 13 |
| TOX1+ TIM3+ CD39+ EOMES+ | 13 |
| TOX1+ CD39+ CD38+ T-BET+ | 13 |
| PD-1+ TOX1+ TIM3+ CD38+ TCF-1+ T-BET+ | 13 |
| PD-1+ TOX1+ TIM3+ EOMES+ TCF-1+ T-BET+ | 13 |
| LAG-3+ CD39+ CD38+ CD44+ | 12 |
| TIM3+ CD39+ TCF-1+ T-BET+ | 12 |
| PD-1+ TOX1+ LAG-3+ EOMES+ CD44+ TCF-1+ | 12 |
| PD-1+ TOX1+ LAG-3+ CD39+ CD38+ TCF-1+ | 12 |
| CD39+ CD38+ TCF-1+ T-BET+ | 12 |
| PD-1+ TOX1+ LAG-3+ CD44+ TCF-1+ | 12 |
| PD-1+ TOX1+ LAG-3+ CD44+ TCF-1+ T-BET+ | 12 |
| PD-1+ TOX1+ CD39+ CD38+ | 12 |
| PD-1+ TOX1+ EOMES+ CD38+ TCF-1+ | 12 |
| TOX1+ LAG-3+ EOMES+ CD44+ TCF-1+ | 12 |
| PD-1+ TOX1+ TIM3+ LAG-3+ EOMES+ TCF-1+ | 12 |
| PD-1+ TIM3+ TCF-1+ | 12 |
| TOX1+ LAG-3+ CD39+ CD38+ TCF-1+ T-BET+ | 12 |
| TOX1+ TIM3+ EOMES+ CD38+ TCF-1+ T-BET+ | 12 |
| PD-1+ LAG-3+ CD39+ EOMES+ TCF-1+ | 12 |
| LAG-3+ EOMES+ CD44+ T-BET+ | 12 |
| PD-1+ TIM3+ CD44+ | 12 |
| TOX1+ LAG-3+ CD38+ | 12 |
| TOX1+ LAG-3+ CD39+ EOMES+ CD38+ CD44+ TCF-1+ T-BET+ | 12 |
| TOX1+ LAG-3+ CD44+ T-BET+ | 12 |
| TIM3+ LAG-3+ CD39+ | 12 |
| TIM3+ LAG-3+ EOMES+ CD38+ TCF-1+ T-BET+ | 12 |
| TOX1+ LAG-3+ EOMES+ CD38+ CD44+ | 12 |
| LAG-3+ CD38+ T-BET+ | 12 |
| CD39+ EOMES+ CD38+ TCF-1+ T-BET+ | 12 |
| TIM3+ LAG-3+ CD38+ T-BET+ | 12 |
| PD-1+ CD39+ EOMES+ TCF-1+ T-BET+ | 12 |
| TIM3+ LAG-3+ CD39+ EOMES+ CD44+ TCF-1+ | 12 |
| PD-1+ CD39+ T-BET+ | 12 |
| PD-1+ TOX1+ LAG-3+ CD39+ EOMES+ CD44+ TCF-1+ | 12 |
| PD-1+ TOX1+ EOMES+ CD38+ TCF-1+ T-BET+ | 11 |
| PD-1+ LAG-3+ EOMES+ CD44+ TCF-1+ T-BET+ | 11 |
| PD-1+ LAG-3+ EOMES+ TCF-1+ T-BET+ | 11 |
| LAG-3+ CD39+ EOMES+ CD38+ CD44+ TCF-1+ T-BET+ | 11 |
| TOX1+ LAG-3+ CD39+ TCF-1+ T-BET+ | 11 |
| TOX1+ CD39+ EOMES+ CD44+ T-BET+ | 11 |
| PD-1+ LAG-3+ CD44+ TCF-1+ | 11 |
| CD39+ EOMES+ CD38+ CD44+ TCF-1+ T-BET+ | 11 |
| PD-1+ TOX1+ TIM3+ LAG-3+ CD38+ TCF-1+ T-BET+ | 11 |
| TIM3+ CD38+ T-BET+ | 11 |
| TOX1+ TIM3+ EOMES+ CD38+ CD44+ | 11 |
| TIM3+ LAG-3+ EOMES+ | 11 |
| TOX1+ TIM3+ EOMES+ CD44+ T-BET+ | 11 |
| TOX1+ TIM3+ LAG-3+ CD39+ EOMES+ CD44+ TCF-1+ T-BET+ | 11 |
| PD-1+ TOX1+ LAG-3+ CD39+ CD44+ TCF-1+ | 11 |
| TIM3+ LAG-3+ CD44+ | 11 |
| LAG-3+ CD39+ EOMES+ CD44+ TCF-1+ | 11 |
| LAG-3+ CD39+ EOMES+ CD44+ | 11 |
| PD-1+ TOX1+ TIM3+ CD38+ | 11 |
| TOX1+ TIM3+ CD38+ T-BET+ | 11 |
| PD-1+ TOX1+ LAG-3+ EOMES+ CD38+ CD44+ TCF-1+ T-BET+ | 11 |
| PD-1+ CD38+ | 11 |
| TOX1+ TIM3+ CD38+ TCF-1+ | 11 |
| TIM3+ CD38+ TCF-1+ T-BET+ | 10 |
| PD-1+ TOX1+ TIM3+ LAG-3+ EOMES+ CD44+ TCF-1+ T-BET+ | 10 |
| PD-1+ TOX1+ LAG-3+ EOMES+ CD38+ CD44+ T-BET+ | 10 |
| PD-1+ LAG-3+ CD39+ EOMES+ CD38+ CD44+ TCF-1+ T-BET+ | 10 |
| TIM3+ CD38+ CD44+ T-BET+ | 10 |
| TIM3+ LAG-3+ CD38+ CD44+ T-BET+ | 10 |
| PD-1+ TIM3+ LAG-3+ CD39+ CD38+ CD44+ T-BET+ | 10 |
| TIM3+ LAG-3+ EOMES+ TCF-1+ T-BET+ | 10 |
| PD-1+ TOX1+ CD39+ CD38+ CD44+ | 10 |
| PD-1+ LAG-3+ EOMES+ T-BET+ | 10 |
| PD-1+ TOX1+ CD39+ EOMES+ CD44+ T-BET+ | 10 |
| PD-1+ TOX1+ TIM3+ CD39+ | 10 |
| PD-1+ TOX1+ CD39+ EOMES+ CD38+ CD44+ TCF-1+ | 10 |
| PD-1+ TOX1+ TIM3+ LAG-3+ | 10 |
| PD-1+ TOX1+ TIM3+ LAG-3+ CD38+ CD44+ T-BET+ | 10 |
| TOX1+ LAG-3+ CD39+ EOMES+ T-BET+ | 10 |
| LAG-3+ CD39+ CD44+ | 10 |
| TIM3+ CD39+ CD44+ TCF-1+ | 10 |
| TOX1+ TIM3+ LAG-3+ EOMES+ TCF-1+ | 10 |
| TIM3+ EOMES+ CD38+ CD44+ TCF-1+ T-BET+ | 9 |
| PD-1+ TIM3+ EOMES+ CD38+ CD44+ TCF-1+ T-BET+ | 9 |
| TIM3+ CD39+ CD38+ TCF-1+ | 9 |
| PD-1+ CD39+ CD44+ TCF-1+ | 9 |
| PD-1+ CD38+ TCF-1+ | 9 |
| PD-1+ LAG-3+ CD38+ CD44+ TCF-1+ T-BET+ | 9 |
| PD-1+ TIM3+ CD44+ TCF-1+ T-BET+ | 9 |
| TOX1+ EOMES+ CD38+ CD44+ T-BET+ | 9 |
| PD-1+ TOX1+ LAG-3+ EOMES+ CD44+ TCF-1+ T-BET+ | 9 |
| PD-1+ TOX1+ TIM3+ CD39+ TCF-1+ T-BET+ | 9 |
| LAG-3+ CD39+ CD38+ CD44+ TCF-1+ T-BET+ | 9 |
| PD-1+ TOX1+ TIM3+ CD39+ EOMES+ TCF-1+ | 9 |
| TOX1+ TIM3+ CD39+ EOMES+ TCF-1+ | 9 |
| TOX1+ LAG-3+ CD38+ T-BET+ | 9 |
| PD-1+ TOX1+ LAG-3+ CD38+ | 9 |
| PD-1+ TOX1+ TIM3+ LAG-3+ EOMES+ | 9 |
| TOX1+ TIM3+ CD39+ CD38+ TCF-1+ T-BET+ | 9 |
| TOX1+ TIM3+ CD38+ CD44+ TCF-1+ | 9 |
| PD-1+ TOX1+ TIM3+ LAG-3+ CD39+ EOMES+ CD38+ | 9 |
| TOX1+ LAG-3+ CD39+ CD44+ TCF-1+ | 9 |
| TIM3+ CD38+ CD44+ TCF-1+ | 9 |
| PD-1+ LAG-3+ EOMES+ CD38+ CD44+ | 8 |
| TOX1+ LAG-3+ EOMES+ CD44+ | 8 |
| PD-1+ TOX1+ TIM3+ CD39+ CD44+ TCF-1+ T-BET+ | 8 |
| PD-1+ TOX1+ TIM3+ EOMES+ CD44+ TCF-1+ | 8 |
| PD-1+ LAG-3+ CD39+ EOMES+ TCF-1+ T-BET+ | 8 |
| PD-1+ TOX1+ TIM3+ EOMES+ CD44+ | 8 |
| PD-1+ TOX1+ TIM3+ CD39+ EOMES+ | 8 |
| PD-1+ LAG-3+ CD39+ CD38+ TCF-1+ T-BET+ | 8 |
| PD-1+ TOX1+ TIM3+ CD44+ T-BET+ | 8 |
| LAG-3+ EOMES+ CD38+ T-BET+ | 8 |
| PD-1+ TOX1+ TIM3+ LAG-3+ CD38+ CD44+ | 8 |
| PD-1+ TOX1+ TIM3+ LAG-3+ CD39+ CD44+ TCF-1+ | 8 |
| CD39+ EOMES+ CD38+ CD44+ T-BET+ | 8 |
| PD-1+ TOX1+ TIM3+ LAG-3+ CD39+ EOMES+ TCF-1+ | 8 |
| CD39+ EOMES+ CD38+ T-BET+ | 8 |
| CD39+ EOMES+ CD44+ T-BET+ | 8 |
| PD-1+ CD39+ EOMES+ CD44+ TCF-1+ | 8 |
| TOX1+ CD39+ CD38+ TCF-1+ | 8 |
| PD-1+ CD39+ EOMES+ CD44+ | 8 |
| PD-1+ TOX1+ TIM3+ LAG-3+ CD39+ EOMES+ CD38+ CD44+ | 8 |
| TOX1+ LAG-3+ CD39+ CD44+ TCF-1+ T-BET+ | 8 |
| PD-1+ CD39+ EOMES+ TCF-1+ | 8 |
| TOX1+ TIM3+ LAG-3+ TCF-1+ | 8 |
| PD-1+ TOX1+ CD39+ CD44+ T-BET+ | 8 |
| PD-1+ TOX1+ CD39+ EOMES+ CD38+ CD44+ T-BET+ | 8 |
| TOX1+ LAG-3+ CD39+ EOMES+ CD44+ | 8 |
| PD-1+ TOX1+ LAG-3+ EOMES+ CD44+ T-BET+ | 8 |
| PD-1+ TOX1+ CD39+ CD44+ TCF-1+ T-BET+ | 8 |
| TOX1+ TIM3+ LAG-3+ CD39+ EOMES+ CD38+ TCF-1+ T-BET+ | 8 |
| PD-1+ TOX1+ CD38+ CD44+ TCF-1+ T-BET+ | 8 |
| PD-1+ TOX1+ LAG-3+ CD39+ CD38+ CD44+ TCF-1+ T-BET+ | 8 |
| TIM3+ EOMES+ CD38+ CD44+ | 8 |
| TIM3+ LAG-3+ EOMES+ CD44+ TCF-1+ T-BET+ | 8 |
| PD-1+ TIM3+ EOMES+ | 8 |
| TOX1+ TIM3+ LAG-3+ CD39+ | 8 |
| TOX1+ TIM3+ LAG-3+ CD39+ CD38+ TCF-1+ T-BET+ | 8 |
| PD-1+ TOX1+ CD39+ EOMES+ CD38+ CD44+ TCF-1+ T-BET+ | 8 |
| PD-1+ LAG-3+ CD38+ | 7 |
| TIM3+ EOMES+ CD38+ | 7 |
| PD-1+ TOX1+ CD38+ CD44+ T-BET+ | 7 |
| PD-1+ TOX1+ TIM3+ CD38+ T-BET+ | 7 |
| PD-1+ TOX1+ TIM3+ LAG-3+ EOMES+ CD38+ CD44+ | 7 |
| PD-1+ TOX1+ TIM3+ LAG-3+ EOMES+ CD38+ TCF-1+ | 7 |
| PD-1+ TOX1+ LAG-3+ CD44+ T-BET+ | 7 |
| PD-1+ TOX1+ TIM3+ CD39+ EOMES+ CD44+ TCF-1+ T-BET+ | 7 |
| PD-1+ TOX1+ LAG-3+ CD39+ CD38+ | 7 |
| PD-1+ TIM3+ LAG-3+ CD39+ EOMES+ CD38+ CD44+ TCF-1+ | 7 |
| LAG-3+ CD39+ CD38+ TCF-1+ | 7 |
| PD-1+ TOX1+ LAG-3+ CD39+ EOMES+ CD44+ | 7 |
| PD-1+ TOX1+ LAG-3+ CD39+ EOMES+ CD44+ TCF-1+ T-BET+ | 7 |
| PD-1+ TOX1+ LAG-3+ CD39+ EOMES+ CD38+ CD44+ T-BET+ | 7 |
| PD-1+ TOX1+ EOMES+ CD38+ T-BET+ | 7 |
| TOX1+ TIM3+ LAG-3+ CD39+ CD38+ T-BET+ | 7 |
| TOX1+ LAG-3+ CD39+ CD38+ T-BET+ | 7 |
| TIM3+ LAG-3+ CD38+ CD44+ TCF-1+ | 7 |
| TOX1+ LAG-3+ EOMES+ CD38+ CD44+ TCF-1+ T-BET+ | 7 |
| TOX1+ LAG-3+ CD39+ CD44+ | 7 |
| TIM3+ LAG-3+ CD39+ CD38+ CD44+ TCF-1+ T-BET+ | 7 |
| TOX1+ TIM3+ LAG-3+ CD38+ TCF-1+ T-BET+ | 7 |
| TIM3+ LAG-3+ CD39+ EOMES+ TCF-1+ | 7 |
| TOX1+ CD39+ EOMES+ CD38+ TCF-1+ T-BET+ | 7 |
| PD-1+ CD38+ CD44+ TCF-1+ | 7 |
| PD-1+ CD38+ CD44+ | 7 |
| TOX1+ LAG-3+ CD39+ CD38+ CD44+ TCF-1+ | 7 |
| TOX1+ TIM3+ LAG-3+ EOMES+ TCF-1+ T-BET+ | 7 |
| TOX1+ LAG-3+ CD39+ CD38+ CD44+ TCF-1+ T-BET+ | 7 |
| TIM3+ LAG-3+ CD39+ EOMES+ CD44+ TCF-1+ T-BET+ | 7 |
| TIM3+ LAG-3+ TCF-1+ T-BET+ | 7 |
| TOX1+ TIM3+ CD39+ TCF-1+ T-BET+ | 7 |
| TOX1+ TIM3+ CD39+ T-BET+ | 7 |
| TIM3+ LAG-3+ EOMES+ CD38+ TCF-1+ | 7 |
| LAG-3+ EOMES+ CD38+ CD44+ T-BET+ | 6 |
| TOX1+ TIM3+ LAG-3+ CD39+ CD38+ CD44+ T-BET+ | 6 |
| PD-1+ TOX1+ LAG-3+ CD39+ EOMES+ CD38+ T-BET+ | 6 |
| PD-1+ TOX1+ TIM3+ LAG-3+ CD39+ EOMES+ CD38+ T-BET+ | 6 |
| LAG-3+ CD39+ EOMES+ T-BET+ | 6 |
| PD-1+ TOX1+ TIM3+ CD39+ EOMES+ TCF-1+ T-BET+ | 6 |
| TIM3+ LAG-3+ CD39+ EOMES+ | 6 |
| PD-1+ TOX1+ CD39+ EOMES+ CD38+ TCF-1+ T-BET+ | 6 |
| PD-1+ LAG-3+ EOMES+ CD38+ TCF-1+ T-BET+ | 6 |
| PD-1+ TIM3+ LAG-3+ CD39+ EOMES+ TCF-1+ T-BET+ | 6 |
| TOX1+ TIM3+ LAG-3+ CD44+ TCF-1+ T-BET+ | 6 |
| TOX1+ TIM3+ LAG-3+ CD39+ EOMES+ | 6 |
| TOX1+ LAG-3+ CD44+ TCF-1+ T-BET+ | 6 |
| LAG-3+ CD39+ EOMES+ CD38+ T-BET+ | 6 |
| PD-1+ TOX1+ TIM3+ LAG-3+ TCF-1+ T-BET+ | 6 |
| TIM3+ LAG-3+ CD39+ CD38+ | 6 |
| TOX1+ TIM3+ CD39+ CD38+ CD44+ | 6 |
| PD-1+ TIM3+ CD44+ TCF-1+ | 6 |
| TOX1+ LAG-3+ CD39+ CD44+ T-BET+ | 6 |
| PD-1+ CD39+ CD38+ T-BET+ | 6 |
| TOX1+ CD39+ CD38+ CD44+ TCF-1+ | 6 |
| TIM3+ CD39+ CD44+ TCF-1+ T-BET+ | 6 |
| TIM3+ EOMES+ CD38+ TCF-1+ T-BET+ | 6 |
| PD-1+ TOX1+ TIM3+ CD39+ T-BET+ | 6 |
| TOX1+ LAG-3+ EOMES+ CD38+ T-BET+ | 6 |
| PD-1+ TOX1+ TIM3+ CD39+ TCF-1+ | 6 |
| PD-1+ LAG-3+ CD44+ | 6 |
| PD-1+ LAG-3+ CD44+ T-BET+ | 6 |
| LAG-3+ CD39+ CD44+ TCF-1+ T-BET+ | 6 |
| PD-1+ TIM3+ EOMES+ CD44+ TCF-1+ | 6 |
| TOX1+ TIM3+ CD39+ CD44+ | 6 |
| TIM3+ CD39+ CD38+ T-BET+ | 6 |
| TOX1+ CD39+ CD38+ CD44+ TCF-1+ T-BET+ | 6 |
| PD-1+ TOX1+ LAG-3+ CD38+ TCF-1+ T-BET+ | 6 |
| PD-1+ TOX1+ TIM3+ LAG-3+ CD39+ TCF-1+ T-BET+ | 6 |
| TIM3+ CD39+ CD38+ TCF-1+ T-BET+ | 6 |
| PD-1+ CD38+ TCF-1+ T-BET+ | 6 |
| TOX1+ CD39+ EOMES+ CD38+ T-BET+ | 6 |
| PD-1+ LAG-3+ CD39+ EOMES+ T-BET+ | 6 |
| TIM3+ LAG-3+ CD39+ CD38+ T-BET+ | 6 |
| PD-1+ TOX1+ LAG-3+ EOMES+ CD38+ T-BET+ | 5 |
| PD-1+ TIM3+ LAG-3+ EOMES+ CD38+ TCF-1+ | 5 |
| TOX1+ TIM3+ CD39+ CD38+ CD44+ T-BET+ | 5 |
| TOX1+ TIM3+ LAG-3+ EOMES+ CD38+ T-BET+ | 5 |
| PD-1+ TOX1+ LAG-3+ EOMES+ CD38+ TCF-1+ T-BET+ | 5 |
| TOX1+ TIM3+ CD39+ CD38+ T-BET+ | 5 |
| TOX1+ TIM3+ CD39+ CD38+ | 5 |
| TOX1+ TIM3+ LAG-3+ CD39+ CD38+ | 5 |
| PD-1+ TIM3+ EOMES+ TCF-1+ T-BET+ | 5 |
| PD-1+ TIM3+ EOMES+ CD44+ TCF-1+ T-BET+ | 5 |
| PD-1+ TOX1+ TIM3+ CD38+ CD44+ TCF-1+ | 5 |
| TOX1+ LAG-3+ CD39+ CD38+ TCF-1+ | 5 |
| TOX1+ LAG-3+ CD39+ EOMES+ TCF-1+ | 5 |
| PD-1+ TOX1+ LAG-3+ CD39+ CD44+ | 5 |
| PD-1+ TOX1+ LAG-3+ CD39+ CD38+ T-BET+ | 5 |
| PD-1+ TOX1+ LAG-3+ CD39+ CD38+ TCF-1+ T-BET+ | 5 |
| TOX1+ TIM3+ CD39+ CD44+ TCF-1+ | 5 |
| TOX1+ LAG-3+ CD38+ CD44+ | 5 |
| TOX1+ TIM3+ LAG-3+ CD38+ TCF-1+ | 5 |
| PD-1+ LAG-3+ CD39+ TCF-1+ T-BET+ | 5 |
| TOX1+ CD39+ EOMES+ CD38+ CD44+ TCF-1+ T-BET+ | 5 |
| PD-1+ LAG-3+ CD39+ EOMES+ CD44+ TCF-1+ | 5 |
| PD-1+ LAG-3+ EOMES+ CD44+ | 5 |
| PD-1+ EOMES+ CD38+ TCF-1+ | 5 |
| TIM3+ LAG-3+ CD38+ TCF-1+ | 5 |
| PD-1+ TOX1+ TIM3+ LAG-3+ CD39+ CD44+ | 5 |
| TOX1+ TIM3+ LAG-3+ CD38+ CD44+ TCF-1+ | 5 |
| TOX1+ TIM3+ LAG-3+ EOMES+ | 5 |
| TIM3+ CD39+ EOMES+ TCF-1+ T-BET+ | 5 |
| TOX1+ TIM3+ LAG-3+ EOMES+ CD44+ TCF-1+ T-BET+ | 5 |
| TIM3+ EOMES+ CD38+ CD44+ T-BET+ | 5 |
| PD-1+ TOX1+ TIM3+ LAG-3+ CD39+ EOMES+ | 5 |
| PD-1+ TOX1+ TIM3+ LAG-3+ CD39+ EOMES+ TCF-1+ T-BET+ | 5 |
| PD-1+ CD39+ CD38+ TCF-1+ | 5 |
| TIM3+ LAG-3+ CD39+ EOMES+ CD38+ CD44+ TCF-1+ | 5 |
| PD-1+ LAG-3+ CD39+ TCF-1+ | 5 |
| PD-1+ TOX1+ LAG-3+ CD39+ CD38+ CD44+ T-BET+ | 5 |
| PD-1+ LAG-3+ CD39+ EOMES+ | 5 |
| PD-1+ TOX1+ LAG-3+ CD38+ CD44+ | 5 |
| TOX1+ TIM3+ CD39+ EOMES+ CD44+ TCF-1+ | 5 |
| TOX1+ TIM3+ LAG-3+ TCF-1+ T-BET+ | 5 |
| PD-1+ TOX1+ LAG-3+ CD38+ CD44+ TCF-1+ T-BET+ | 5 |
| PD-1+ TIM3+ LAG-3+ CD39+ CD38+ T-BET+ | 5 |
| PD-1+ TOX1+ LAG-3+ CD38+ T-BET+ | 5 |
| TIM3+ LAG-3+ CD38+ | 5 |
| PD-1+ LAG-3+ CD39+ CD44+ TCF-1+ | 5 |
| TOX1+ LAG-3+ CD38+ TCF-1+ | 5 |
| TIM3+ LAG-3+ CD39+ CD38+ TCF-1+ T-BET+ | 5 |
| PD-1+ TIM3+ LAG-3+ EOMES+ CD38+ CD44+ TCF-1+ | 4 |
| PD-1+ TIM3+ LAG-3+ T-BET+ | 4 |
| TOX1+ TIM3+ LAG-3+ CD39+ CD38+ CD44+ | 4 |
| TIM3+ EOMES+ CD38+ TCF-1+ | 4 |
| TOX1+ TIM3+ CD39+ EOMES+ TCF-1+ T-BET+ | 4 |
| TIM3+ EOMES+ CD38+ T-BET+ | 4 |
| TOX1+ TIM3+ CD39+ CD38+ CD44+ TCF-1+ | 4 |
| PD-1+ TIM3+ LAG-3+ CD38+ CD44+ TCF-1+ T-BET+ | 4 |
| TOX1+ TIM3+ CD39+ EOMES+ CD44+ TCF-1+ T-BET+ | 4 |
| PD-1+ TOX1+ CD39+ CD38+ T-BET+ | 4 |
| TOX1+ TIM3+ LAG-3+ CD39+ EOMES+ CD38+ CD44+ | 4 |
| TOX1+ TIM3+ LAG-3+ CD39+ EOMES+ CD38+ CD44+ T-BET+ | 4 |
| PD-1+ TOX1+ LAG-3+ CD39+ CD38+ CD44+ TCF-1+ | 4 |
| PD-1+ TIM3+ LAG-3+ CD39+ EOMES+ CD38+ CD44+ T-BET+ | 4 |
| PD-1+ TIM3+ LAG-3+ EOMES+ CD38+ TCF-1+ T-BET+ | 4 |
| PD-1+ TIM3+ EOMES+ TCF-1+ | 4 |
| TIM3+ LAG-3+ EOMES+ CD44+ T-BET+ | 4 |
| PD-1+ EOMES+ CD38+ CD44+ TCF-1+ T-BET+ | 4 |
| PD-1+ TOX1+ TIM3+ LAG-3+ CD39+ CD38+ TCF-1+ T-BET+ | 4 |
| PD-1+ TOX1+ TIM3+ CD39+ CD38+ CD44+ | 4 |
| TOX1+ LAG-3+ CD38+ CD44+ TCF-1+ | 4 |
| TOX1+ LAG-3+ CD38+ CD44+ T-BET+ | 4 |
| PD-1+ CD39+ CD44+ | 4 |
| PD-1+ LAG-3+ CD39+ CD44+ TCF-1+ T-BET+ | 4 |
| TIM3+ CD39+ EOMES+ CD44+ TCF-1+ | 4 |
| PD-1+ LAG-3+ CD44+ TCF-1+ T-BET+ | 4 |
| PD-1+ TIM3+ TCF-1+ T-BET+ | 4 |
| PD-1+ EOMES+ CD38+ CD44+ TCF-1+ | 4 |
| TIM3+ LAG-3+ CD39+ CD38+ CD44+ T-BET+ | 4 |
| PD-1+ LAG-3+ CD38+ T-BET+ | 4 |
| PD-1+ TOX1+ TIM3+ LAG-3+ CD39+ CD44+ T-BET+ | 4 |
| PD-1+ TOX1+ TIM3+ LAG-3+ CD38+ CD44+ TCF-1+ | 4 |
| PD-1+ TOX1+ TIM3+ LAG-3+ CD39+ TCF-1+ | 4 |
| PD-1+ TOX1+ TIM3+ LAG-3+ EOMES+ TCF-1+ T-BET+ | 4 |
| PD-1+ LAG-3+ CD39+ EOMES+ CD38+ CD44+ TCF-1+ | 4 |
| PD-1+ LAG-3+ CD39+ EOMES+ CD44+ TCF-1+ T-BET+ | 4 |
| PD-1+ TOX1+ TIM3+ EOMES+ T-BET+ | 4 |
| TOX1+ LAG-3+ CD39+ CD38+ CD44+ | 4 |
| PD-1+ TIM3+ CD44+ T-BET+ | 4 |
| PD-1+ TOX1+ TIM3+ EOMES+ CD44+ TCF-1+ T-BET+ | 4 |
| TOX1+ TIM3+ CD38+ CD44+ T-BET+ | 4 |
| TOX1+ LAG-3+ EOMES+ CD38+ CD44+ T-BET+ | 4 |
| PD-1+ TOX1+ TIM3+ LAG-3+ CD39+ EOMES+ CD44+ T-BET+ | 4 |
| PD-1+ TOX1+ TIM3+ EOMES+ CD44+ T-BET+ | 4 |
| PD-1+ TIM3+ CD39+ TCF-1+ | 4 |
| PD-1+ TOX1+ TIM3+ EOMES+ CD38+ CD44+ TCF-1+ | 4 |
| PD-1+ TIM3+ EOMES+ CD44+ T-BET+ | 4 |
| TIM3+ LAG-3+ CD39+ EOMES+ CD38+ CD44+ | 3 |
| TOX1+ CD39+ CD38+ TCF-1+ T-BET+ | 3 |
| TIM3+ LAG-3+ CD39+ EOMES+ CD38+ TCF-1+ | 3 |
| PD-1+ TOX1+ TIM3+ LAG-3+ CD39+ CD38+ CD44+ TCF-1+ | 3 |
| TOX1+ TIM3+ LAG-3+ CD38+ CD44+ T-BET+ | 3 |
| TOX1+ TIM3+ LAG-3+ CD39+ TCF-1+ T-BET+ | 3 |
| PD-1+ TOX1+ EOMES+ CD38+ CD44+ T-BET+ | 3 |
| TOX1+ TIM3+ LAG-3+ EOMES+ CD44+ | 3 |
| TIM3+ CD38+ TCF-1+ | 3 |
| TIM3+ LAG-3+ CD39+ EOMES+ CD38+ | 3 |
| TOX1+ TIM3+ LAG-3+ EOMES+ CD44+ TCF-1+ | 3 |
| TIM3+ LAG-3+ CD39+ EOMES+ CD44+ T-BET+ | 3 |
| PD-1+ TOX1+ TIM3+ LAG-3+ CD39+ CD38+ T-BET+ | 3 |
| TOX1+ TIM3+ LAG-3+ CD38+ CD44+ | 3 |
| PD-1+ TOX1+ TIM3+ LAG-3+ CD39+ | 3 |
| TOX1+ TIM3+ LAG-3+ T-BET+ | 3 |
| PD-1+ TOX1+ TIM3+ LAG-3+ EOMES+ CD38+ CD44+ T-BET+ | 3 |
| TIM3+ LAG-3+ CD39+ EOMES+ TCF-1+ T-BET+ | 3 |
| TIM3+ LAG-3+ EOMES+ CD38+ T-BET+ | 3 |
| TIM3+ LAG-3+ CD39+ T-BET+ | 3 |
| TIM3+ LAG-3+ EOMES+ CD44+ | 3 |
| TIM3+ LAG-3+ CD39+ TCF-1+ | 3 |
| TIM3+ LAG-3+ CD39+ TCF-1+ T-BET+ | 3 |
| TOX1+ TIM3+ CD39+ CD44+ T-BET+ | 3 |
| TIM3+ LAG-3+ CD39+ CD44+ TCF-1+ T-BET+ | 3 |
| TOX1+ LAG-3+ EOMES+ CD44+ TCF-1+ T-BET+ | 3 |
| PD-1+ TOX1+ TIM3+ CD39+ CD44+ TCF-1+ | 3 |
| TIM3+ LAG-3+ CD39+ CD38+ CD44+ | 3 |
| TOX1+ LAG-3+ CD38+ CD44+ TCF-1+ T-BET+ | 3 |
| PD-1+ TOX1+ LAG-3+ CD38+ CD44+ TCF-1+ | 3 |
| PD-1+ TOX1+ TIM3+ CD39+ EOMES+ CD38+ TCF-1+ T-BET+ | 3 |
| PD-1+ TOX1+ TIM3+ CD39+ EOMES+ CD38+ CD44+ | 3 |
| LAG-3+ CD38+ CD44+ T-BET+ | 3 |
| TOX1+ TIM3+ CD39+ EOMES+ CD38+ TCF-1+ T-BET+ | 3 |
| TOX1+ TIM3+ CD39+ EOMES+ CD38+ CD44+ T-BET+ | 3 |
| TIM3+ LAG-3+ EOMES+ CD44+ TCF-1+ | 3 |
| PD-1+ TOX1+ TIM3+ LAG-3+ T-BET+ | 3 |
| PD-1+ TOX1+ TIM3+ LAG-3+ CD44+ TCF-1+ | 3 |
| LAG-3+ CD39+ EOMES+ CD38+ CD44+ T-BET+ | 3 |
| TIM3+ LAG-3+ CD39+ CD38+ CD44+ TCF-1+ | 3 |
| PD-1+ TOX1+ TIM3+ LAG-3+ EOMES+ T-BET+ | 3 |
| TOX1+ TIM3+ LAG-3+ CD39+ CD44+ T-BET+ | 3 |
| PD-1+ LAG-3+ CD39+ EOMES+ CD38+ T-BET+ | 3 |
| TIM3+ LAG-3+ CD44+ TCF-1+ | 3 |
| PD-1+ TIM3+ EOMES+ CD38+ TCF-1+ | 3 |
| TIM3+ CD39+ CD44+ T-BET+ | 3 |
| PD-1+ LAG-3+ CD39+ EOMES+ CD44+ | 3 |
| TIM3+ LAG-3+ CD44+ TCF-1+ T-BET+ | 3 |
| PD-1+ EOMES+ CD38+ | 3 |
| PD-1+ TIM3+ EOMES+ CD44+ | 3 |
| TIM3+ CD39+ EOMES+ CD44+ TCF-1+ T-BET+ | 3 |
| PD-1+ LAG-3+ EOMES+ CD38+ | 3 |
| PD-1+ LAG-3+ EOMES+ CD38+ TCF-1+ | 3 |
| PD-1+ TIM3+ EOMES+ T-BET+ | 3 |
| PD-1+ TIM3+ LAG-3+ CD39+ EOMES+ CD44+ TCF-1+ | 3 |
| PD-1+ LAG-3+ EOMES+ CD38+ CD44+ TCF-1+ | 3 |
| PD-1+ TIM3+ LAG-3+ CD39+ EOMES+ TCF-1+ | 3 |
| PD-1+ LAG-3+ EOMES+ CD38+ CD44+ TCF-1+ T-BET+ | 3 |
| PD-1+ CD38+ T-BET+ | 3 |
| TIM3+ CD39+ CD38+ CD44+ TCF-1+ T-BET+ | 3 |
| PD-1+ TIM3+ LAG-3+ EOMES+ TCF-1+ T-BET+ | 3 |
| PD-1+ LAG-3+ CD39+ EOMES+ CD38+ TCF-1+ | 3 |
| PD-1+ TIM3+ LAG-3+ CD39+ CD44+ TCF-1+ T-BET+ | 3 |
| TIM3+ CD39+ EOMES+ CD38+ CD44+ TCF-1+ | 3 |
| PD-1+ TIM3+ LAG-3+ EOMES+ CD44+ TCF-1+ T-BET+ | 3 |
| TIM3+ CD39+ EOMES+ CD38+ CD44+ TCF-1+ T-BET+ | 3 |
| PD-1+ TIM3+ LAG-3+ CD38+ TCF-1+ | 3 |
| TOX1+ TIM3+ LAG-3+ CD39+ EOMES+ CD38+ TCF-1+ | 3 |
| PD-1+ CD38+ CD44+ TCF-1+ T-BET+ | 3 |
| PD-1+ TIM3+ CD39+ EOMES+ CD44+ TCF-1+ | 3 |
| PD-1+ CD39+ CD38+ | 3 |
| PD-1+ CD39+ EOMES+ T-BET+ | 3 |
| PD-1+ CD39+ EOMES+ CD38+ TCF-1+ | 3 |
| PD-1+ CD39+ CD38+ TCF-1+ T-BET+ | 3 |
| TOX1+ TIM3+ LAG-3+ CD39+ EOMES+ TCF-1+ T-BET+ | 3 |
| PD-1+ CD39+ EOMES+ CD44+ TCF-1+ T-BET+ | 3 |
| PD-1+ CD39+ EOMES+ CD38+ CD44+ T-BET+ | 3 |
| PD-1+ TIM3+ CD39+ EOMES+ CD38+ CD44+ TCF-1+ T-BET+ | 3 |
| PD-1+ CD39+ EOMES+ CD38+ CD44+ TCF-1+ T-BET+ | 3 |
| PD-1+ TIM3+ CD39+ | 3 |
| PD-1+ TIM3+ CD39+ CD38+ CD44+ TCF-1+ T-BET+ | 3 |
| PD-1+ TIM3+ EOMES+ CD38+ CD44+ TCF-1+ | 3 |
| PD-1+ CD39+ CD44+ T-BET+ | 3 |
| PD-1+ LAG-3+ CD39+ EOMES+ CD38+ | 2 |
| PD-1+ TOX1+ TIM3+ CD39+ CD38+ CD44+ TCF-1+ | 2 |
| TOX1+ LAG-3+ EOMES+ CD44+ T-BET+ | 2 |
| PD-1+ LAG-3+ CD39+ EOMES+ CD38+ CD44+ T-BET+ | 2 |
| PD-1+ TIM3+ EOMES+ CD38+ CD44+ | 2 |
| PD-1+ TIM3+ CD39+ CD38+ T-BET+ | 2 |
| TIM3+ LAG-3+ EOMES+ CD38+ CD44+ T-BET+ | 2 |
| PD-1+ TOX1+ TIM3+ EOMES+ CD38+ CD44+ | 2 |
| PD-1+ TIM3+ EOMES+ CD38+ CD44+ T-BET+ | 2 |
| PD-1+ TIM3+ CD39+ CD44+ T-BET+ | 2 |
| TOX1+ LAG-3+ CD39+ EOMES+ CD38+ T-BET+ | 2 |
| TIM3+ LAG-3+ CD44+ T-BET+ | 2 |
| TIM3+ LAG-3+ CD39+ CD44+ TCF-1+ | 2 |
| TOX1+ TIM3+ LAG-3+ CD39+ CD44+ TCF-1+ | 2 |
| TOX1+ LAG-3+ CD39+ CD38+ CD44+ T-BET+ | 2 |
| PD-1+ TIM3+ CD38+ CD44+ TCF-1+ T-BET+ | 2 |
| PD-1+ TOX1+ TIM3+ LAG-3+ CD44+ | 2 |
| PD-1+ LAG-3+ CD39+ CD38+ CD44+ T-BET+ | 2 |
| PD-1+ TOX1+ TIM3+ CD39+ EOMES+ CD38+ TCF-1+ | 2 |
| PD-1+ CD39+ CD38+ CD44+ TCF-1+ T-BET+ | 2 |
| TIM3+ LAG-3+ CD39+ EOMES+ CD38+ TCF-1+ T-BET+ | 2 |
| PD-1+ TOX1+ TIM3+ LAG-3+ CD39+ EOMES+ T-BET+ | 2 |
| PD-1+ CD39+ EOMES+ CD38+ T-BET+ | 2 |
| TIM3+ LAG-3+ CD39+ EOMES+ CD38+ T-BET+ | 2 |
| PD-1+ TOX1+ TIM3+ LAG-3+ CD39+ CD38+ CD44+ | 2 |
| PD-1+ CD39+ EOMES+ CD38+ CD44+ TCF-1+ | 2 |
| PD-1+ TOX1+ TIM3+ LAG-3+ CD39+ CD38+ | 2 |
| TIM3+ CD39+ EOMES+ CD44+ | 2 |
| TIM3+ CD39+ EOMES+ TCF-1+ | 2 |
| PD-1+ LAG-3+ CD38+ CD44+ TCF-1+ | 2 |
| TIM3+ LAG-3+ CD39+ EOMES+ CD44+ | 2 |
| TIM3+ CD39+ EOMES+ T-BET+ | 2 |
| PD-1+ EOMES+ CD38+ TCF-1+ T-BET+ | 2 |
| TOX1+ CD39+ EOMES+ CD38+ CD44+ T-BET+ | 2 |
| PD-1+ TOX1+ TIM3+ LAG-3+ EOMES+ CD44+ TCF-1+ | 2 |
| PD-1+ TOX1+ TIM3+ LAG-3+ EOMES+ CD44+ | 2 |
| TIM3+ LAG-3+ CD39+ EOMES+ T-BET+ | 2 |
| TIM3+ CD39+ EOMES+ CD38+ T-BET+ | 2 |
| PD-1+ LAG-3+ EOMES+ CD38+ CD44+ T-BET+ | 2 |
| PD-1+ TOX1+ TIM3+ LAG-3+ CD38+ T-BET+ | 2 |
| PD-1+ TOX1+ TIM3+ LAG-3+ CD38+ | 2 |
| PD-1+ TOX1+ TIM3+ LAG-3+ CD44+ T-BET+ | 2 |
| PD-1+ TIM3+ CD39+ EOMES+ TCF-1+ T-BET+ | 2 |
| TIM3+ CD39+ EOMES+ CD38+ CD44+ | 2 |
| PD-1+ TOX1+ TIM3+ LAG-3+ TCF-1+ | 2 |
| PD-1+ LAG-3+ CD39+ CD38+ | 2 |
| PD-1+ LAG-3+ CD39+ CD38+ T-BET+ | 2 |
| PD-1+ TOX1+ TIM3+ CD39+ EOMES+ CD38+ CD44+ T-BET+ | 2 |
| PD-1+ TOX1+ LAG-3+ CD39+ EOMES+ CD44+ T-BET+ | 2 |
| PD-1+ CD39+ CD38+ CD44+ | 2 |
| TIM3+ LAG-3+ EOMES+ CD38+ | 2 |
| TOX1+ TIM3+ LAG-3+ CD39+ EOMES+ CD44+ TCF-1+ | 2 |
| PD-1+ CD39+ CD38+ CD44+ T-BET+ | 2 |
| TOX1+ TIM3+ LAG-3+ EOMES+ CD38+ | 2 |
| TOX1+ TIM3+ LAG-3+ CD39+ EOMES+ TCF-1+ | 2 |
| TOX1+ TIM3+ LAG-3+ CD39+ EOMES+ CD44+ | 2 |
| PD-1+ TOX1+ CD39+ CD38+ TCF-1+ | 2 |
| PD-1+ TOX1+ CD39+ CD38+ CD44+ T-BET+ | 2 |
| PD-1+ TIM3+ LAG-3+ EOMES+ CD44+ T-BET+ | 2 |
| PD-1+ TIM3+ LAG-3+ EOMES+ CD44+ TCF-1+ | 2 |
| TOX1+ TIM3+ CD39+ EOMES+ T-BET+ | 2 |
| PD-1+ TOX1+ CD39+ CD38+ CD44+ TCF-1+ T-BET+ | 2 |
| PD-1+ TIM3+ LAG-3+ CD39+ T-BET+ | 2 |
| TOX1+ TIM3+ LAG-3+ EOMES+ CD38+ CD44+ | 2 |
| TOX1+ TIM3+ LAG-3+ EOMES+ T-BET+ | 2 |
| TOX1+ TIM3+ LAG-3+ CD39+ EOMES+ CD38+ T-BET+ | 2 |
| TIM3+ CD39+ CD38+ CD44+ T-BET+ | 2 |
| PD-1+ TIM3+ LAG-3+ CD39+ CD44+ T-BET+ | 2 |
| PD-1+ TOX1+ LAG-3+ CD38+ CD44+ T-BET+ | 2 |
| PD-1+ TOX1+ LAG-3+ CD38+ TCF-1+ | 2 |
| PD-1+ TIM3+ LAG-3+ CD39+ EOMES+ CD38+ T-BET+ | 2 |
| PD-1+ TIM3+ LAG-3+ CD39+ CD38+ CD44+ TCF-1+ | 2 |
| TOX1+ TIM3+ LAG-3+ CD44+ TCF-1+ | 2 |
| PD-1+ TIM3+ LAG-3+ CD39+ EOMES+ T-BET+ | 2 |
| PD-1+ TIM3+ LAG-3+ EOMES+ | 2 |
| PD-1+ TIM3+ LAG-3+ EOMES+ T-BET+ | 2 |
| LAG-3+ CD39+ CD38+ CD44+ T-BET+ | 2 |
| TOX1+ TIM3+ EOMES+ CD38+ CD44+ T-BET+ | 2 |
| PD-1+ TIM3+ CD39+ EOMES+ CD44+ | 2 |
| TOX1+ TIM3+ EOMES+ CD38+ | 2 |
| TOX1+ TIM3+ EOMES+ CD38+ T-BET+ | 2 |
| TOX1+ TIM3+ LAG-3+ CD39+ CD38+ TCF-1+ | 2 |
| PD-1+ TIM3+ CD39+ EOMES+ CD38+ TCF-1+ T-BET+ | 2 |
| TIM3+ LAG-3+ CD38+ CD44+ | 2 |
| PD-1+ TIM3+ CD39+ EOMES+ CD38+ CD44+ T-BET+ | 2 |
| PD-1+ TOX1+ LAG-3+ CD39+ CD38+ CD44+ | 2 |
| TOX1+ TIM3+ LAG-3+ CD44+ | 2 |
| PD-1+ TIM3+ LAG-3+ TCF-1+ T-BET+ | 2 |
| PD-1+ TOX1+ LAG-3+ CD39+ CD44+ T-BET+ | 2 |
| PD-1+ TIM3+ LAG-3+ CD38+ T-BET+ | 2 |
| TOX1+ TIM3+ LAG-3+ EOMES+ CD38+ CD44+ T-BET+ | 2 |
| TOX1+ TIM3+ LAG-3+ CD39+ CD44+ | 1 |
| TOX1+ TIM3+ LAG-3+ CD39+ CD38+ CD44+ TCF-1+ | 1 |
| PD-1+ TIM3+ LAG-3+ CD39+ EOMES+ CD38+ CD44+ | 1 |
| PD-1+ CD39+ EOMES+ CD38+ CD44+ | 1 |
| PD-1+ TIM3+ LAG-3+ CD39+ EOMES+ CD38+ TCF-1+ | 1 |
| PD-1+ CD39+ EOMES+ CD38+ | 1 |
| TOX1+ TIM3+ LAG-3+ EOMES+ CD44+ T-BET+ | 1 |
| PD-1+ TOX1+ TIM3+ LAG-3+ EOMES+ CD38+ | 1 |
| PD-1+ LAG-3+ EOMES+ CD38+ T-BET+ | 1 |
| TIM3+ LAG-3+ CD39+ EOMES+ CD38+ CD44+ T-BET+ | 1 |
| TOX1+ TIM3+ LAG-3+ CD38+ | 1 |
| PD-1+ TIM3+ LAG-3+ CD39+ EOMES+ CD44+ TCF-1+ T-BET+ | 1 |
| PD-1+ CD39+ CD38+ CD44+ TCF-1+ | 1 |
| TOX1+ TIM3+ LAG-3+ CD44+ T-BET+ | 1 |
| TOX1+ TIM3+ LAG-3+ CD38+ T-BET+ | 1 |
| PD-1+ TOX1+ TIM3+ LAG-3+ EOMES+ CD38+ T-BET+ | 1 |
| PD-1+ TOX1+ TIM3+ LAG-3+ CD38+ TCF-1+ | 1 |
| TOX1+ TIM3+ LAG-3+ CD39+ T-BET+ | 1 |
| PD-1+ TOX1+ CD39+ CD38+ TCF-1+ T-BET+ | 1 |
| PD-1+ EOMES+ CD38+ CD44+ T-BET+ | 1 |
| PD-1+ TOX1+ CD39+ CD38+ CD44+ TCF-1+ | 1 |
| PD-1+ LAG-3+ CD38+ CD44+ | 1 |
| PD-1+ TOX1+ TIM3+ LAG-3+ CD39+ EOMES+ CD44+ | 1 |
| PD-1+ LAG-3+ CD38+ CD44+ T-BET+ | 1 |
| PD-1+ TOX1+ TIM3+ LAG-3+ CD39+ T-BET+ | 1 |
| PD-1+ CD39+ EOMES+ CD44+ T-BET+ | 1 |
| PD-1+ LAG-3+ EOMES+ CD44+ T-BET+ | 1 |
| PD-1+ TOX1+ TIM3+ LAG-3+ CD39+ CD38+ TCF-1+ | 1 |
| TOX1+ TIM3+ LAG-3+ CD39+ EOMES+ CD44+ T-BET+ | 1 |
| PD-1+ EOMES+ CD38+ CD44+ | 1 |
| TOX1+ TIM3+ LAG-3+ CD39+ EOMES+ T-BET+ | 1 |
| TOX1+ TIM3+ LAG-3+ CD39+ EOMES+ CD38+ | 1 |
| PD-1+ EOMES+ CD38+ T-BET+ | 1 |
| TOX1+ CD39+ CD38+ CD44+ T-BET+ | 1 |
| TOX1+ TIM3+ LAG-3+ CD39+ TCF-1+ | 1 |
| PD-1+ TIM3+ LAG-3+ CD39+ CD38+ TCF-1+ | 1 |
| PD-1+ TIM3+ LAG-3+ CD39+ EOMES+ | 1 |
| TIM3+ CD39+ EOMES+ CD38+ TCF-1+ T-BET+ | 1 |
| TOX1+ TIM3+ CD39+ CD38+ TCF-1+ | 1 |
| PD-1+ TOX1+ TIM3+ EOMES+ CD38+ TCF-1+ T-BET+ | 1 |
| PD-1+ TOX1+ TIM3+ EOMES+ CD38+ TCF-1+ | 1 |
| PD-1+ TIM3+ CD38+ | 1 |
| PD-1+ TIM3+ CD38+ T-BET+ | 1 |
| PD-1+ TIM3+ CD38+ TCF-1+ | 1 |
| PD-1+ TIM3+ CD38+ TCF-1+ T-BET+ | 1 |
| PD-1+ TIM3+ CD38+ CD44+ | 1 |
| PD-1+ TIM3+ LAG-3+ CD38+ CD44+ TCF-1+ | 1 |
| PD-1+ TIM3+ LAG-3+ CD38+ CD44+ T-BET+ | 1 |
| PD-1+ TIM3+ LAG-3+ CD44+ T-BET+ | 1 |
| PD-1+ TIM3+ LAG-3+ CD44+ | 1 |
| PD-1+ TIM3+ LAG-3+ TCF-1+ | 1 |
| PD-1+ TIM3+ CD39+ T-BET+ | 1 |
| TIM3+ LAG-3+ CD39+ CD44+ T-BET+ | 1 |
| PD-1+ TIM3+ CD39+ TCF-1+ T-BET+ | 1 |
| TIM3+ LAG-3+ CD39+ CD44+ | 1 |
| TOX1+ LAG-3+ CD39+ EOMES+ CD38+ CD44+ T-BET+ | 1 |
| PD-1+ TIM3+ CD39+ CD44+ | 1 |
| LAG-3+ CD39+ CD44+ T-BET+ | 1 |
| PD-1+ TIM3+ CD39+ CD38+ | 1 |
| PD-1+ TIM3+ CD39+ CD38+ CD44+ T-BET+ | 1 |
| TIM3+ LAG-3+ EOMES+ CD38+ CD44+ | 1 |
| PD-1+ TOX1+ TIM3+ EOMES+ CD38+ CD44+ T-BET+ | 1 |
| PD-1+ TIM3+ LAG-3+ EOMES+ TCF-1+ | 1 |
| PD-1+ CD38+ CD44+ T-BET+ | 1 |
| TOX1+ TIM3+ CD39+ EOMES+ CD38+ | 1 |
| TOX1+ TIM3+ CD39+ EOMES+ CD38+ CD44+ TCF-1+ | 1 |
| PD-1+ TIM3+ LAG-3+ CD39+ CD38+ CD44+ TCF-1+ T-BET+ | 1 |
| PD-1+ TIM3+ LAG-3+ CD39+ CD38+ TCF-1+ T-BET+ | 1 |
| PD-1+ LAG-3+ CD39+ CD44+ | 1 |
| PD-1+ LAG-3+ CD39+ CD44+ T-BET+ | 1 |
| PD-1+ TIM3+ CD39+ EOMES+ TCF-1+ | 1 |
| PD-1+ LAG-3+ CD39+ CD38+ TCF-1+ | 1 |
| PD-1+ TIM3+ LAG-3+ CD39+ TCF-1+ T-BET+ | 1 |
| PD-1+ TOX1+ TIM3+ CD39+ EOMES+ CD38+ | 1 |
| PD-1+ LAG-3+ CD39+ CD38+ CD44+ | 1 |
| PD-1+ LAG-3+ CD39+ CD38+ CD44+ TCF-1+ | 1 |
| PD-1+ TIM3+ LAG-3+ EOMES+ CD44+ | 1 |
| LAG-3+ CD39+ EOMES+ CD44+ T-BET+ | 1 |
| TIM3+ CD39+ CD38+ CD44+ TCF-1+ | 1 |
| TOX1+ TIM3+ CD39+ EOMES+ CD44+ T-BET+ | 1 |
| TIM3+ LAG-3+ CD39+ CD38+ TCF-1+ | 1 |
| PD-1+ LAG-3+ CD39+ EOMES+ CD44+ T-BET+ | 1 |
| PD-1+ TOX1+ TIM3+ CD39+ CD38+ T-BET+ | 1 |
| PD-1+ TOX1+ TIM3+ CD39+ CD38+ | 1 |
| PD-1+ TOX1+ TIM3+ CD39+ CD44+ T-BET+ | 1 |
| PD-1+ TOX1+ TIM3+ CD39+ CD44+ | 1 |
| PD-1+ LAG-3+ CD39+ EOMES+ CD38+ TCF-1+ T-BET+ | 1 |
| TOX1+ TIM3+ LAG-3+ CD39+ CD44+ TCF-1+ T-BET+ | 1 |

**Supplementary Table S5.** Raw counts of cell-cell spatial interactions present in the dataset.

| Cell-Cell Spatial Interaction | Count |
| --- | --- |
| Ki-67- Neoplastic & Ki-67- Neoplastic | 2876770 |
| Myeloid cells & Myeloid cells | 1480053 |
| Myeloid cells & Ki-67- Neoplastic | 465886 |
| B cells & B cells | 450530 |
| Ki-67- Neoplastic & Ki-67+ Neoplastic | 437638 |
| CD44- CD4 Th & CD44- CD4 Th | 226250 |
| Mesenchymal & Mesenchymal | 220227 |
| CD44- CD4 Th & B cells | 219910 |
| CD44- CD8 T Other & CD44- CD4 Th | 206383 |
| Mesenchymal & Ki-67- Neoplastic | 202107 |
| Ki-67+ Neoplastic & Ki-67+ Neoplastic | 165702 |
| CD44- CD8 T Other & CD44- CD8 T Other | 158210 |
| CD44- CD8 T Other & B cells | 111164 |
| CD44- CD4 Th & Myeloid cells | 107677 |
| Myeloid cells & Mesenchymal | 90234 |
| CD44- CD8 T Other & Myeloid cells | 89976 |
| B cells & Myeloid cells | 62552 |
| Myeloid cells & Ki-67+ Neoplastic | 58793 |
| CD44- CD8 T Other & Ki-67- Neoplastic | 49618 |
| B cells & Ki-67- Neoplastic | 49336 |
| CD44- CD4 Th & Ki-67- Neoplastic | 48406 |
| CD44- CD4 Th1 Other & CD44- CD4 Th | 42570 |
| CD44- CD8 T Other & CD44- CD4 Th1 Other | 34844 |
| CD44- CD4 Th & mTREG | 32692 |
| CD44- CD4 Th & CD44+ CD4 Th | 31233 |
| CD44- CD4 Th & Mesenchymal | 29024 |
| CD44- CD8 T Other & mTREG | 27872 |
| CD44+ CD4 Th & B cells | 27606 |
| CD44- CD8 T Other & Mesenchymal | 26494 |
| CD44+ CD4 Th & CD44+ CD4 Th | 24718 |
| CD44+ CD4 Th & Myeloid cells | 24454 |
| CD44- CD4 Th1 Other & B cells | 24437 |
| CD44+ CD8 T Other & Myeloid cells | 23784 |
| mTREG & Myeloid cells | 23570 |
| CD44- CD8 T Other & CD44+ CD8 T Other | 23249 |
| Naive TREG & B cells | 23177 |
| CD44- CD4 Th & Naive TREG | 22405 |
| CD44+ CD8 T Other & CD44+ CD8 T Other | 22351 |
| Mesenchymal & Ki-67+ Neoplastic | 20910 |
| CD44+ CD8 T Other & CD44+ CD4 Th | 20823 |
| CD44+ CD8 T Other & CD44- CD4 Th | 20030 |
| CD44+ CD8 T Other & B cells | 19305 |
| CD44- CD8 T Other & Naive TREG | 16461 |
| B cells & Mesenchymal | 15010 |
| mTREG & B cells | 14656 |
| mTREG & Ki-67- Neoplastic | 14362 |
| CD44- CD8 T Other & CD44+ CD4 Th | 12734 |
| CD44- CD4 Th1 Other & CD44- CD4 Th1 Other | 11730 |
| mTREG & Mesenchymal | 10419 |
| CD44- CD4 Th1 Other & Myeloid cells | 9616 |
| mTREG & mTREG | 9087 |
| CD8 T NAIVE & CD44- CD8 T Other | 8303 |
| Naive TREG & Naive TREG | 7839 |
| CD8 T NAIVE & CD44- CD4 Th | 7103 |
| B cells & Ki-67+ Neoplastic | 6512 |
| CD44- CD4 Th & Ki-67+ Neoplastic | 6452 |
| CD44- CD8 T Other & Ki-67+ Neoplastic | 6392 |
| CD44- CD4 Th & T-BET+ TREG | 6102 |
| Naive TREG & Myeloid cells | 6035 |
| CD44+ CD4 Th & mTREG | 5918 |
| CD44- CD4 Th1 Other & Mesenchymal | 5711 |
| CD44- CD4 Th1 Other & Ki-67- Neoplastic | 5640 |
| CD8 TEMRA & CD44+ CD4 Th | 5367 |
| CD44- CD8 T Other & T-BET+ TREG | 5300 |
| CD44+ CD8 T Other & mTREG | 5186 |
| Naive TREG & Ki-67- Neoplastic | 5185 |
| CD8 T NAIVE & B cells | 5149 |
| T-BET+ TREG & B cells | 5086 |
| Naive TREG & mTREG | 4921 |
| CD44+ CD8 T Other & Ki-67- Neoplastic | 4863 |
| CD8 TEMRA & CD44+ CD8 T Other | 4772 |
| CD44+ CD4 Th & Mesenchymal | 4593 |
| CD44- CD4 Th1 Other & mTREG | 4305 |
| CD44+ CD4 Th & Naive TREG | 4166 |
| CD8 TEMRA & B cells | 3878 |
| CD44+ CD8 T Other & Mesenchymal | 3536 |
| CD44+ CD4 Th & Ki-67- Neoplastic | 3476 |
| CD8 TEMRA & CD44- CD8 T Other | 3454 |
| CD8 TEMRA & CD44- CD4 Th | 3399 |
| CD44- CD4 Th1 Other & T-BET+ TREG | 3382 |
| CD8 TEMRA & CD8 TEMRA | 3337 |
| CD8 TEMRA & Myeloid cells | 3226 |
| CD44- CD4 Th1 Other & CD44+ CD4 Th | 3112 |
| CD44- CD4 Th1 Other & Naive TREG | 3052 |
| CD44+ CD8 T Other & CD44- CD4 Th1 Other | 2766 |
| CD44+ CD8 T Other & Naive TREG | 2300 |
| CD4 Th1EMRA & CD44- CD4 Th | 1997 |
| CD4 Th1EM & CD44- CD4 Th | 1949 |
| mTREG & Ki-67+ Neoplastic | 1924 |
| T-BET+ TREG & Myeloid cells | 1857 |
| Naive TREG & Mesenchymal | 1852 |
| CD4 Th1EMRA & B cells | 1839 |
| Naive TREG & T-BET+ TREG | 1681 |
| CD4 Th1EM & Myeloid cells | 1667 |
| CD4 Th1EMRA & CD44+ CD4 Th | 1607 |
| CD4 Th1EM & CD44+ CD4 Th | 1542 |
| CD44- CD8 T Other & CD4 Th1EM | 1486 |
| mTREG & T-BET+ TREG | 1458 |
| CD8 TEM & CD44- CD8 T Other | 1322 |
| CD44+ CD8 T Other & CD4 Th1EM | 1311 |
| CD8 TEM & CD44+ CD8 T Other | 1285 |
| CD8 TEM & CD44+ CD4 Th | 1283 |
| CD8 T NAIVE & CD44- CD4 Th1 Other | 1238 |
| CD44- CD8 T Other & CD4 Th1EMRA | 1162 |
| CD8 TEM & CD44- CD4 Th | 1140 |
| T-BET+ TREG & T-BET+ TREG | 1119 |
| CD4 Th1EM & CD44- CD4 Th1 Other | 1112 |
| CD44+ CD4 Th & T-BET+ TREG | 1092 |
| CD8 TEX & CD44+ CD8 T Other | 1071 |
| CD44+ CD8 T Other & T-BET+ TREG | 1020 |
| T-BET+ TREG & Ki-67- Neoplastic | 1013 |
| T-BET+ TREG & Mesenchymal | 969 |
| CD44+ CD8 T Other & CD4 Th1EMRA | 969 |
| CD4 Th1EMRA & CD44- CD4 Th1 Other | 934 |
| CD4 Th1EMRA & Myeloid cells | 932 |
| CD8 TEMRA & Ki-67- Neoplastic | 878 |
| CD8 TEM & Myeloid cells | 847 |
| CD8 T NAIVE & CD44+ CD8 T Other | 837 |
| CD8 TEM & B cells | 808 |
| CD4 Th1EM & Mesenchymal | 790 |
| CD8 TEMRA & Naive TREG | 769 |
| CD8 TEX & CD44- CD8 T Other | 754 |
| CD8 T NAIVE & Myeloid cells | 740 |
| CD4 Th1EMRA & CD4 Th1EMRA | 739 |
| CD44- CD4 Th1 Other & Ki-67+ Neoplastic | 727 |
| CD44+ CD8 T Other & Ki-67+ Neoplastic | 710 |
| CD4 Th1EM & mTREG | 685 |
| CD8 T NAIVE & CD8 T NAIVE | 678 |
| CD8 TEM & CD8 TEM | 628 |
| CD8 T NAIVE & Naive TREG | 618 |
| CD4 Th1EM & B cells | 609 |
| CD44+ CD4 Th1 Other & Ki-67- Neoplastic | 580 |
| CD8 TEMRA & CD4 Th1EMRA | 576 |
| CD8 T NAIVE & mTREG | 574 |
| CD44+ CD4 Th1 Other & Myeloid cells | 550 |
| CD8 TEX & CD44+ CD4 Th | 522 |
| CD8 TEX & CD44- CD4 Th | 514 |
| CD4 Th1EM & Ki-67- Neoplastic | 510 |
| CD4 Th1EMRA & Naive TREG | 507 |
| CD44+ CD4 Th & Ki-67+ Neoplastic | 495 |
| CD8 TEM & CD8 TEMRA | 478 |
| CD44- CD4 Th1 Other & CD44+ CD4 Th1 Other | 477 |
| CD4 Th1EMRA & T-BET+ TREG | 476 |
| CD8 TEMRA & CD44- CD4 Th1 Other | 467 |
| CD8 TEMRA & mTREG | 459 |
| CD8 T NAIVE & CD44+ CD4 Th | 457 |
| Naive TREG & Ki-67+ Neoplastic | 451 |
| CD8 TEX & B cells | 442 |
| CD4 Th1EM & CD4 Th1EM | 431 |
| CD8 TEMRA & Mesenchymal | 383 |
| CD8 TEX & CD8 TEX | 368 |
| CD44+ CD4 Th1 Other & B cells | 353 |
| CD8 TEMRA & T-BET+ TREG | 350 |
| CD8 TEM & CD4 Th1EM | 340 |
| CD44+ CD4 Th1 Other & CD44+ CD4 Th1 Other | 338 |
| CD8 TEX & mTREG | 316 |
| CD8 TEX & Myeloid cells | 316 |
| CD8 TEM & mTREG | 309 |
| CD44+ CD4 Th1 Other & CD44- CD4 Th | 303 |
| CD4 Th1EMRA & CD44+ CD4 Th1 Other | 290 |
| CD8 TTEX & CD44+ CD8 T Other | 285 |
| CD4 Th1EM & T-BET+ TREG | 281 |
| CD44+ CD4 Th1 Other & Mesenchymal | 278 |
| CD8 TEMRA & CD8 TEX | 250 |
| CD4 Th1EMRA & Ki-67- Neoplastic | 247 |
| CD4 Th1EMRA & mTREG | 239 |
| CD8 TTEX & Myeloid cells | 221 |
| CD44- CD8 T Other & CD44+ CD4 Th1 Other | 215 |
| CD8 T NAIVE & T-BET+ TREG | 213 |
| CD44+ CD4 Th1 Other & CD44+ CD4 Th | 206 |
| CD8 TEM & CD44- CD4 Th1 Other | 199 |
| CD8 TEX & Ki-67- Neoplastic | 199 |
| CD8 TTEX & CD44- CD8 T Other | 194 |
| CD8 T NAIVE & Mesenchymal | 193 |
| CD44+ CD8 T Other & CD44+ CD4 Th1 Other | 186 |
| CD8 TTEX & CD44+ CD4 Th | 176 |
| CD8 TEMRA & CD4 Th1EM | 174 |
| CD4 Th1EMRA & Mesenchymal | 172 |
| CD8 TTEX & CD44- CD4 Th | 168 |
| CD8 TEM & T-BET+ TREG | 159 |
| T-BET+ TREG & Ki-67+ Neoplastic | 150 |
| CD8 T NAIVE & CD8 TEMRA | 150 |
| CD4 Th1EM & CD4 Th1EMRA | 146 |
| CD4 Th1EM & Naive TREG | 137 |
| CD8 TEM & CD4 Th1EMRA | 131 |
| CD8 T NAIVE & Ki-67- Neoplastic | 116 |
| CD8 TTEX & B cells | 111 |
| CD8 TEFF & CD44+ CD8 T Other | 108 |
| CD8 TEX & Naive TREG | 101 |
| CD8 TEX & CD44- CD4 Th1 Other | 96 |
| CD8 TEM & CD8 TEX | 91 |
| CD8 TEFF & B cells | 88 |
| CD8 TEM & Mesenchymal | 85 |
| CD8 TEX & T-BET+ TREG | 83 |
| CD8 TEX & CD8 TTEX | 83 |
| CD8 TEM & Naive TREG | 82 |
| CD8 TEMRA & Ki-67+ Neoplastic | 81 |
| CD8 TTEX & mTREG | 80 |
| CD8 TTEX & Ki-67- Neoplastic | 77 |
| CD8 TEX & Mesenchymal | 75 |
| CD8 TEX & CD4 Th1EM | 72 |
| CD8 TEM & Ki-67- Neoplastic | 68 |
| CD44+ CD4 Th1 Other & Ki-67+ Neoplastic | 65 |
| CD8 TEFF & CD44- CD8 T Other | 64 |
| CD44+ CD4 Th1 Other & T-BET+ TREG | 61 |
| CD8 TEFF & CD8 TEMRA | 56 |
| CD4 Th1EFF & B cells | 56 |
| CD44+ CD4 Th1 Other & mTREG | 54 |
| CD8 TEFF & Myeloid cells | 50 |
| CD8 TEFF & CD44- CD4 Th | 49 |
| CD4 Th1EFF & Myeloid cells | 47 |
| CD4 Th1EM & Ki-67+ Neoplastic | 45 |
| CD8 TEFF & CD8 TEFF | 42 |
| CD4 Th1EM & CD44+ CD4 Th1 Other | 41 |
| CD8 T NAIVE & CD4 Th1EMRA | 35 |
| CD8 TEMRA & CD8 TTEX | 34 |
| CD44+ CD4 Th1 Other & Naive TREG | 34 |
| CD8 TEMRA & CD44+ CD4 Th1 Other | 31 |
| CD8 TTEX & Mesenchymal | 30 |
| CD8 TEX & CD4 Th1EMRA | 29 |
| CD8 T NAIVE & CD4 Th1EM | 28 |
| CD4 Th1EFF & CD44+ CD4 Th | 28 |
| CD8 TEFF & CD44- CD4 Th1 Other | 26 |
| CD8 TEFF & Mesenchymal | 25 |
| CD8 TTEX & Naive TREG | 25 |
| CD8 TEFF & Ki-67- Neoplastic | 24 |
| CD4 Th1EMRA & Ki-67+ Neoplastic | 23 |
| CD8 T NAIVE & CD8 TEM | 22 |
| CD8 TEX & Ki-67+ Neoplastic | 22 |
| CD4 Th1EFF & CD44+ CD4 Th1 Other | 21 |
| CD8 TTEX & CD44- CD4 Th1 Other | 21 |
| CD8 T NAIVE & CD8 TEX | 20 |
| CD8 T NAIVE & CD44+ CD4 Th1 Other | 20 |
| CD4 Th1EFF & Ki-67- Neoplastic | 19 |
| CD4 Th1EFF & CD44- CD4 Th1 Other | 18 |
| CD8 TTEX & CD4 Th1EM | 18 |
| CD8 TTEX & CD8 TTEX | 17 |
| CD8 TTEX & Ki-67+ Neoplastic | 15 |
| CD4 Th1EFF & CD4 Th1EMRA | 15 |
| CD8 TEFF & CD44+ CD4 Th | 13 |
| CD8 TEM & CD8 TTEX | 11 |
| CD8 TEMRA & CD4 Th1EFF | 11 |
| CD8 TEX & CD44+ CD4 Th1 Other | 10 |
| CD44- CD8 T Other & CD4 Th1EFF | 8 |
| CD8 T NAIVE & Ki-67+ Neoplastic | 8 |
| CD4 Th1EFF & Ki-67+ Neoplastic | 8 |
| CD44+ CD8 T Other & CD4 Th1EFF | 7 |
| CD4 Th1EFF & CD44- CD4 Th | 7 |
| CD8 TTEX & CD4 Th1EMRA | 7 |
| CD8 TTEX & T-BET+ TREG | 7 |
| CD8 TEM & Ki-67+ Neoplastic | 5 |
| CD4 Th1EFF & Naive TREG | 5 |
| CD8 TEFF & CD8 TTEX | 5 |
| CD8 TEFF & T-BET+ TREG | 4 |
| CD8 TEFF & CD44+ CD4 Th1 Other | 4 |
| CD8 TEM & CD44+ CD4 Th1 Other | 4 |
| CD4 Th1EFF & CD4 Th1EM | 4 |
| CD8 TEX & CD4 Th1EFF | 4 |
| CD4 Th1EFF & Mesenchymal | 4 |
| CD8 TEFF & Naive TREG | 4 |
| CD8 T NAIVE & CD8 TEFF | 3 |
| CD4 Th1EFF & CD4 Th1EFF | 3 |
| CD4 Th1EFF & T-BET+ TREG | 3 |
| CD4 Th1EFF & mTREG | 2 |
| CD8 T NAIVE & CD8 TTEX | 2 |
| CD8 TEM & CD4 Th1EFF | 2 |
| CD8 TEFF & mTREG | 1 |
| CD8 TEFF & CD8 TEM | 1 |

## Notes

https://doi.org/10.5281/zenodo.8357193

https://github.com/kblise/PDAC_mIHC_paper

